# Synthetic Homoserine Lactone Sensors for Gram-Positive *Bacillus subtilis* using LuxR-type Regulators

**DOI:** 10.1101/2023.08.17.553781

**Authors:** Min Zeng, Biprodev Sarker, Nathaniel Howitz, Ishita Shah, Lauren B. Andrews

## Abstract

A universal biochemical signal for bacterial cell-cell communication could facilitate programming dynamic responses in diverse bacterial consortia. However, the classical quorum sensing paradigm is that gram-negative and gram-positive bacteria generally communicate via homoserine lactones (HSL) or oligopeptide molecular signals, respectively, to elicit population responses. Here, we create synthetic HSL sensors for gram-positive *Bacillus subtilis* 168 using allosteric LuxR-type regulators (RpaR, LuxR, RhlR, and CinR) and synthetic promoters. Promoters were combinatorially designed from different sequence elements (–35, –16, –10, and transcriptional start regions). We quantified the effects of these combinatorial promoters on sensor activity and determined how regulator expression affects its activation, achieving up to 293-fold activation. Using statistical design of experiments, we identified significant effects of promoter regions and pairwise interactions on sensor activity, which helped to understand the sequence-function relationships for synthetic promoter design. We present the first known set of functional HSL sensors (≥ 20-fold dynamic range) in *B. subtilis* for four different HSL chemical signals: *p*-coumaroyl-HSL, 3-oxohexanoyl-HSL, *n*-butyryl-HSL, and *n*-(3-hydroxytetradecanoyl)-HSL. This set of synthetic HSL sensors for a gram-positive bacterium can pave the way for designable interspecies communication within microbial consortia.

## Introduction

Synthetic microbial consortia have emerged as a powerful platform to utilize living systems for a variety of applications, ranging from biomanufacturing to bioremediation^1–6^. These collections of microbial cells can effectively distribute and specialize tasks among individual members, thereby reducing metabolic burden of each constituent microorganism^7,8^. Interaction between cells is required to elicit coordinated multicellular responses and can be achieved by cell-to-cell communication^5,9^. Bacterial intercellular communication can be implemented using direct DNA exchange by conjugation^10–12^ or molecular signals^13–19^, though each approach is typically limited to a subset of bacteria. Expanding the toolset for programmable intercellular communication could allow for a greater diversity of bacteria to be harnessed in engineered consortia.

Chemical signals from natural bacterial quorum sensing have been widely used in synthetic biology to engineer bacterial multicellular behaviors, and among them, diffusible homoserine lactone (HSL) signals have been used prevalently^13,17,20–31^. HSL signals are common in natural quorum sensing of gram-negative bacteria using LuxR family transcriptional regulators^18,32,33^. Most known examples have acylated HSL molecules, but others do not^34,35^. HSL quorum sensing has been well-studied and offers tremendous natural diversity of regulators and ligands^36–38^, making it a promising mode of bacterial intercellular communication for engineering consortia^39,40^. However, HSL sensors have been limited to use in gram-negative bacteria within engineered consortia. In nature, gram-positive bacteria are believed to commonly use oligopeptides for quorum sensing^18,32,33^, which require exporters for secretion and are sensed by two-component membrane-bound sensors^41^. While arguably more complicated to implement, peptide-based interspecies communication was demonstrated between gram-negative *Escherichia coli* and gram-positive *Bacillus megaterium*^42^. Other recent studies logically led us to question whether HSL sensing in gram-positive bacteria could be a viable alternative approach. Analyses of sequence conservation have shown that LuxR homologs are well-conserved among gram-positive bacteria^43^. Further, a gram-positive *Exiguobacterium* was reported to produce and sense a HSL^44^, and recently, a sensor using LuxR was engineered in gram-positive *Bacillus subtilis* for autoinduction^45,46^. These studies inspired us to study HSL-mediated signaling. Here, we investigate new HSL sensors engineered for *B. subtilis* and sequence-function relationships that govern their activity.

HSL transcription factor-based sensors typically contain a LuxR-type quorum sensing regulator (QSR) and a cognate quorum sensing promoter (P_QS_)^18,47,48^. For sensing, the regulator binds the HSL signal molecule and dimerizes to form the activated complex that then binds the DNA binding site in the P_QS_ to activate transcription by RNA polymerase^49,50^. To design a functional synthetic P_QS_ promoter, it must contain the DNA binding site for the regulator and other necessary promoter motifs. While the housekeeping sigma factors of *E. coli* (σ^70^) and *B. subtilis* (σ^A^) are homologous^51^ and share identical consensus hexamer –35 and –10 sequences^52,53^, promoters from gram-negative bacteria are usually not transferable to gram-positive bacteria^54–59^. The σ^A^ -dependent promoters in *B. subtilis* are known to differ from *E. coli* σ^70^-dependent promoters by commonly having a 4-bp motif known as the – 16 region (TRTG, where R = A or G), sometimes also referred to as the extended –10 region^56,60–62^. Far fewer *E. coli* promoters have a consensus sequence upstream of the –10 region (∼20% of natural promoters), and those that do have only a shorter 2-bp motif (NNTG)^63,64^. The promoter’s sequence downstream of the –10 region, called the transcription start region (TSR), has also been shown to significantly affect promoter activity in *B. subtilis*^65,66^. For a two-component light sensor in *B. subtilis*, its activity was improved by substituting a TSR from a strong promoter for *B. subtilis* and the consensus –10 motif into the original *E. coli* promoter sequence^67^. Very few studies have explored the generality of this strategy for re-designing sensor promoters from *E. coli* for *B. subtilis*.

Here, we develop HSL sensors in *B. subtilis* for four different HSL chemical signals using combinatorial design of synthetic promoters and tuning of regulator expression (Figure 1A). Sensors containing the quorum sensing promoters from *E. coli* had little or no activity in *B. subtilis*, yet by screening of sensor designs containing promoters comprised of different DNA sequences (–35, –16, –10, and TSR regions), we identified highly functional sensors for each HSL signal. In total, 99 sensor designs and 86 synthetic promoters were assayed in these experiments. All sensors were characterized in standard relative promoter units (RPU). This work resulted in the creation of a set of characterized HSL sensors for *B. subtilis* with at least 20-fold dynamic range and up to 293-fold induction. We systematically quantified the effects of the regulator’s expression level on sensor activity and identified that it is crucial for high activation. We also assayed a complete library of combinatorial promoters generated by statistical design of experiments (DOE) to quantify the effects of each promoter region and their interactions for one sensor (Figure 1B). Through this work we demonstrate the feasibility of creating HSL sensors and synthetic inducible promoters regulated by LuxR-type regulators in *B. subtilis*, which could be used to expand the use of HSL-mediated intercellular communication in bacteria.

**Figure 1.**
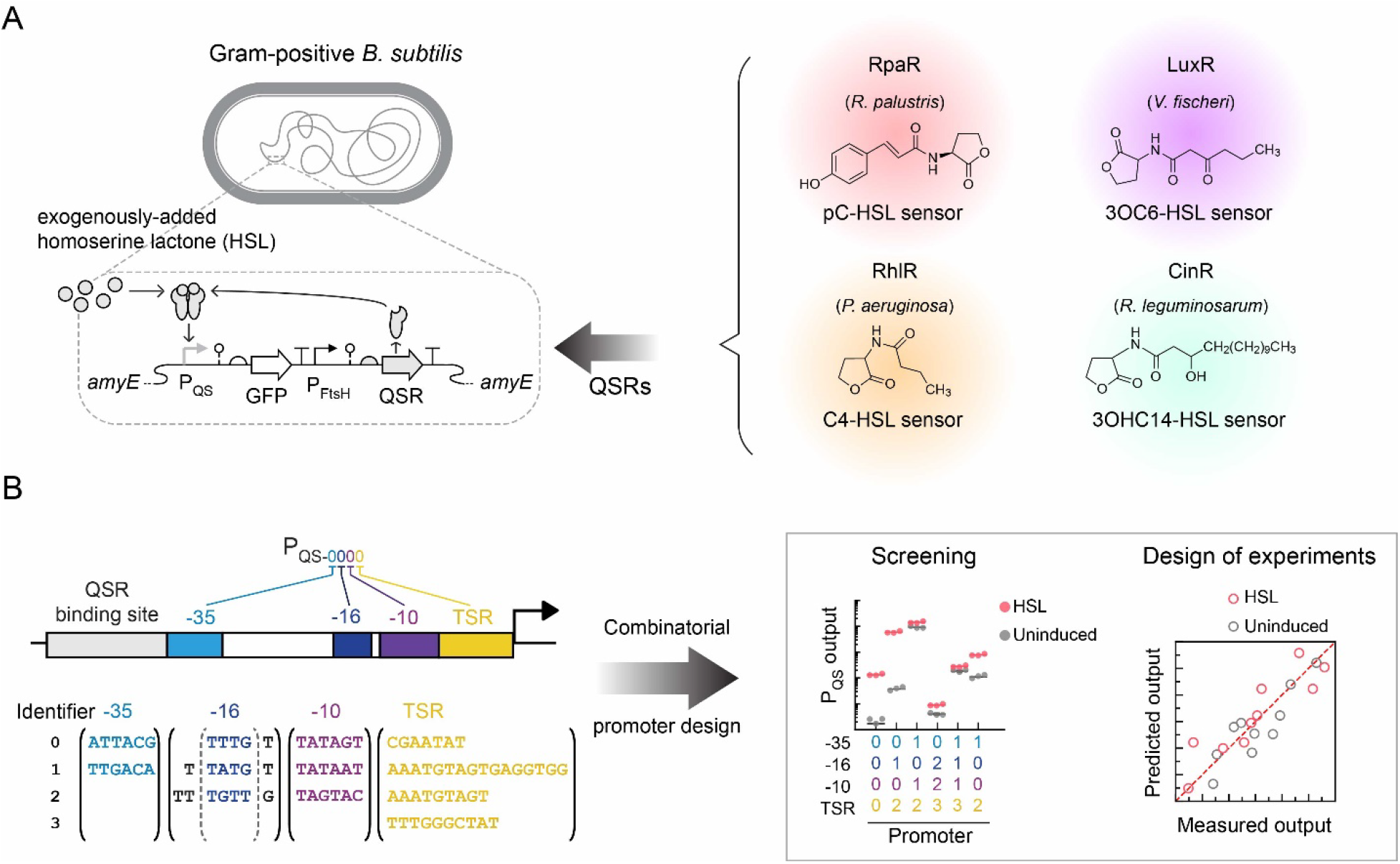
Homoserine lactone (HSL) sensor engineering in *B. subtilis*. **(A)** The engineered HSL sensors for gram-positive *B. subtilis* 168. The HSL sensors consist of a LuxR-type quorum sensing regulator (QSR) that binds the HSL ligand and the cognate quorum sensing promoter (P_QS_) that it regulates. Four LuxR-type regulators were selected from different gram-negative bacteria. **(B)** Synthetic P_QS_ promoters for *B. subtilis* are combinatorially designed using different DNA sequences for each promoter region: –35 region, –16 region, –10 region, and transcription start region (TSR). A sensor containing each promoter is constructed and screened. Design of experiments (DOE) further quantitatively studies the effect of each promoter region and their interactions on the activity of an HSL sensor.

## Results

### Genetic design of HSL biosensors for B. subtilis

First, we designed the basic genetic architecture to assay HSL sensors chromosomally integrated in the *B. subtilis* genome. In our design, we expressed the LuxR-type quorum sensing regulator (QSR) using a medium-strength constitutive promoter for *B. subtilis* (P_FtsH_)^68^. We included a hammerhead ribozyme (PlmJ^69^) downstream of the promoter for insulation and used a synthetic ribosome binding site (RBS) designed using the RBS Calculator^70–75^ to express the QSR in *B. subtilis*. A terminator (L3S2P15^76^), selected from a previously reported library of strong terminators, was placed after the QSR coding sequence. To be able to assay the promoter output of the sensor via cell fluorescence, the corresponding synthetic P_QS_ (i.e. sensor output promoter) expressed green fluorescent protein (GFP) using another ribozyme (RiboJ^69^), synthetic RBS (GFP-rbs), and terminator (L3S2P21^76^) (Figure 1A). The DNA construct for each HSL sensor was assembled in *E. coli* flanked by homology arms for *amyE* (552 and 1,002 bp in length) and subsequently transformed and integrated in the *B. subtilis* 168 genome (328,167 – 328,605 bp from *oriC*)^77^. To assay the sensor designs, we grew the cells with different concentrations of HSLs, measured the cell fluorescence via flow cytometry, and converted arbitrary fluorescence units to relative promoter units (RPU) (Methods).

Many bacterial studies have demonstrated and used an *in vivo* insulated relative promoter standard to facilitate reproducible measurements and quantitative comparisons between labs, cytometer instruments, and assay conditions (e.g. laser settings). Here, we adopted a similar strategy to convert arbitrary fluorescence units to relative promoter units (RPU) in *B. subtilis.* For our RPU standard, we used a medium-strength promoter (P_FtsH_) as the reference promoter in an insulated genetic construct containing the identical parts used to assay the sensor output (P_FtsH_, RiboJ, GFP-rbs, GFP, and L3S2P21), and the construct was also integrated at the *amyE* locus^77^ (Figure S1). Compared to a previously reported reference standard for *B. subtilis*^55^, we importantly included a self-cleaving ribozyme (RiboJ) to maintain an identical 5′ untranslated region of the mRNA transcript as we substituted different P_QS_ sequences, which reduces effects of the genetic context^67,69^ and allows us to measures the promoter activity by cell fluorescence. In the conversion to RPU, the activity of P_FtsH_ is equal to 1 RPU, and the autofluorescence of the wildtype cells is equal to 0 RPU. Of note, the conversion to RPU has also been shown to facilitate the interconnection of HSL sensors to logic gates in genetic circuits^78–81^, which could be useful in future work.

We selected four bacterial HSL quorum sensing systems for this study: (1) RpaR from *Rhodopseudomonas palustris* to sense *p*-coumaroyl-HSL (pC-HSL), (2) LuxR from *Aliivibrio fischeri* to sense 3-oxohexanoyl-HSL (3OC6-HSL), RhlR from *Pseudomonas aeruginosa* to sense *n*-butyryl-HSL (C4-HSL), and CinR from *Rhizobium leguminosarum* to sense *n*-(3-hydroxytetradecanoyl)-HSL (3OHC14-HSL). Each quorum sensing system senses a different HSL chemical species and has been widely used in *E. coli*^13,20–23,49,50,82,83^. Initially, we built and tested sensor constructs in *B. subtilis* containing the quorum sensing promoters used in *E. coli* for each system (P_Rpa-Ec_, P_Lux-Ec_, P_Rhl-Ec_, or P_Cin-Ec_, respectively) to determine the baseline activity for directly transferring the promoter DNA sequences. *B. subtilis* cells containing each sensor were assayed without induction and with addition of HSL at typical concentrations for maximal induction in *E. coli* (2 μM pC-HSL, 20 μM 3OC6-HSL, 3200 μM C4-HSL, or 20 μM 3OHC14-HSL, respectively). By performing this sensor characterization assay, we quantify three important attributes of each sensor design, which are the sensor OFF state output (i.e. uninduced basal promoter activity), sensor ON state activity (i.e. promoter activity when induced with HSL at high concentration), and the fold induction (the ratio of the ON to OFF state promoter activity), also known as the dynamic range of the sensor. Sensors containing the *E. coli* quorum sensing promoters for the Lux, Cin, and Rhl systems had low OFF state output (0.002 ± 0.001 RPU, 0.003 ± 0.001, 0.002 ± 0.001 RPU, respectively) but insignificant activation in *B. subtilis* with addition of the HSL signals (*p* > 0.05) (Figure S2). Only the sensor for the Rpa system containing the *E. coli* P_Rpa_ promoter (P_Rpa-Ec_) was significantly activated in *B. subtilis* (*p* < 0.05). Its sensor output (P_Rpa-Ec_) increased 82-fold on average with addition of pC-HSL. While this sensor showed a moderately large dynamic range, its activated promoter output was fairly low (0.13 ± 0.01 RPU), which would limit its use in downstream applications for control and actuation of gene expression. Overall, these results show poor transferability of gram-negative *E. coli* P_QS_ promoters to gram-positive *B. subtilis*, which was expected based on promoter sequence requirements that have been reported for *B. subtilis* (e.g. –16 region and TSR)^56–60,62^. Next, we re-designed each sensor for *B. subtilis*, aiming to improve the sensor’s activity and better understand the promoter requirements to activate each HSL sensor.

### Engineering the P_Rpa_ sensor output promoter for B. subtilis enhanced sensor activity

To investigate the relationship between the sensor’s promoter sequence and activity of the sensor in *B. subtilis*, we performed combinatorial engineering of the P_QS_ promoter and screening of the resulting sensor constructs in *B. subtilis*. We started with the pC-HSL sensor containing RpaR, given that activity was observed with the initial *E. coli* promoter sequence. By designing new P_Rpa_ output promoter sequences for the sensor construct (Figure 2A), we sought to increase the activated sensor output to at least 0.5 RPU, while maintaining a dynamic range of at least 20-fold. Based on prior studies reporting the importance of the -16 region and TSR in combination with the –10 and –35 sequences for σ^A^-dependent promoter activity in *B. subtilis*^56–60,62,84^, we defined four variable regions for the promoter sequences (–35 region, –16 region, –10 region, and TSR). For naming of the synthetic promoter sequences, we used a 4-digit number with each place value representing a promoter region (–35 region, –16 region, –10 region, and then TSR) and the numerical value representing a different DNA sequence (Figure 2B). With this nomenclature, we designated the sequence of each region in the original *E. coli* P_Rpa_ sequence as “0” and renamed the *E. coli* P_Rpa_ promoter accordingly (P_Rpa-Ec_ as P_Rpa-0000_). The DNA binding site for RpaR^34^ was maintained adjacent to and upstream of the –35 region (Figure 2B).

**Figure 2.**
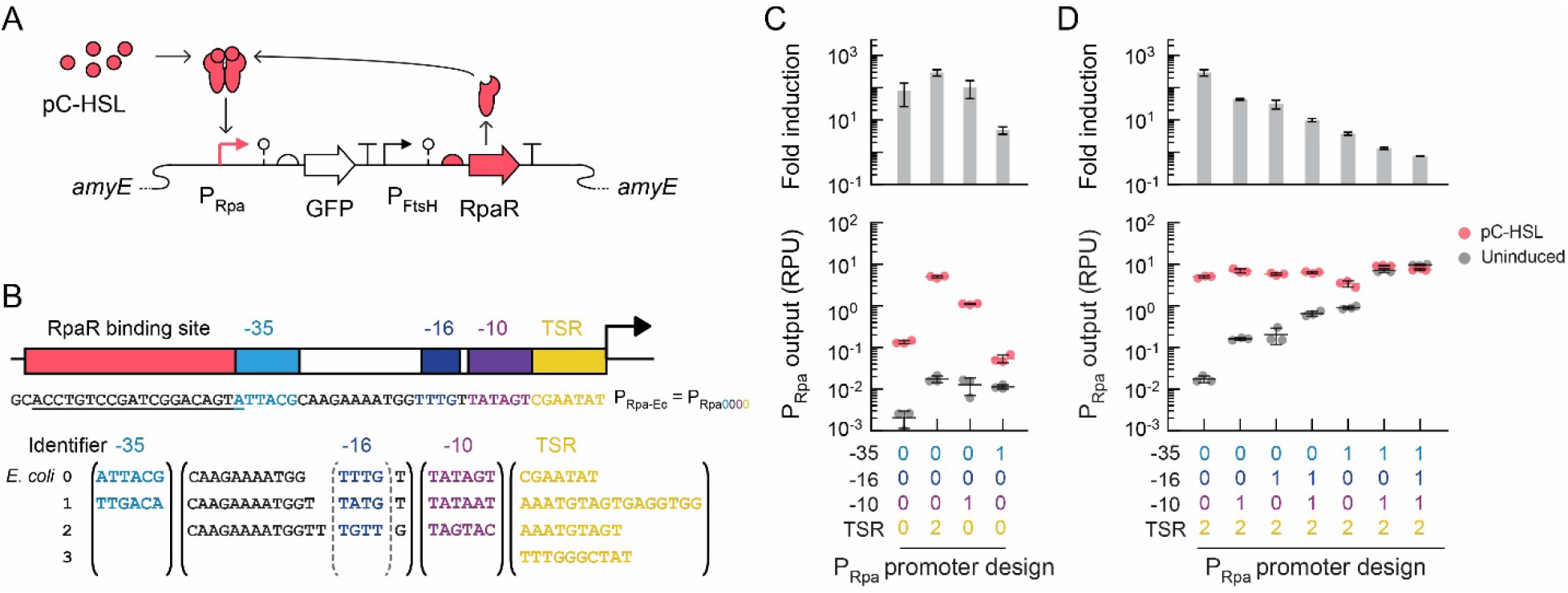
Combinatorial design of synthetic P_Rpa_ promoters created functional pC-HSL sensors. **(A)** The genetic illustration of the pC-HSL sensor. **(B)** DNA sequences for each promoter region were selected for P_Rpa_ promoter engineering. Identifier 0 represents sequences from *E. coli* P_Rpa_ promoter (P_Rpa-0000_). **(C)** pC-HSL sensors containing each P_Rpa_ promoter design were assayed without (gray circles) and with 2 µM pC-HSL (red circles). Cell fluorescence was measured by flow cytometry (≥ 10,000 cells per sample). The geometric mean fluorescence in arbitrary units was converted to relative promoter units (RPU) for the P_Rpa_ sensor promoter output (Methods). The bars represent the average of three independent experiments performed on different days. Error bars are the standard deviation.

We designed 14 synthetic P_Rpa_ promoters comprised of different combinations of DNA sequences for each region of the promoter. We selected DNA sequences from *B. subtilis* for each variable promoter region. The consensus sequences for σ^A^-dependent promoters were included for the –35 (TTGACA), –16 (TATG), and –10 (TATAAT) regions and designated as “1” (Figure 2B). Given that we aimed to increase the activated promoter output of P_Rpa_, we also chose DNA sequences based on two strong promoters in *B. subtilis* (P_SrfA_ and P_Veg_^45,67^). For the –16 region, we included the DNA sequence (TGTT, identified as “2”) from P_SrfA_ and P_Veg_. For the –10 region, we selected the sequence from P_SrfA_ (TAGTAC, identified as “2”). For the TSR, we chose DNA sequences from P_Veg,_ including a longer reported sequence^67^ (AAATGTAGTGAGGTGG, designated as “1”) and a shorter reported sequence^45^ (AAATGTAGT, designated as “2”), and the TSR sequence from P_SrfA_ (TTTGGGCTAT, designated as “3”). These DNA sequences for each promoter region were used to generate the combinatorial promoter library for the pC-HSL sensor. The sensors were constructed identically, except for the promoter sequence, and then assayed in *B. subtilis* with 2 μM pC-HSL and without induction.

Among the 15 sensor designs each containing a different P_Rpa_ promoter sequence, the sensor output ranged > 4,700-fold for both the OFF state (from 0.002 ± 0.001 RPU to 9.7 ± 0.3 RPU) and the ON state (from 0.0016 ± 0.0005 RPU to 9.2 ± 0.2 RPU), and the sensors’ average fold induction ranged from to 0.8 to 293-fold, showing vastly different sensor performance with changes to the promoter sequence (Figure S3). Notably, substitution of only one promoter region in the original *E. coli* P_Rpa_ promoter (P_Rpa-0000_) elicited large variations in the activity (Figure 2C). For example, replacement of the –35 region with the consensus sequence (P_Rpa-1000_) worsened sensor performance by both increasing the OFF state and decreasing the ON state (*p* < 0.05), resulting in only 4.8 ± 1.2-fold dynamic range. On the other hand, replacement of the TSR with 9 bp from P_Veg_ (P_Rpa-0002_) successfully increased (*p <* 0.05) the activated output to 5.0 ± 0.3 RPU, while only increasing the basal promoter activity to 0.017 ± 0.003 RPU. Indeed, this relatively small change in the promoter sequence yielded the best sensor in the library with 293 ± 65-fold dynamic range. Interestingly, substituting one or more consensus promoter elements into the P_Rpa-0002_ promoter deteriorated sensor performance by significantly increasing (*p* < 0.05) the uninduced promoter activity up to 9.6 ± 0.3 RPU (Figure 2D). All four promoters tested with both consensus –35 and –10 sequences (P_Rpa-1012_, P_Rpa-1112_, P_Rpa-1113_, and P_Rpa-1213_) exhibited a very small dynamic range (≤ 1.4-fold induction) (Figure S3). In total, 5 of the 14 sensors with new synthetic P_Rpa_ promoters met the specified criteria for a functional sensor (activated sensor output ≥ 0.5 RPU and dynamic range ≥ 20-fold). These functional sensors contained P_Rpa_ promoters with different promoter elements for the –16 region (sequences “0” and “1”), –10 region (sequences “0” and “1”), and TSR (sequences “0”, “1” and “2”), yet the results underscore the importance of the combination of promoter elements.

### Tuning expression of LuxR activated the synthetic 3OC6-HSL sensor in B. subtilis

For the 3OC6-HSL sensor containing LuxR, synthetic P_Lux_ promoters were designed based on combinations of promoter regions sequences from the P_Rpa_ library. We selected the 4 P_Rpa_ promoters from the pC-HSL sensors having the largest fold induction and substituted the RpaR binding site with the 20-bp palindromic LuxR binding site^85^ to generate the resulting set of P_Lux_ promoters (P_Lux-0002_, P_Lux-0011_, P_Lux-0012_, and P_Lux-0102_) (Figure 3A). In these initial sensor designs, a synthetic RBS for LuxR was designed using the RBS Calculator^70–75^ (predicted 9K au). We constructed the sensors and performed the sensor characterization assays (without and with 20 μM 3OC6-HSL). Insignificant activation by 3OC6-HSL (*p* = 0.19 – 0.93) was observed for all 4 of these sensor designs (Figure 3B). We hypothesized that an insufficient intracellular concentration of LuxR could be preventing activation^86^ and designed a synthetic RBS with > 5-fold higher strength for LuxR (50K au predicted strength^70–75^). Four sensors were constructed with the new RBS and other parts identical to the initial set. The sensor characterization assays were performed, and indeed, these four 3OC6-HSL sensors had significant activation (*p* = 0.0034 – 0.0035) and at least 4-fold induction (Figure 3B). The sensor having the highest dynamic range (20.2 ± 4.6-fold induction with P_Lux-0002_) has a lower ON state output (0.57 ± 0.05 RPU) than others, while the next closest (11.6 ± 0.3-fold induction with P_Lux-0012_) has a 4.6-fold greater ON state output (2.6 ± 0.3 RPU).

**Figure 3.**
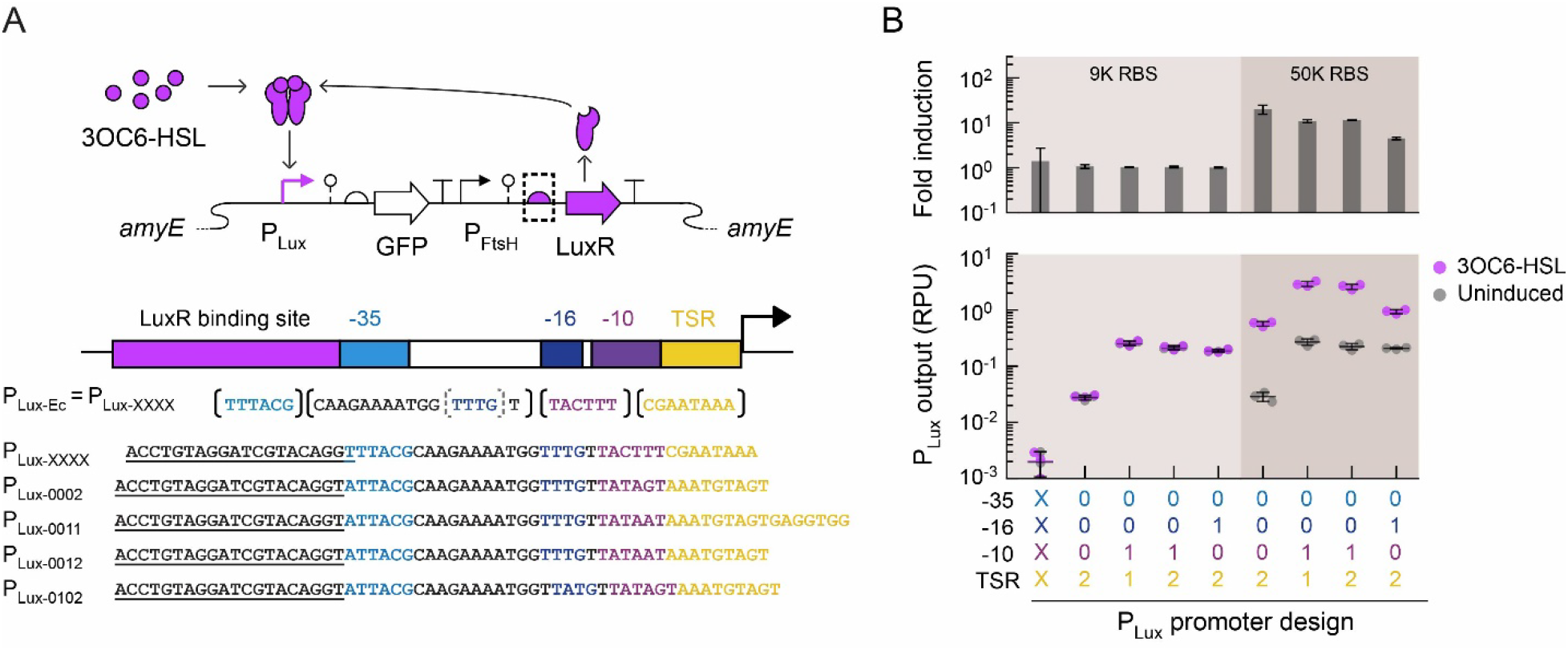
Tuning expression of LuxR activated the synthetic 3OC6-HSL sensor. **(A)** The genetic illustration of the 3OC6-HSL sensor with the P_Lux_ sensor output promoter. Four P_Lux_ promoters (P_Lux-0002_, P_Lux-0011_, P_Lux-0012_, and P_Lux-0102_) were designed by substituting the DNA binding site for LuxR into engineered P_Rpa_ promoters with different combinations of –35, –16, –10, and TSR sequences. Regions of the *E. coli* P_Lux_ promoter were designated as version “X” (P_Lux-Ec_ is identical to P_Lux_-_XXXX_). **(B)** 3OC6-HSL sensors containing the P_Lux_ promoters were assayed without (gray circles) and with 20 µM 3OC6-HSL (purple circles). Two different RBSs for LuxR expression were used (beige background colors). Cell fluorescence was measured by flow cytometry (≥ 10,000 cells per sample). The geometric mean fluorescence in arbitrary units was converted to relative promoter units (RPU) for the P_Lux_ sensor promoter output (Methods). The bars represent the average of three independent experiments performed on different days. Error bars are the standard deviation.

Next, additional sequences for the –35, and –10 regions were selected from prior studies to generate new synthetic P_Lux_ promoters to test in the sensor. Given that the – 35 and –10 consensus sequences of *E. coli* σ^70^ and *B. subtilis* σ^A^ are identical^64^, we selected six –35 or –10 regions for *E. coli* RNAP/σ^70^ that were shown to improve activation of inducible promoters in *E. coli*^87^. Similar studies of combinatorial promoter engineering in *B. subtilis* have not been reported to our knowledge. We substituted these six sequence elements into P_Lux-0002_ to create 6 new sensor designs (Figure S4). None of the sensors yielded an improved dynamic range, and only one design (with P_Lux-4012_) had significant activation (*p* < 0.05, 8.8 ± 0.5-fold induction) (Figure S4B). We also briefly tested if nucleotides outside the –16 region (4 bp) and between the –35 and –10 regions (sometimes referred to as the spacer sequence) affect sensor activity in *B. subtilis* as previously reported^88–91^. To do this, we substituted a reported weak spacer or strong spacer^89^ in place of the original 17-bp spacer in P_Lux-0102_. This failed to improve the sensor’s dynamic range (Figure S5). However, we observed large variation in the ON and OFF output among these sensors (19-fold and 19-fold, respectively) for these promoter permutations across all sensor designs tested. Out of the 19 sensors assayed, the sensor with the highest dynamic range (P_Lux-0002_) is notably identical to the P_Rpa_ promoter having the highest dynamic range (P_Rpa-0002_) except for the operator (Figure 3B).

### Activating the C4-HSL sensor by engineering synthetic P_Rhl_ promoters and RBS tuning

Based on the results from the previous two HSL sensing systems, we engineered the C4-HSL sensor containing RhlR by designing synthetic P_Rhl_ promoters in which we substituted the 20-bp RhlR binding site^92,93^ into four P_Rpa_ promoters from the pC-HSL sensors with the highest fold induction (resulting in P_Rhl-0002_, P_Rhl-0011_, P_Rhl-0012_, and P_Rhl-0102_) (Figure 4A). Two synthetic RBSs for RhlR were designed (8K and 800K au predicted strength^70–75^) for the sensors. We constructed each sensor design and performed the sensor characterization assays (3200 μM C4-HSL). All sensors with the weaker RBS (8K au) had a small dynamic range (≤ 1.2-fold) (Figure 4B). However, all 4 sensors with the stronger RBS (800K au) had significant activation (*p* < 0.05). Their induction spanned a large range (21.7 ± 3.5-fold induction with P_Rhl-0002_ to 1.8 ± 0.1-fold induction with P_Rhl-0102_), and the sensor displaying the largest dynamic range also had a moderate ON promoter output (0.36 ± 0.03 RPU with P_Rhl-0002_) (Figure 4B). Notably among these combinatorial promoter designs, we also observed that substitution of the –16 region in the sensor output promoter (P_Rhl-0002_ compared to P_Rhl-0102_) can elicit a drastic 12-fold decrease in the sensor’s dynamic range (21.7-fold induction to 1.8-fold induction). Two of the P_Rhl_ promoters were also tested in sensors with a medium-strength RBS (80K au predicted strength^70–75^), and both yielded insignificant activation (*p* > 0.05) (Figure S6).

**Figure 4.**
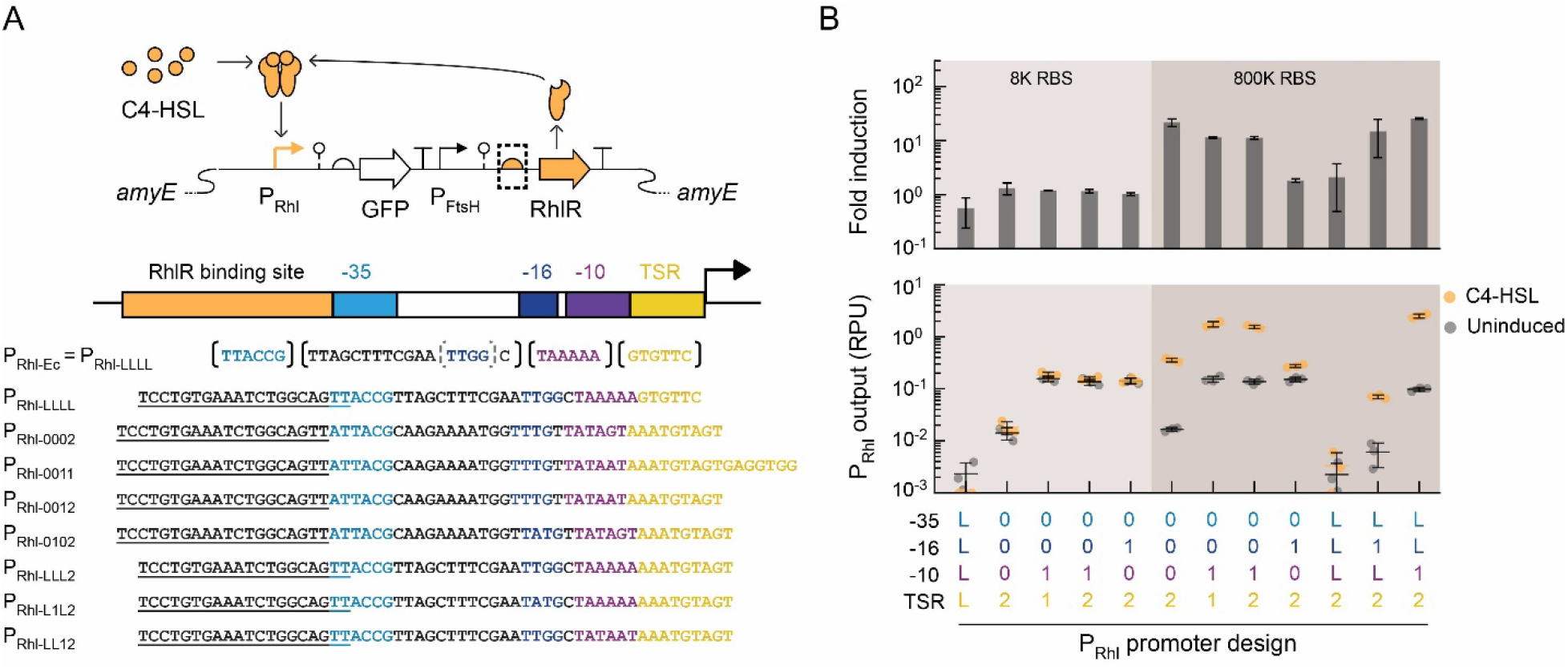
Engineering synthetic P_Rhl_ promoters for C4-HSL sensors in *B. subtilis*. **(A)** The genetic illustration of the C4-HSL sensor with the P_Rhl_ sensor output promoter. Four promoters (P_Rhl-0002_, P_Rhl-0011_, P_Rhl-0012_, and P_Rhl-0102_) were designed by substituting the DNA binding site for RhlR into P_Rpa_ promoters from top-performing sensors. Three more promoters (P_Rhl-LLL2_, P_Rhl-LL12_, and P_Rhl-L1L2)_ were designed by combinatorially substituting sequences from regions of the *E. coli* P_Rhl_ promoter, designated as the version “L” sequence (P_Rhl-Ec_ is identical to P_Rhl-LLLL_). **(B)** C4-HSL sensors containing different P_Rhl_ promoter designs were assayed without (gray circles) and with 3200 µM C4-HSL (orange circles). Two different RBSs for RhlR are shown (beige background colors). Cell fluorescence was measured by flow cytometry (≥ 10,000 cells per sample). The geometric mean fluorescence in arbitrary units was converted to relative promoter units (RPU) for the P_Rhl_ sensor promoter output (Methods). The bars represent the average of three independent experiments performed on different days. Error bars are the standard deviation.

We next sought to investigate the effect of substituting the –35, –16, and –10 sequences from the *E. coli* P_Rhl_ promoter (P_Rhl-Ec_) in this top-performing C4-HSL sensor promoter (P_Rhl-0002_). Therefore, we replaced these promoter regions, which were from the pC-HSL sensor’s *E. coli* P_Rpa_ promoter (identified as version “0”)^49^, with the corresponding sequences from P_Rhl-Ec_ and named each sequence version “L” (i.e. P_Rhl-Ec_ is identical to P_Rhl-LLLL,_ with “L” representing the last letter of Rhl)^49^. With this resulting synthetic promoter (P_Rhl-LLL2_), the sensor’s OFF output was very low (0.002 ± 0.001 RPU), yet insignificant induction was observed (*p* > 0.05), demonstrating that the TSR substitution alone was not sufficient to activate the *E. coli* P_Rhl_ promoter (Figure 4B). However, integrating the *B. subtilis* consensus –16 sequence into the promoter (P_Rhl-L1L2_) activated the sensor (14.8 ± 10.0-fold induction), albeit with a low ON state promoter output (0.07 ± 0.01 RPU) (Figure 4B). Surprisingly, instead substituting the – 10 region with *B. subtilis* consensus sequence (P_Rhl-LL12_) yielded a highly functional sensor with a large dynamic range (25.6 ± 0.9-fold induction), high ON output (2.5 ± 0.2 RPU), and low OFF output (0.2 ± 0.1 RPU) (Figure 4B). This further suggests that replacing promoter region sequences can be a useful strategy for engineering functional inducible promoters in *B. subtilis*.

### Engineering synthetic 3OHC14-HSL sensors by truncating the CinR operator

For the fourth HSL sensing system, the CinR regulator was used to construct a sensor for 3OHC14-HSL in *B. subtilis* (Figure 5A). We sought to design synthetic promoters regulated by CinR (P_Cin_) for the sensor. However, the sequence of the DNA binding site for CinR has not been explicitly identified, unlike for the other LuxR-type regulators. Some studies have utilized a long P_Cin_ promoter sequence (228-232 bp)^94,95^. Other studies in *E. coli* have used an 86-bp^21,96^ (designated version “A” here) or 19-bp^21,50,87,96^ CinR operator sequence (version “B” here), with neither being the canonical 20-bp palindromic sequence motif for LuxR-type regulator binding. In a separate study, a 29-bp DNA sequence (binding site version “C” here) was predicted to bind the CinR homodimer but was not experimentally tested^97^. In addition to these three putative CinR binding sequences, we combined versions “B” and “C” to design a fourth 43-bp sequence (version “D” here), which results in the latter half of version “A” (Figure 5B).

**Figure 5.**
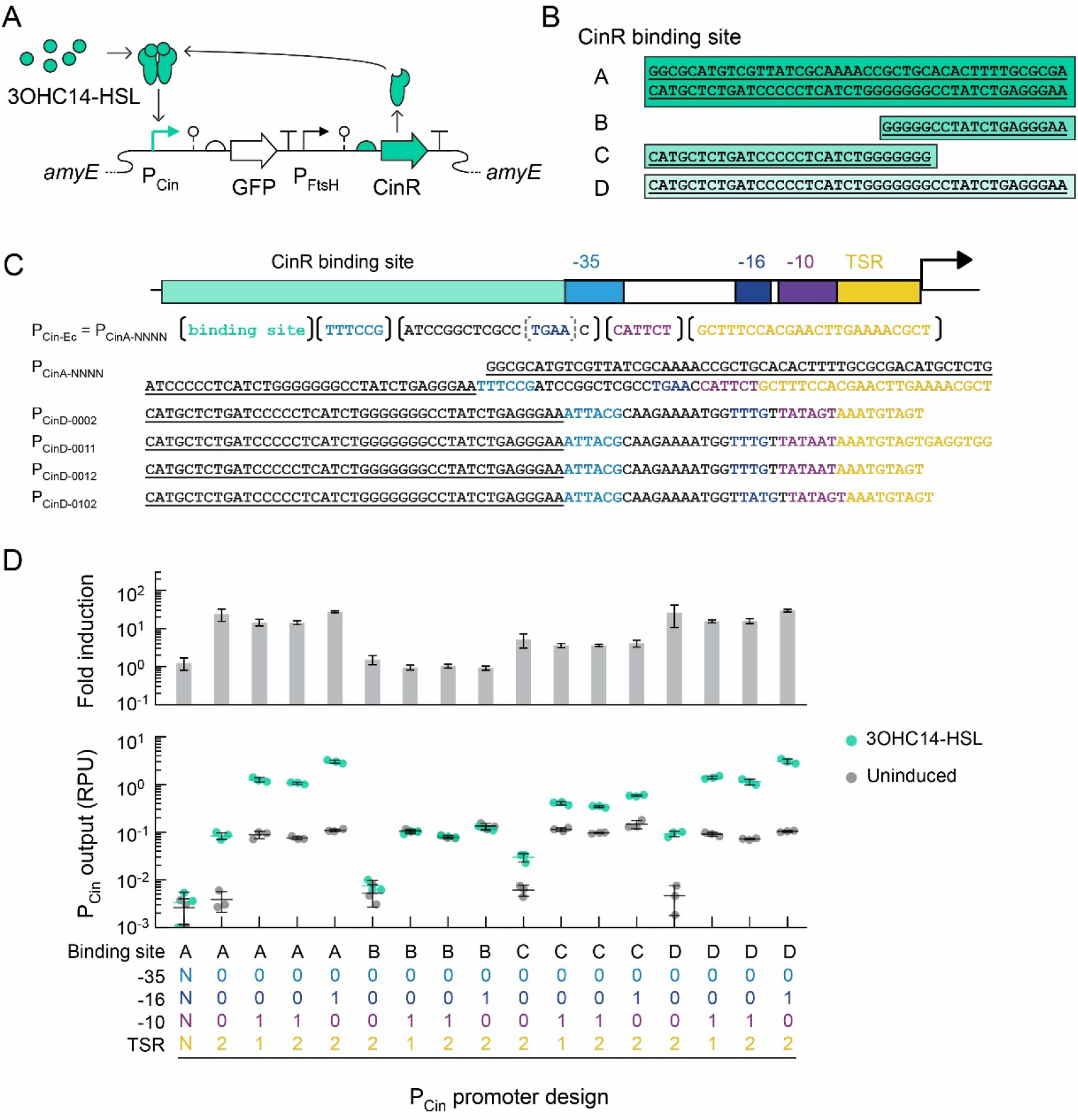
Engineering synthetic P_Cin_ promoters using different CinR binding sites. **(A)** Genetic schematic of the 3OHC14-HSL sensor. **(B)** Aligned DNA sequences of the four different CinR binding sites used. CinR binding sites A, B, C, and D have lengths of 86 bp, 19 bp, 29 bp and 43 bp, respectively. **(C)** Synthetic promoters were created with different combinations of binding site, –10, –16, –35, and TSR sequences. P_Cin_ promoter sequences for the subset containing the CinR binding site D are shown, and others are listed in Table S1. **(D)** 3OHC14-HSL sensors containing different P_Cin_ promoter designs were assayed without (gray circles) and with 20 µM 3OHC14-HSL (green circles). Cell fluorescence was measured by flow cytometry (≥ 10,000 cells per sample). The geometric mean fluorescence in arbitrary units was converted to relative promoter units (RPU) for the P_Cin_ sensor promoter output (Methods). The bars represent the average of three independent experiments performed on different days. Error bars are the standard deviation.

For each of these four binding site sequences, we designed four synthetic P_Cin_ promoters. We again selected 4 P_Rpa_ promoters from the pC-HSL sensors having the largest fold induction and substituted each CinR binding site in place of the RpaR binding site to generate a set of 16 synthetic P_Cin_ promoters. In the promoter naming, we included the identifier for the variable binding site sequence (e.g. each letter as in P_CinA-0002_ containing binding site “A”) (Figure 5C). These sixteen 3OHC14-HSL sensor designs were built and assayed in *B. subtilis*. CinR was constitutively expressed (P_FtsH_ and 15K au predicted RBS strength^70–75^) as in the other sensors. The basal promoter activity (uninduced) was comparable for each combination of –35, –16, –10, and TSR sequences independent of the binding site sequence (one-way ANOVA *p* > 0.05) with one exception (P_CinC-0012_). (Figure 5D). However, the binding site sequence largely affected the sensor’s ON state output and activation (Figure 5D). All sensors with binding site version “B” showed a very small dynamic range (1.1 ± 0.3-fold induction for 4 sensor designs), and all other 12 sensors showed significant activation (*p* < 0.05). The sensors containing the 86-bp long binding site “A” showed comparable activation to sensors containing the shortened 43-bp binding site “D” and identical –35, –16, –10, and TSR sequences (one-way ANOVA *p* > 0.05), which shows that the first half of the long binding site is unnecessary for activation by CinR. The four sensors containing the CinR binding site “C” exhibited lower ON state output (3-fold to 5-fold less) and a smaller dynamic range (4-fold to 7-fold less) compared to the corresponding sensors with binding site “D” (Figure 5D). Compared to other HSL sensors containing the same combination of –35, –16, –10, and TSR sequences, the induced output of P_CinD-0002_ was relatively low (0.09 ± 0.01 RPU). While the sensor with the P_CinD-0102_ promoter exhibited a comparable dynamic range (*p* > 0.05, 29.5 ± 2.3-fold induction), its activated promoter output was 33-fold greater (3.1 ± 0.4 RPU) and resulted in a highly functional 3OHC14-HSL sensor in *B. subtilis*.

### Characterizing the response functions for a set of HSL sensors in B. subtilis

In engineered microbial consortia that have HSL sensors for intercellular communication, the HSL signal can exist at a range of concentrations (e.g. temporally and/or spatially). The sensor response (i.e. sensor output) is dependent on the signal concentration. Knowing the quantitative relationship between the input signal and sensor output, which is called the sensor response function, facilitates designing a desired cellular response, such as integrating the sensors into designed genetic circuits^79–81,98–100^. Therefore, we next sought to experimentally determine the response functions for a set of the HSL sensors created for *B. subtilis* above.

We chose the design for each sensor with the greatest fold activation and a high induced output for response function characterization (HSL sensors containing P_Rpa-0002_, P_Rhl-LL12_, P_Lux-0002_, and P_CinD-0102_). The dose response for each selected HSL sensor was assayed using at least 12 concentrations of its cognate HSL. The experimental measurements were fitted to the Hill equation to determine the parameter values for each sensor response function (Methods, Table S2). The sensor output was assayed after incubation with the HSL inducer for 21 h, which is consistent with earlier sensor assays in this study (Figure 6). Additionally, we measured the response functions after a 5-h induction, which is commonly used for sensor characterization in *E. coli*^79,81^. The response functions for 5-h and 21-h induction were highly similar (Figure S7).

**Figure 6.**
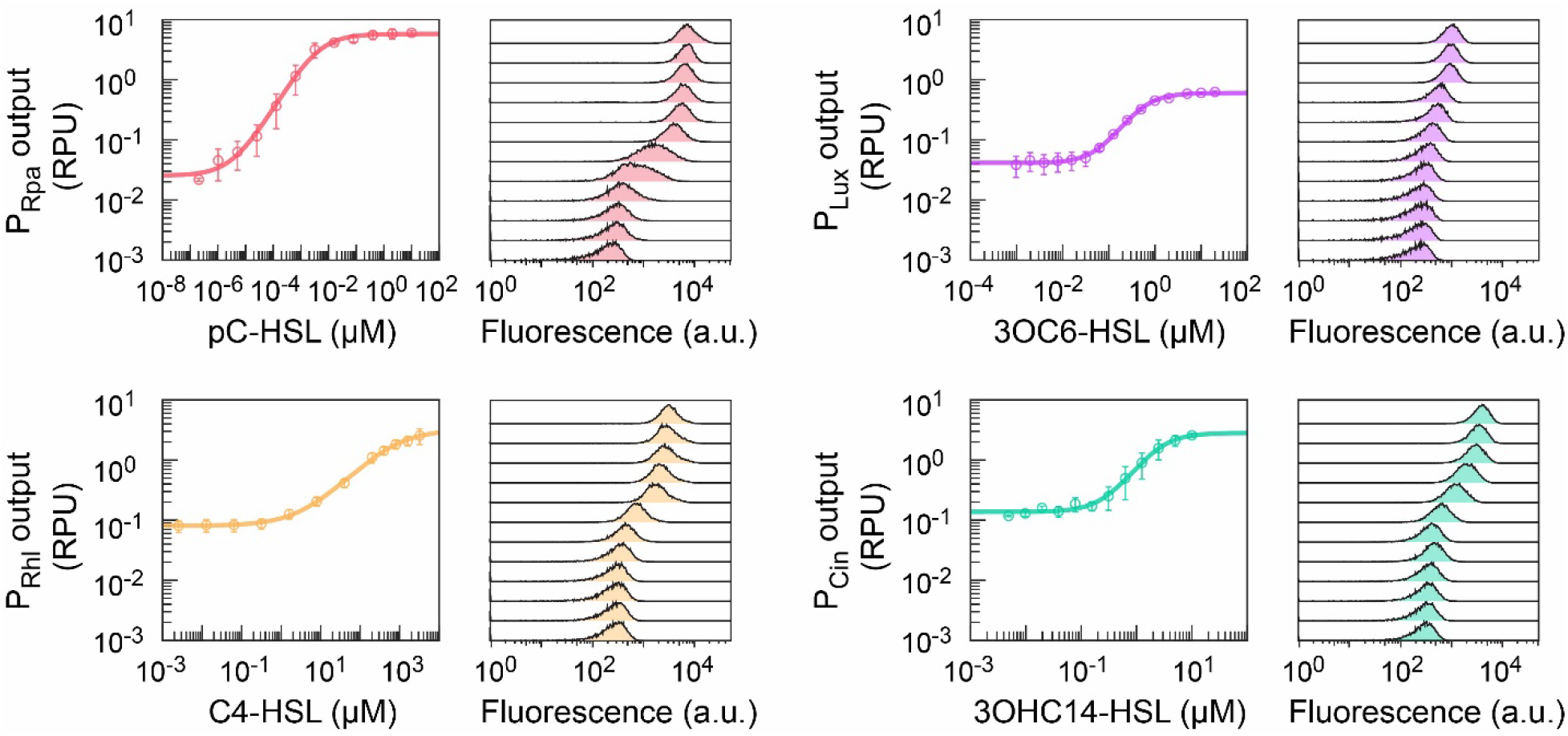
A set of synthetic HSL sensors with characterized response functions in *B. subtilis.* Sensor response functions were assayed for one sensor design for each HSL (greatest fold activation and high induced output). pC-HSL sensor containing P_Rpa-0002_ (red), 3OC6-HSL sensor containing P_Lux-0002_ (purple), C4-HSL sensor containing P_Rhl-LL12_ (orange), and 3OHC14-HSL sensor containing P_CinD-0102_ (green) are shown. The cell fluorescence was measured by flow cytometry (≥ 10,000 cells per sample). For each sensor, representative histograms spanning the concentration range are shown. The geometric mean of fluorescence in arbitrary units was converted to standard RPU, and the markers represent the average of three independent experiments performed on different days. Error bars are the standard deviation. The colored curves are the response functions fitted to the experimental data.

HSL sensors can be activated by noncognate HSLs, known as signal crosstalk, and this is well-known for quorum sensors with LuxR-family regulators in *E. coli*^49,50^. We measured the sensor output for the set of four HSL with each HSL signal at a high concentration (2 μM pC-HSL, 20 μM 3OC6-HSL, 3200 μM C4-HSL, and 20 μM 3OHC14-HSL). Signal crosstalk was observed for the C4-HSL sensor (RhlR) with 3OC6-HSL (46 ± 9% of cognate activation), pC-HSL sensor (RpaR) with C4-HSL (30 ± 13% of cognate activation), and 3OHC14-HSL sensor (CinR) with 3OC6-HSL (26 ± 12% of cognate activation) (Figure S8). The remaining nine pairs of sensors and non-cognate were orthogonal (<8 % of cognate activation for each) (Figure S8). Crosstalk for 3OC6-HSL with RhlR and C4-HSL with RpaR has been reported^49,50^. However, we also observed crosstalk in *B. subtilis* for 3OC6-HSL with CinR, which was not observed in *E. coli*^50^.

### Surveying the effects of regulator expression on HSL sensors’ response

To determine the effects of regulator expression on the response of these HSL sensors, we utilized a prototyping design that has inducible expression of the regulator in an insulated genetic construct, which is an approach we previously developed for engineering other transcription factor-based sensors^98^. We constructed four prototyping HSL sensor designs with inducible expression of each corresponding LuxR-type regulator using a xylose-inducible promoter (P_Xyl_) and the set of top-performing sensor promoter (P_Rpa-0002_, P_Rhl-LL12_, P_Lux-0002_, and P_CinD-0102_) (Figure 7A). We first characterized the response of the P_Xyl_ promoter so that we could determine the xylose inducer concentrations to span a 34-fold range for induction of QSR expression (Figure S9).

**Figure 7.**
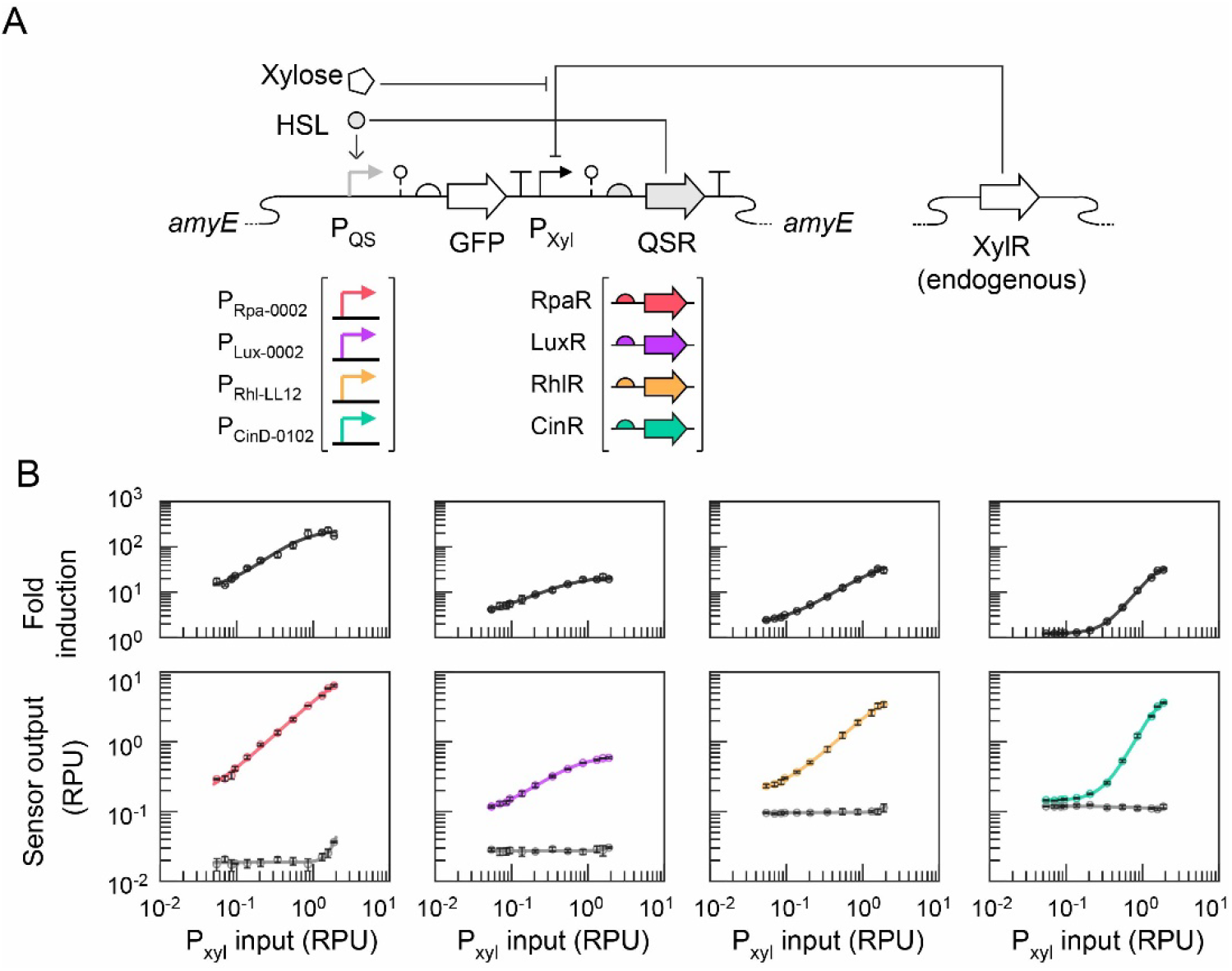
Surveying effects of regulator expression on HSL sensor activation in *B. subtilis*. **(A)** The genetic design to measure the effect of LuxR-type regulator expression on HSL sensor performance. A characterized xylose-inducible promoter (P_Xyl_) was used to express the QSR. **(B)** pC-HSL sensor (P_Rpa-0002_ and RpaR, red), 3OC6-HSL sensor (P_Lux-0002_ and LuxR, purple), C4-HSL sensor (P_Rhl-LL12_ and RhlR, orange), and 3OHC14-HSL sensor (P_CinD-0102_ and CinR, green) were assayed with 12 xylose concentrations to vary the P_Xyl_ input. The P_Xyl_ input value is from the assayed response function (Figure S9). The sensor output was measured without HSL (gray circle markers and line) and with the cognate HSL added (colored circle markers and line). The HSL concentrations used for each sensor were 2 µM pC-HSL, 20 µM 3OC6-HSL, 3200 µM C4-HSL, and 20 µM 3OHC14-HSL, respectively. The cell fluorescence was measured by flow cytometry (≥ 10,000 cells per sample). The geometric mean of fluorescence in arbitrary units was converted to standard RPU, and the markers represent the average of three independent experiments performed on different days. Error bars are the standard deviation.

We then assayed each HSL sensor in *B. subtilis* (without and with cognate HSL) for each specified concentration of xylose inducer (Figure 7B). The uninduced output (OFF state) of the sensors showed negligible change (≤ 1.2-fold) over the range of induction, except for the highest expression of RpaR (P_Xyl_ ≥ 1 RPU). However, the induced output (ON state) of each HSL sensor increased with increasing P_Xyl_ input, except for the 3OHC14-HSL sensor at very low induction (P_Xyl_ < 0.2 RPU) (Figure 7B). Across the P_Xyl_ input range (from 0.05 RPU up to 1.88 RPU), the dynamic range of each sensor varies significantly (*p* < 0.05) and by at least 5-fold for each sensor (pC-HSL sensor: 16-fold, 3OC6-HSL sensor: 5-fold, C4-HSL sensor: 13-fold, 3OHC14-HSL sensor: 26-fold). Our sensor designs assayed in earlier sections used the constitutive promoter P_FtsH_ (1 RPU). The results from these prototyping sensor designs suggest that the dynamic range of the 3OHC14-HSL and C4-HSL sensors could be significantly (*p* < 0.05) improved by increasing the expression of CinR and RhlR, respectively, using a stronger constitutive promoter (up to 1.88 RPU assayed here).

### Elucidating effects of P_Rpa_ promoter regions and their interactions on sensor activity

Lastly, we sought to quantitatively determine the effects of each promoter region (–35, –16, –10, and TSR) and their pairwise interactions on an HSL sensor’s activity. From the earlier sensor screening, we observed large differences in sensor activation for different combinations of promoter element sequences, and therefore, we aimed to understand the sequence-function relationships for designing the synthetic promoters. Synthetic hybrid promoters for yeast have been studied using design of experiments (DOE) approaches^101,102^, and the contribution of promoter regions (e.g. UP element, spacer, –10, –35, and 6-bp downstream of –10 region, operator sequence and operator location) to activity of promoters in *E. coli* has been investigated using large combinatorial libraries^61,103,104^. Here, we applied DOE and chose to investigate the pC-HSL sensor containing the RpaR regulator. We created a custom fractional factorial design including the pairwise interaction terms in the model (50% of all combinations of promoter element sequences) and built 36 P_Rpa_ promoters (Table S3). Each promoter was comprised of a different combination of DNA sequences for each region of the promoter (–35, –16, –10, and TSR). Of this set, six of the promoter designs were screened earlier and 30 are new P_Rpa_ promoters.

Each of the 36 sensors designs in the DOE library were constructed and assayed in *B. subtilis* (Figure 8A). Across the library, output spanned 8,300-fold for the ON state (0.002 ± 0.001 RPU – 15.0 ± 1.8 RPU) and 9,000-fold for the OFF state (0.0014 ± 0.0006 RPU – 12.6 ± 1.3 RPU). Among the sensor designs, the previously identified promoter (P_Rpa-0002_) remained to have the largest dynamic range (309 ± 64 fold induction). This dataset was used to build two linear regression models using least square regression for the basal sensor output (uninduced P_Rpa_ activity) and activated sensor output (P_Rpa_ activity with 2 μM pC-HSL), respectively. To reduce skewness of the data, the sensor output was logarithmically transformed (base 10). The variable promoter regions were input as categorical values for each sequence, which were identical to its one-digit identifier. We first evaluated linear models comprised of all 10 terms in each (4 individual effect and 6 pairwise interaction effect terms), which expectedly resulted in strong model agreement for the sensor OFF state (adjusted R^2^ = 0.95) and sensor ON state (adjusted R^2^ = 0.96) (Figure S10 and Table S4-S5). Without the six pairwise interaction terms, the model agreement worsened but were able to describe at least 77% of the observed variability of the sensor response (adjusted R^2^ = 0.77 for OFF state and R^2^ = 0.85 for ON state) (Figure S10 and Table S6-S7).

**Figure 8.**
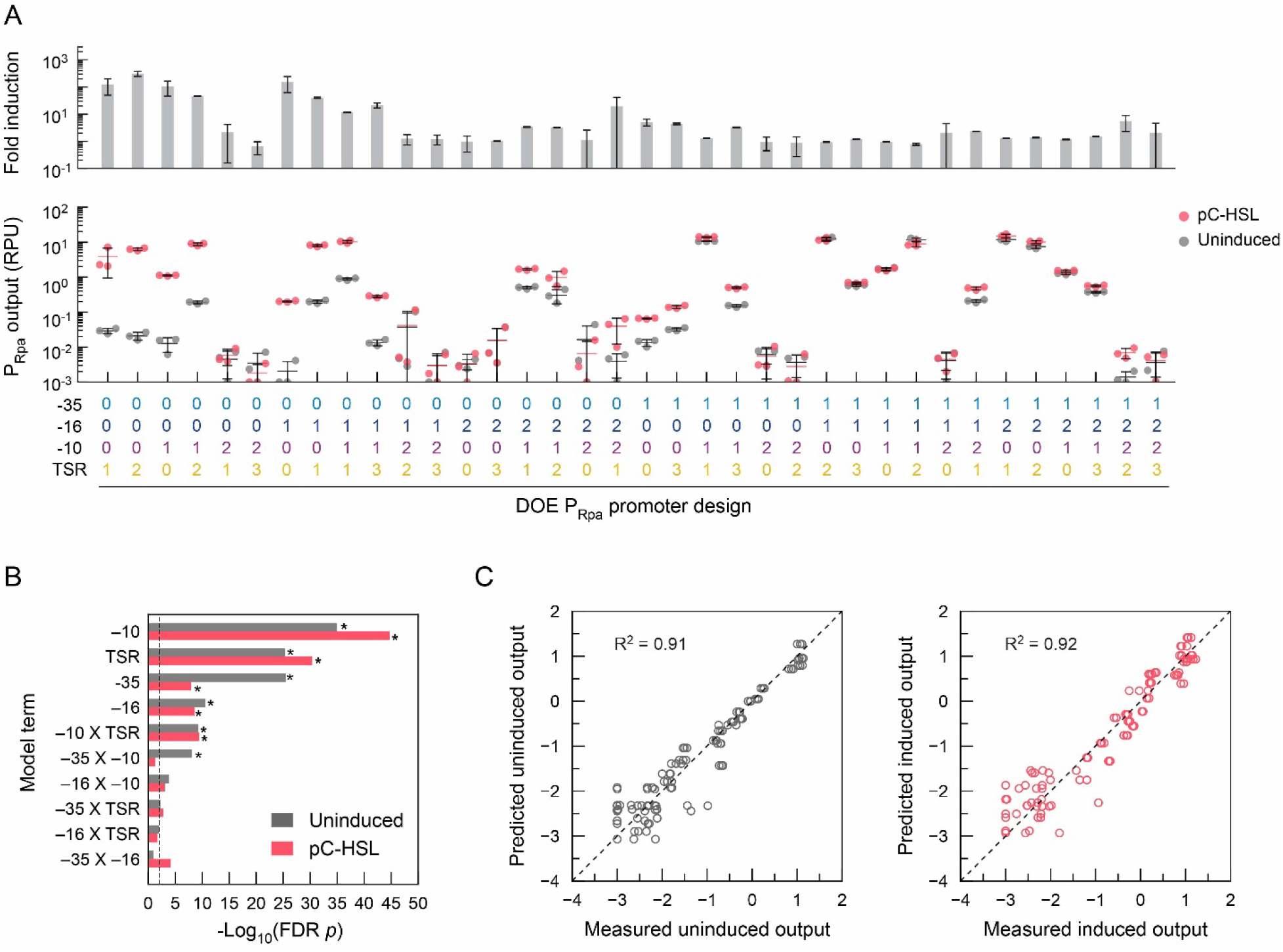
Combinatorial promoters by design of experiments (DOE) quantified effects of each promoter region and their pairwise interaction on pC-HSL sensor activity. **(A)** A DOE matrix for 36 combinatorial P_Rpa_ promoters with variable regions (two –35, three –16, three –10, and four TSR sequences) was created using the JMP software. Each pC-HSL sensor with one of the P_Rpa_ promoters was constructed and tested experimentally in *B. subtilis* (uninduced and 2 μM pC-HSL). Cell fluorescence was measured by flow cytometry (≥ 10,000 cells each sample). The geometric mean of fluorescence was converted to standard RPU (Methods). The output for each sensor is the average of three independent experiments performed on three different days. Error bars represent the standard deviation. **(B)** Linear regression models for the sensor activation as a function of the –10, –16, –35, and TSR regions (input as categorical variables) were generated in JMP using standard least square methods (Figure S10). The measured sensor output for the library of sensors (uninduced or with pC-HSL) were used to fit each model, and the false discovery rate (FDR) was evaluated. The significance of the effect for each 10 terms (4 individual effect and 6 pairwise interaction) was evaluated by calculating its FDR logworth, which is -log_10_(FDR *p-*value). Vertical line (dotted black) indicates *p*-value < 0.01. Asterisk (*) indicates highly significant terms (*p*-value < 0.00001). **(C)** The correlation between measured output for each sensor and the predicted output from the reduced linear regression models (Tables S7-S8). Only highly significant terms (*) were included in the models for the uninduced output (grey) and output with pC-HSL (red). The log_10_ transformed output in RPU is plotted.

We next assessed the significance of each individual and interaction effect of the promoter regions in the model by calculating the false discovery rate (FDR) probability using the Benjamini-Hochberg method^105^. A common criterion for significance in response models is FDR *p*-value < 0.01 (i.e. FDR logworth > 2). The –35, –16, –10, and TSR promoter regions all had a highly significant individual effect on the pC-HSL sensor OFF state activity and ON state activity (*p* < 0.00001), and the –10 region had the greatest significance for both (Figure 8B). In addition, we identified three highly significant (*p* < 0.00001) pairwise interactions between certain promoter regions for the pC-HSL sensor activity. The sensor OFF state was affected by interactions between the –10 with the –35 region and the –10 with the TSR region, while the sensor ON state was affected by interaction between the –10 and TSR regions (Figure 8B). Interaction between –35 and –10 regions were also reported to affect promoter activity in *E. coli*^106^. By constructing linear models with only these highly significant terms (*p* < 0.00001), strong agreement between the predicted and observed pC-HSL sensor OFF state (adjusted R^2^ = 0.91) and sensor ON state (adjusted R^2^ = 0.92) was found (Figure 8C and Table S8-S9). Using a less stringent cutoff for significance (*p* < 0.01), two additional interactions affect the OFF state output (–16 with –10 and –35 with TSR) and 3 additional interaction terms affect the ON state activity (–35 with –16, –16 with –10, and –35 with TSR). Given that we used discrete promoter regions for the input variables in the models, we could also identify the effects of specific variable sequences on sensor activity (Figure S11 and Table S10). This statistical study further supports the large effects individual promoter elements and their interactions can have on activation of a sensor in a gram-positive bacterium.

## Discussion

In this study, we created and assayed 86 synthetic P_QS_ promoters and 99 HSL sensor designs in total to develop the first set of sensors for 4 different HSLs in *B. subtilis*. Combinatorial design of promoters with different –10, –16, –35, and TSR sequences allowed us to assess the effects of these promoter regions on HSL sensor activity. We found that the effects of each region and combinations of sequences differ for the sensor OFF state (basal promoter activity) and sensor ON state (activated P_QS_ promoter output). Through quantitative analysis and statistical design, we determined the varying contribution of individual promoter regions and their pairwise interactions to HSL sensor activity, which could provide valuable insight for future sensor engineering in *B. subtilis*. This set of synthetic HSL sensors developed in this study has many potential applications for engineering interspecies bacterial communication and could enable the construction of tailored multispecies microbial communities with specific functions. While the gram-positive bacterium *B. subtilis* is generally not known to naturally utilize HSL-mediated quorum sensing^48,107,108^, we demonstrated that LuxR-type regulators from gram-negative bacteria can be used to create transcription factor-based biosensors for HSL molecules in *B. subtilis* via careful promoter engineering and tuning of regulator expression. By studying HSL sensors using four different LuxR-type regulators, we showed that this strategy is not limited to only LuxR itself in *B. subtilis*^46^, and this suggests the approach may be extended further and generalizable among the large family of LuxR-type regulators^47,109,110^.

The addition of HSL sensors and HSL-inducible promoters expands the genetic toolbox for tunable gene expression in *B. subtilis*, which is notable given the relative scarcity of inducible promoters in *B. subtilis* compared to *E. coli*^111^. The available inducible promoters for *B. subtilis* mostly utilize native regulators with sugar ligands^112,113^ or antibiotic-based inducible promoters^112,114^, along with a few other transcription factor-based^115,116^ or two-component system^67^ sensors. To complement strategies that have been developed to engineer inducible promoters in *B. subtilis*^115,117^, designing sensors based on the rules learned from this study could serve as another approach to translate transcription factor-based sensors from *E. coli* to *B. subtilis*.

Combinatorial promoter engineering has proven to be a useful strategy for designing new inducible promoters for different microrganisms^87,104,111,118–120^, and inspired by those studies, here we showed that similar approaches were effective to create functional HSL sensors in *B. subtilis.* The limits of these approaches in *B. subtilis* and gram-positive bacteria still need to be determined. We offer our experience with one other LuxR-type regulator as an example. While we constructed synthetic 3-oxododecanoyl-HSL sensors in *B. subtilis* using the LasR regulator from *Pseudomonas aeruginosa* and applying similar design strategies and promoter region sequences, only up to 3-fold induction was achieved by any sensor design (Figure S12). Therefore, there may be other factors and interactions that must be uncovered to extend this approach universally. As more design rules for synthetic inducible promoters are learned, it may become possible to utilize de novo promoter design strategies as have been demonstrated in *E. coli*^61,121^.

Our linear regression models for the P_Rpa_ sensor output confirmed that the –35 and –10 sequences alone are inadequate to explain the observed *B. subtilis* promoter activity^54^ and identified significant effects of the –16 region and TSR. Here, we identified the significant role of interactions between promoter regions affecting sensor activity using statistical design of experiments, and this suggests that statistical design could be a valuable approach to efficiently screen the combinatorial design space of similar promoters in *B. subtilis*. Other strategies, such as large libraries containing many promoter region sequences and regulator binding sites could also aid to systematically elucidate factors impacting sensor activity. Nevertheless, the combinatorial design strategies used in this work effectively developed highly functional sensors for four different HSL ligands in gram-positive *B. subtilis*. We believe these novel HSL sensors could have broad utility to allow diverse bacteria to speak the same biochemical language and establish new channels for interspecies communication in bacterial consortia.

## Materials and Methods

### Strains, Media, and Chemicals

*B. subtilis* 168 acquired from the Bacillus Genetic Stock Center was used to assay all sensors in this study. *E. coli* NEB 5-alpha (New England Biolabs) was used for construction and propagation of plasmids. *B. subtilis* 168 was made competent using SpC and SpII media. SpC media contains T-base (Sigma-Aldrich; 15 mM (NH_4_)_2_SO_4_, 80 mM K_2_HPO_4_, 44 mM KH_2_PO_4_, 3.4 mM Na_3_C_6_H_5_O_7_) supplemented with 1 mM MgSO_4_ (Sigma-Aldrich), 0.5% w/v glucose (Sigma-Aldrich), 0.2% w/v yeast extract (Sigma-Aldrich), 0.025% w/v casamino acids (Acros), and 20 µg/ml L-tryptophan (Sigma-Aldrich). SpII media contains T-base supplemented with 3.5 mM MgSO_4_, 0.5% w/v glucose, 0.1% w/v yeast extract, and 0.01% w/v casamino acids. M9 media used for sensor assaying contains M9 minimal salts (6.78 g/L Na_2_HPO_4_, 3.0 g/L KH_2_PO_4_, 1.0 g/L NH_4_Cl, 0.5 g/L NaCl) supplemented with 0.34 g/L thiamine hydrochloride (Sigma-Aldrich), 0.2% w/v casamino acids, 2 mM MgSO_4_, 0.4% w/v D-glucose, 0.1mM CaCl_2_ (Sigma-Aldrich), and was supplemented with 20 µg/ml L-tryptophan for *B. subtilis* 168, which is a tryptophan-requiring auxotroph. Luria Broth (LB) Miller media (Fisher Bioreagents) was used for cloning.

Homoserine lactone inducer chemicals were commercially purchased for *n*-butyryl-HSL (Sigma-Aldrich, #09945), 3-oxohexanoyl-HSL (Sigma-Aldrich, #K3007), *n*- (3-hydroxytetradecanoyl)-HSL (Sigma-Aldrich, #51481), and *p*-coumaroyl-HSL (Sigma-Aldrich, #07077). Antibiotics used for plasmid maintenance in *E. coli* were 50 µg/ml kanamycin (GoldBio) or 100 µg/ml carbenicillin (GoldBio). For selection after genome integration in *B. subtilis*, 5 µg/ml chloramphenicol (GoldBio) was used. HSL stock solutions were dissolved in pure DMSO (Fisher Bioreagents), and other chemicals were aqueous stock solutions.

### Plasmid Construction and DNA Assembly

HSL sensors were constructed on plasmids using hierarchical Type IIS DNA assembly in two sequential DNA assembly reactions. In the first DNA assembly reaction, each transcription unit (TU) construct (promoter, ribozyme, RBS, gene, and terminator) was assembled in one-pot Type IIS DNA assembly reaction using BsaI-HFv2 (New England Biolabs, NEB). Transcription units to assay the sensor output contained the P_QS_ promoter expressing the *gfp* gene. Other transcription units to express the QSR regulator for the sensor contained a promoter part (P_FtsH_ or P_Xyl_) to express the gene. For DNA assembly, P_FtsH_, P_Xyl_, and GFP were supplied as part plasmids in the reactions. P_QS_ promoter designs were supplied as oligo sandwiches (two annealed ssDNA oligos), and other genetic parts were supplied as PCR products. Each of these DNA assembly reactions contained: 10 fmol of each PCR product, 10 fmol of each part plasmid, and 50 fmol of an oligo sandwich, 5 fmol of an inverse PCR product of the plasmid backbone pAN202^79^, 5 U BsaI-HFv2 (NEB), 250 U T4 DNA ligase (2,000,000 U/ml, NEB) in 1X T4 ligase buffer (NEB) and the remainder nuclease-free water into the 5 µL total volume reaction. The DNA assembly reactions were incubated in a thermal cycler with heated lid with the following protocol: 37 °C for 5 h, followed by 30 °C for 30 min and deactivated at 80 °C for 20 min. For transformation, 2 µL of each TU assembly reaction was mixed with 5 µL chemically competent *E. coli* NEB 5-alpha (NEB) and otherwise followed the manufacturers recommended protocol. Transformations were plated on LB agar with carbenicillin. Each TU construct was sequenced by Sanger DNA sequencing (Azenta). In the subsequent DNA assembly reactions to build the whole sensor constructs, two TU constructs were assembled into a vector (pNH4) containing a kanamycin resistance cassette for plasmid selection in *E. coli*, two flanking *amyE* homology arms to direct *B. subtilis* genomic integration, and a chloramphenicol resistance cassette for *B. subtilis* to select for genomic integration. The sensor DNA assembly reactions contained: 10 fmol of each plasmid containing a transcription unit construct, 5 fmol of the pNH4 destination vector plasmid, 5 U BbsI (NEB), 250 U T4 DNA ligase in 1X T4 ligase buffer with the remainder nuclease-free water into a 5 µL total reaction. The DNA assembly reaction was then incubated in a thermal cycler with heated lid using the following protocol: alternating steps of 37 °C for 2 min and 16 °C for 2 min for 36 cycles, followed by 30 °C for 30 min and deactivated at 80 °C for 20 min. Transformation was performed by mixing 2 µL assembly reaction into 5 µL chemically competent *E. coli* NEB 5-alpha and plated on LB agar with kanamycin antibiotics. The insert length was confirmed by PCR. The constructed integration vectors containing sensors were amplified in overnight cultures *E. coli*, and the plasmid DNA was purified (Epoch Life Science). All genetic part sequences and plasmids used in this work are in Table S1 and S11, respectively.

### Genomic Integration in B. subtilis

To transform the constructed sensors into *B. subtilis*, first the *B. subtilis* 168 was made competent. To inoculate an overnight culture of *B. subtilis* 168, a colony was picked from a freshly streaked plate and transferred to 5 ml of LB media in a 14-ml snap cap culture tube, which was incubated at 37 °C and 250 rpm (Eppendorf Innova 44) for 16 h. The next morning, the overnight culture of *B. subtilis* was diluted to 0.01 OD_600_ in 12.5 ml SpC media in a 125 ml Erlenmeyer flask and incubated at 37 °C and 250 rpm. The OD_600_ of the culture was measured periodically to monitor cell growth. After 2 h in stationary phase, 2.5 ml of the *B. subtilis* culture was diluted into 25 ml prewarmed SpII media in a 250 ml Erlenmeyer flask and then incubated for 90 mins at 37 °C and 250 rpm. Next, cells were harvested by centrifugation at room temperature and 4,000 G (Eppendorf 5810R) in a 50 ml conical tube in a fixed angle rotor. The supernatant was decanted, and cells were gently resuspended in 2.5 ml supernatant, resulting in a 10-fold concentration of the cells. Cells were directly used for transformation or stored at - 80 °C in 10% v/v glycerol for later use. Prior to transformation, integration plasmids were linearized in a BsaI restriction digestion (BsaI-HFv2, NEB) at 37 °C for 2 h. About 1 µg of linearized DNA was added to 100 µl competent *B. subtilis* in a 14-ml snap cap culture tube and incubated at 250 rpm and 37 °C for 1 h. Then, 500 ml prewarmed LB was added into the culture and incubated at 250 rpm and 37 °C for 2 hr. After incubation, the culture was plated on LB agar with chloramphenicol. Colonies were streak purified and validated by PCR amplification of the *amyE* locus prior to assaying.

### Sensor Characterization Assays

For each sensor design, a single colony of *B. subtilis* 168 containing the constructed sensor was picked from a freshly streaked agar plate and was inoculated into 200 µl M9 with 5 µg/ml chloramphenicol in a well of a sterile 96-well U-bottom microtiter plate. Colonies for *B. subtilis* 168 wildtype and containing the RPU standard construct were similarly inoculated in the plate, but no antibiotic was added for wildtype cells. The plate was sealed with a breathable seal (AeraSeal, Sigma-Aldrich) and incubated at 37 °C, 1000 rpm for 16 h in a digital microplate shaking incubator (Elmi DTS-4). Cultures were diluted 178-fold by two serial dilutions, transferring 15 µl of each overnight culture into 185 µl fresh M9 with corresponding antibiotics in each dilution. The resulting plate containing the starter cultures was then incubated at 37 °C, 1000 rpm for 3 h. After 3 h of cell growth, the culture in each well was diluted 656-fold in two serial dilutions, first diluted 13.3-fold by transferring 15 µl culture to 185 µl fresh M9 with corresponding antibiotics. Then the cells were further diluted 49.3-fold into the final induction plate by transferring 4 µl cell culture into 193 µl M9 with antibiotic, HSLs, and inducers as appropriate. For ON/OFF sensor assays, cells were diluted into two wells, one having HSL added to the media and the other without HSL, and incubated at 37 °C and 1000 rpm for 21 h. The induction time was 21 h, except as otherwise noted for 5 h response function determination. Prior to flow cytometry, 1 µl for each culture was transferred to 199 µl PBS with 2 mg/ml kanamycin (to halt cell growth) and incubated for 30 min at room temperature.

The concentrations of the HSL inducers in the experiments are as follows. For ON/OFF sensor assays (21 h), either 2 µM pC-HSL, 20 µM 3OC6-HSL, 3200 µM C4-HSL, or 20 µM 3OHC14-HSL was used. For experiments assaying the signal crosstalk of the sensors, the concentrations of HSLs were 2 µM pC-HSL, 20 µM 3OC6-HSL, 3200 µM C4-HSL, and 20 µM 3OHC14-HSL.

### Flow Cytometry Analysis

Cell fluorescence was assayed using a BD Accuri C6 flow cytometer equipped with a 20 mW 480 nm solid state blue laser. At least 10,000 gated cell events were collected for each sample. Data were analyzed using FlowJo. A cell gate was created to distinguish *B. subtilis* 168 from electronic noise or debris. The geometric mean fluorescence of the cells was determined and then converted to relative promoter units (RPU). The conversion to RPU is as follows:

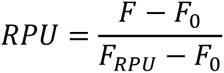

where *RPU* is the output of the sensor and was calculated using the fluorescence of the sample cells (*F*), autofluorescence of wildtype *B. subtilis* 168 cells (*F_0_*), and fluorescence of *B. subtilis* 168 cells harboring the RPU standard construct (*F_RPU_*). Samples with fluorescence equal to or below the autofluorescence of wildtype *B. subtilis* cells were set to the minimum limit of detection value of 0.001 RPU.

### Response Function Measurement and Fitting

The response functions of sensors were assayed under identical conditions as the sensor characterization assays, except additional inducer concentrations and induction times were assayed. Each selected sensor was assayed with an induction time of 5 h and 21 h by incubating the final inducer plate for 5 h or 21 h, respectively. The highest concentration of pC-HSL for 5 h and 21 h was 2 µM and 10 µM, respectively, and then was diluted by 5-fold for each subsequent concentration. The highest concentration of 3OC6-HSL for 5 h was 20 µM and then diluted by 2-fold for each subsequent concentration. The concentrations of 3OC6-HSL for 21 h were 20 µM, 10 µM, 5 µM, 2 µM and then diluted by 2-fold for each subsequent concentration. The concentrations of C4-HSL for both 5 h and 21 h were 3200 µM, 1600 µM, 800 µM, 400 µM, 200 µM, and then diluted by 5-fold for each subsequent concentration. The highest concentration of 3OHC14-HSL for 5 h and 2 h was 10 µM and then diluted by 2-fold for each subsequent concentration. The concentrations used for each sensor were selected with the aim of highly resolving the response function.

To determine response functions for selected sensor variants, the sensor outputs were fitted to the Hill equation^122^. The curve fit was performed in Python using the least squares method for the logarithmically transformed (base 10) output, minimizing the sum of the error between the fitted model and measured data points (https://www.scipy.org/citing-scipy/) using the equation below.

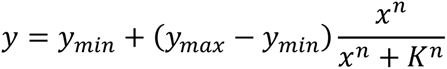

In this equation, the sensor output (*y*) is a function of the signal molecule concentration (*x*) and Hill parameters are *y_min_* (minimum output), *y_max_* (maximal output), *K* (Hill constant), and *n* (Hill coefficient).

### Sensor Assays with Inducible QSR Expression

The prototyping HSL sensor constructs that express the QSR by an inducible P_Xyl_ promoter were assayed identically as the sensor characterization assayed, except with the addition of the L-xylose inducer to the final inducer plate. The concentration of xylose used was 66 mM and then diluted by 2-fold for each subsequent concentration. For each xylose concentration, each HSL sensor was assayed without or with its cognate HSL. The concentration of each HSL used was: 2 µM pC-HSL, 20 µM 3OC6-HSL, 3200 µM C4-HSL, and 20 µM 3OHC14-HSL. The response function for the xylose sensor containing P_Xyl_ was experimentally determined as described above and used to determine the input promoter activity (Figure S9).

### DOE Library Design and Analysis

The JMP Pro 15 (SAS Institute) software was used to create the design of experiments (DOE) matrix and analyze the data, including model generation and false discovery rate (FDR) calculations. The fractional factorial DOE matrix for the pC-HSL sensor was created using the default settings for custom DOE design by specifying four categorical factors (–35, –16, –10 and TSR). Each categorical factors have different levels corresponding to the number of identifiers of each promoter region (Figure 2B), which are 2 levels for –35 (“0” and “1”), three for –16 (“0”, “1” and “2”), three for –10 (“0”, “1” and “2”) and four for TSR (“0”, “1”, “2” and “3”). For the model fitting of the response, we included one output of each model, and generated separate models for the ON state sensor output and OFF state sensor output. In the model options, we included the main effects terms for each individual promoter region (–35, –16, –10, and TSR regions) and their second-order (i.e. pairwise) interactions (6 terms), which resulted in a matrix of 36 combinatorial promoter designs (Table S2). The set of pC-HSL sensors containing each P_Rpa_ promoter design were experimentally tested using the sensor characterization assay as described above. Using the standard least squares method in JMP Pro, linear regression models for the log_10_(uninduced output) and log_10_(induced output) were constructed. For 10 model effects (4 promoter regions and their 6 pairwise interactions), we constructed full linear regression models. The linear regression models without any pairwise interaction effects were also constructed. The effect summary for each full linear regression model was generated by JMP Pro. We constructed the reduced linear regression models by removing insignificant effects (Figure 8B, false discovery rate *p* > 0.00001) for each linear regression model.

### Statistical Analysis

The uninduced and induced outputs of a sensor design was compared using a paired, two-tailed Student’s *t*-test. Different promoter designs and sensor designs were compared using independent, two-tailed Student’s *t*-tests. Student’s *t*-tests and ANOVA analysis were performed using the Scipy package of Python (https://www.scipy.org/citing-scipy/).

## Author Contributions

LBA, MZ, and BS conceived of the study and designed experiments. MZ, BS, NH, and IS performed the experiments. MZ, BS, and LBA analyzed data. MZ, BS, and LBA wrote the manuscript with support from all authors. LBA directed the research and acquired funding.

## Acknowledgments

This work was supported by funds from the National Science Foundation under Grants No. CBET-1943695 and MCB-2211039 to LBA. Additional funding was provided by start-up funds from the University of Massachusetts Amherst and funding from the Marvin and Eva Schlanger faculty fellowship to LBA. MZ received support from a Douglas fellowship. We thank the Andrews group members for their feedback on experimental design and manuscript preparation.

