## Supporting Information for "Synthetic Homoserine Lactone Sensors for Gram-Positive *Bacillus subtilis* using LuxR-type Regulators"

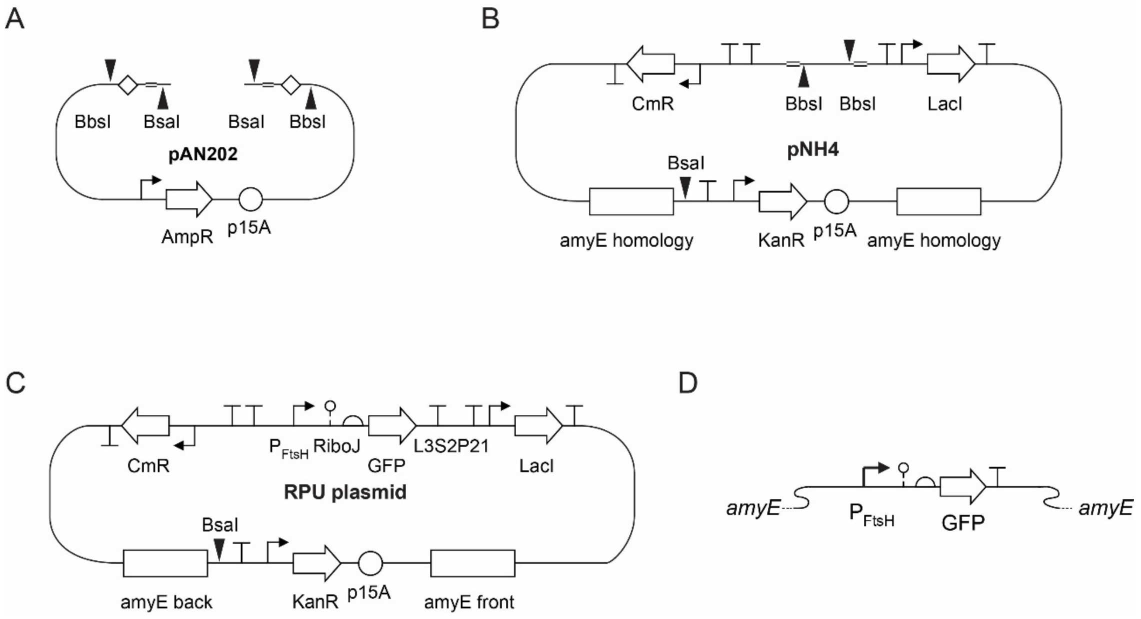

**Figure S1. Plasmids map used in this study. (A)** The plasmid map of pAN202 for the construction of transcription units. Diamonds represent linker sequence for TypeIIS assembly using BbsI. **(B)** The shuttle vector pNH4. The kanamycin resistance (KanR) is used for plasmid selection in *E. coli*. The chloramphenicol resistance (CmR) is used for genome integration selection in *B. subtilis*. The BsaI is used for linearization of plasmids in *B. subtilis* transformation. **(C)** The relative promoter units (RPU) construct on the integration plasmid contains a constitutive promoter $P_{FtsH}$ and characterization cassette (RiboJ, rbs-GFP, and GFP). **(D)** RPU reference standard after integration into the *B. subtilis* genome. The chloramphenicol resistance cassette and LacI cassette are not shown.

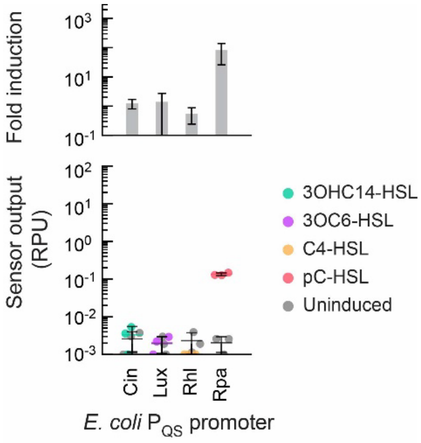

**Figure S2. Characterization of HSL sensors containing *E. coli* quorum sensing (QS) promoters in *B. subtilis*.** The activity of quorum sensor designs with *E. coli* QS promoters ($P_{Cin\text{-}Ec}$, $P_{Lux\text{-}Ec}$, $P_{Rhl\text{-}Ec}$, and $P_{Rpa\text{-}Ec}$) were assayed in *B. subtilis*. The sensor constructs from the corresponding integration plasmids (pMZ1000, pMZ2000, pMZ3000, and pMZ4000) were chromosomally integrated into *B. subtilis* at the *amyE* locus. The sensor output was measured using a fluorescent reporter. Sensor characterization assays were performed for *B. subtilis* cells harboring each sensor without inducer (gray markers) or with the corresponding cognate HSL exogenously added (colored circles). Cell fluorescence was measured via flow cytometry and converted into relative promoter units (RPU) (Methods). Markers represent the geometric mean output of a population of at least 10,000 cells. Assays were performed in identical experiments performed on three separate days. The concentrations used for each HSL species were: 20 μM 3OHC14-HSL, 20 μM 3OC6-HSL, 3200 μM C4-HSL, and 2 μM pC-HSL, respectively. For each experiment, the fold induction was calculated as the ratio of the sensor output for the induced to uninduced samples. Bars represent the average fold induction. All error bars represent one standard deviation.

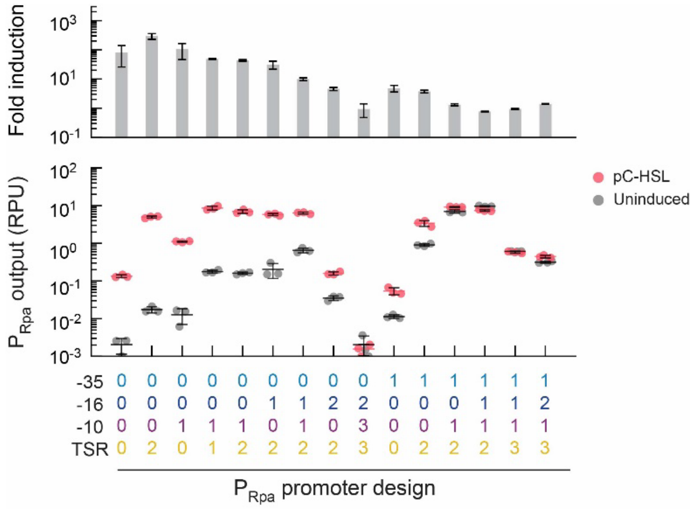

**Figure S3. Combinatorial design of $P_{Rpa}$ promoters created functional pC-HSL sensors.** The pC-HSL sensor containing synthetic $P_{Rpa}$ promoters were assayed without pC-HSL (gray cycles) or with 2 μM pC-HSL (colored cycles). Markers represent the geometric mean output of a population of at least 10,000 cells. Assays were performed in identical experiments performed on three separate days. For each experiment, the fold induction was calculated as the ratio of the sensor output for the induced to uninduced samples. Bars represent the average fold induction. All error bars represent one standard deviation.

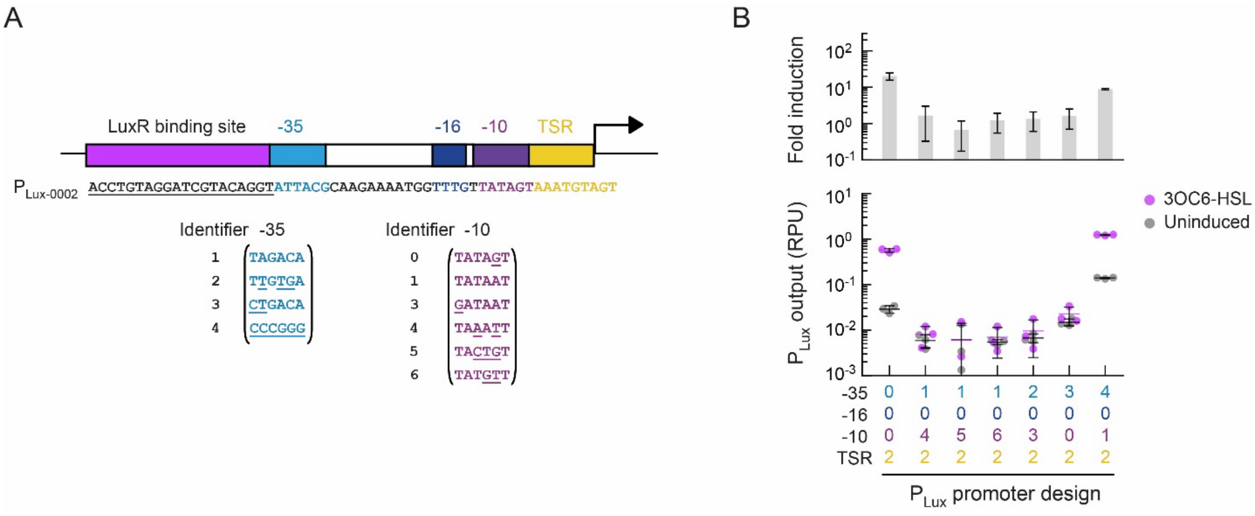

**Figure S4. Tuning synthetic $P_{Lux-0002}$ promoter by combinations of –35 regions and –10 regions. (A)** The –35 region and –10 region in $P_{Lux-0002}$ were substituted with selected 6 combinations, resulting in $P_{Lux-1042}$, $P_{Lux-1052}$, $P_{Lux-1062}$, $P_{Lux-2032}$, $P_{Lux-3002}$, and $P_{Lux-4012}$, respectively. Those combinations had been shown to improve the dynamic range of sensors with transcription activators in *E. coli*[1]. **(B)** The 3OC6-HSL sensors were assayed in *B. subtilis* without (gray cycles) or with 20 µM 3OC6-HSL (colored cycles). The bars represent the average of three independent measurements. Error bars represent the standard deviation. Sensors with the 50K RBS on integration plasmids (pMZ2013 – pMZ2018) were integrated into *B. subtilis amyE* locus before assaying.

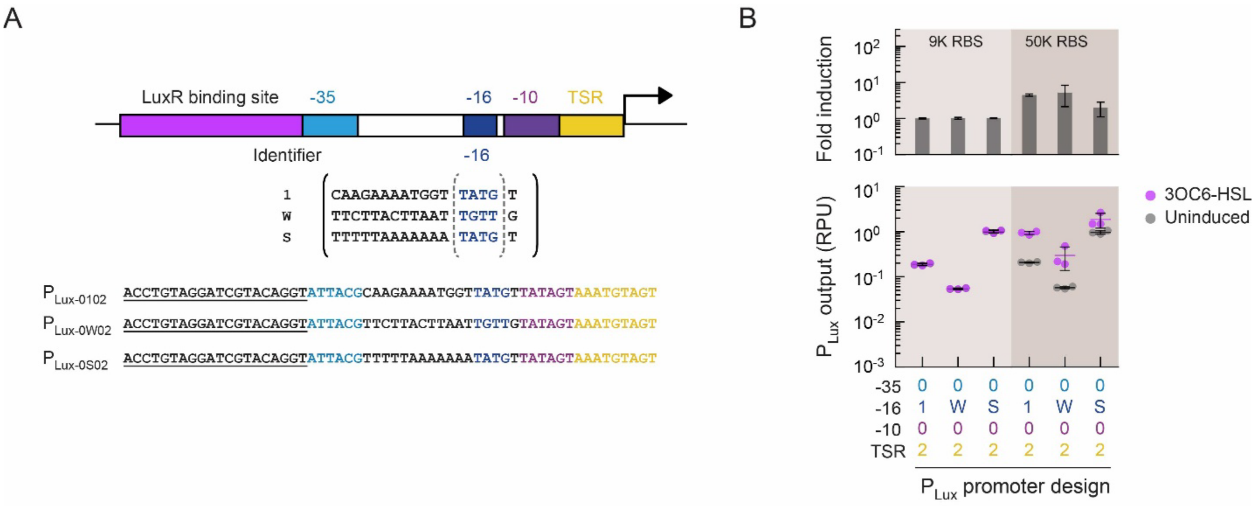

**Figure S5. Sensor performance with spacer sequence substitution. (A)** The $P_{Lux\text{-}0W02}$ and $P_{Lux\text{-}0S02}$ were created by substituting the whole spacer region of $P_{Lux\text{-}0102}$ promoter with 17 bp weak spacer (W) and strong spacer (S) identified from conservation analysis of *B. subtilis* promoters, respectively[2]. **(B)** The 3OC6-HSL sensors containing those two promoter designs were assayed in *B. subtilis* without (gray cycles) or with 20 µM 3OC6-HSL (purple cycles). Markers represent the geometric mean output of a population of at least 10,000 cells. Assays were performed in identical experiments performed on three separate days. For each experiment, the fold induction was calculated as the ratio of the sensor output for the induced to uninduced samples. Bars represent the average fold induction. All error bars represent one standard deviation.

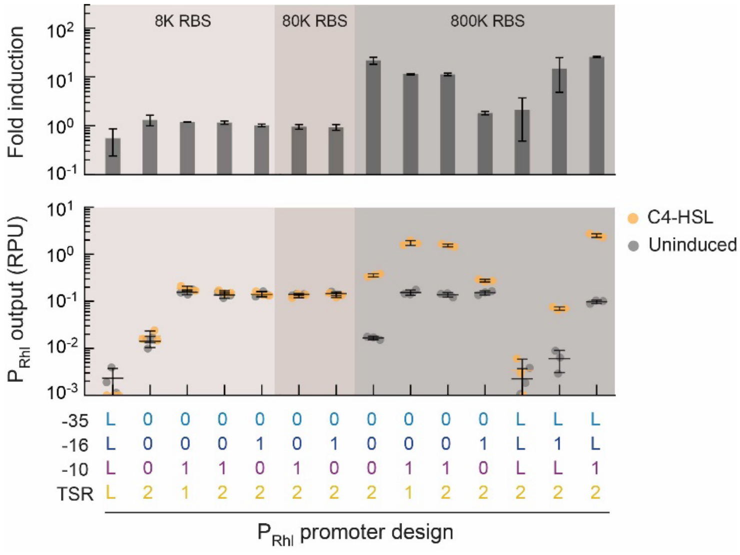

**Figure S6. Engineering synthetic $P_{Rhl}$ promoters for C4-HSL sensors in *B. subtilis*.** The C4-HSL sensors containing different $P_{Rhl}$ promoter designs were assayed without (gray circles) or with 3200 μM C4-HSL (orange circles). Three different RBSs for RhlR were used (beige background colors). Markers represent the geometric mean output of a population of at least 10,000 cells. Assays were performed in identical experiments performed on three separate days. For each experiment, the fold induction was calculated as the ratio of the sensor output for the induced to uninduced samples. Bars represent the average fold induction. All error bars represent one standard deviation.

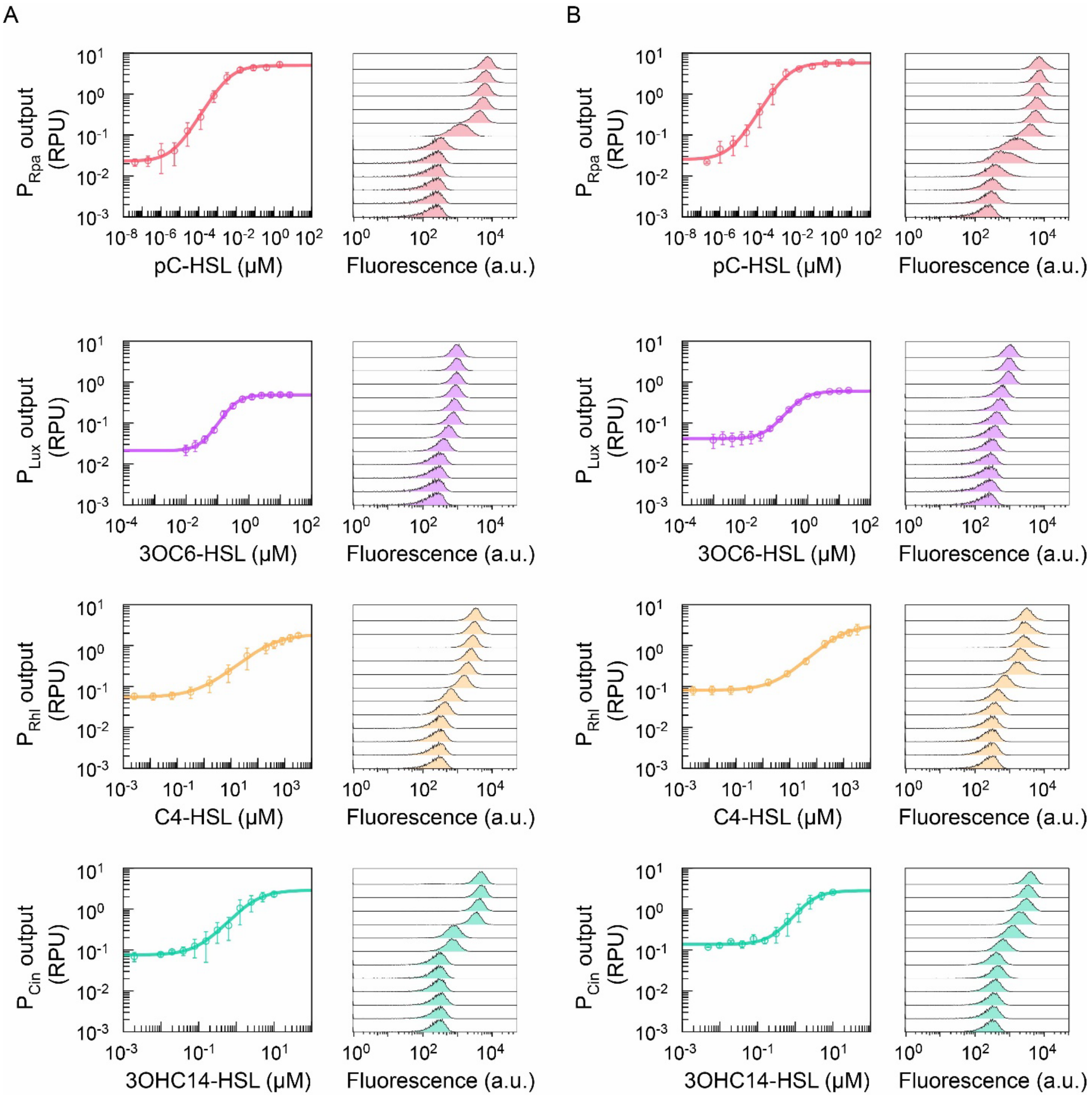

**Figure S7. Comparison of sensor response functions measured at 5 h and 21 h.** The pC-HSL sensor containing $P_{Rpa\text{-}0002}$ (red), 3OC6-HSL sensor containing $P_{Lux\text{-}0002}$ (purple), C4-HSL sensor containing $P_{Rhl\text{-}LL12}$ (orange), and 3OHC14-HSL sensor containing $P_{CinD\text{-}0102}$ (green) are shown. **(A)** Response functions after 5-h induction and **(B)** response functions after 21-h induction were measured. The cell fluorescence was measured by flow cytometry (≥ 10,000 cells per sample). One representative histogram is shown for each sensor at each concentration. The geometric mean of fluorescence in arbitrary units was converted to standard RPU, and the markers represent the average of three independent experiments performed on different days. Error bars are the standard deviation. The colored curves are the response functions fitted to the experimental data.

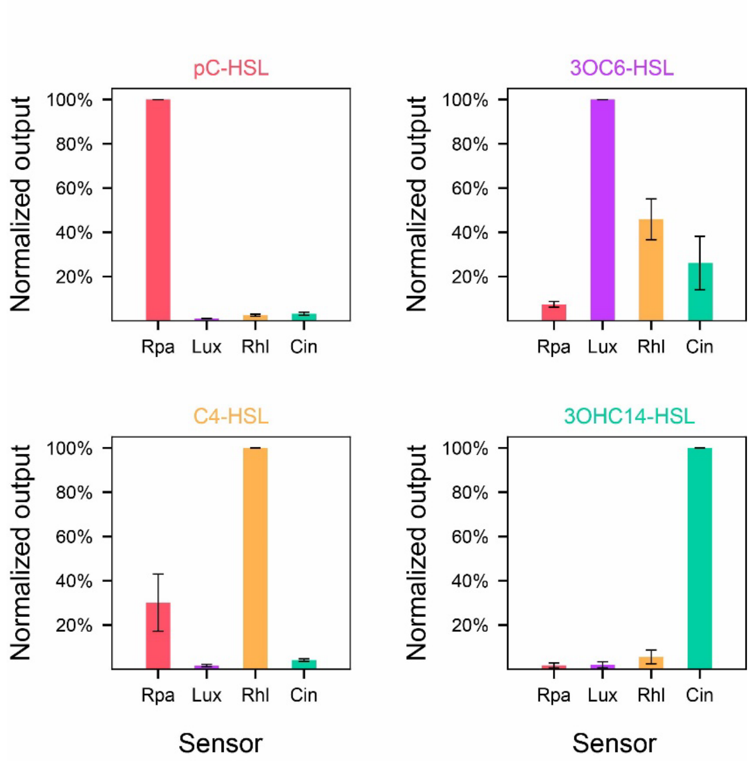

**Figure S8. Signal crosstalk characterization of four HSL sensors.** The pC-HSL sensor containing $P_{Rpa\text{-}0002}$ (red), 3OC6-HSL sensor containing $P_{Lux\text{-}0002}$ (purple), C4-HSL sensor containing $P_{Rhl\text{-}LL12}$ (orange), and 3OHC14-HSL sensor containing $P_{CinD\text{-}0102}$ (green) were characterized for signal crosstalk. Each sensor was induced by cognate and three noncognate HSLs. The concentration of pC-HSL, 3OC6-HSL (C6-HSL), C4-HSL, and 3OHC14-HSL (C14-HSL) were 2 μM, 20 μM, 3200 μM, and 20 μM, respectively. The cell fluorescence was measured by flow cytometry (≥ 10,000 cells per sample). The geometric mean of fluorescence in arbitrary units was converted to standard RPU and then normalized to the output of the cognate sensor of each HSL. Bars are the average of three independent experiments performed on different days. Error bars are the standard deviation.

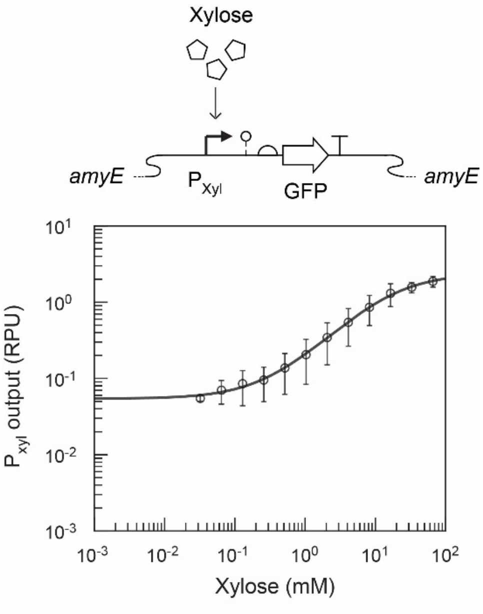

**Figure S9. $P_{Xyl}$ characterization in *B. subtilis* 168.** $P_{xyl}$ sensor was characterized in *B. subtilis* 168 with a range of xylose concentrations. The cell fluorescence was measured by flow cytometry (≥ 10,000 cells per sample). The geometric mean of fluorescence in arbitrary units was converted to standard RPU, and the markers represent the average of three independent experiments performed on different days. Error bars indicate the standard deviation. The solid line is determined by fitting the experimental measurements to the Hill equation ($y_{min}$=0.054, $y_{max}$=2.39, $K$=16.2, and $n$= 0.95).

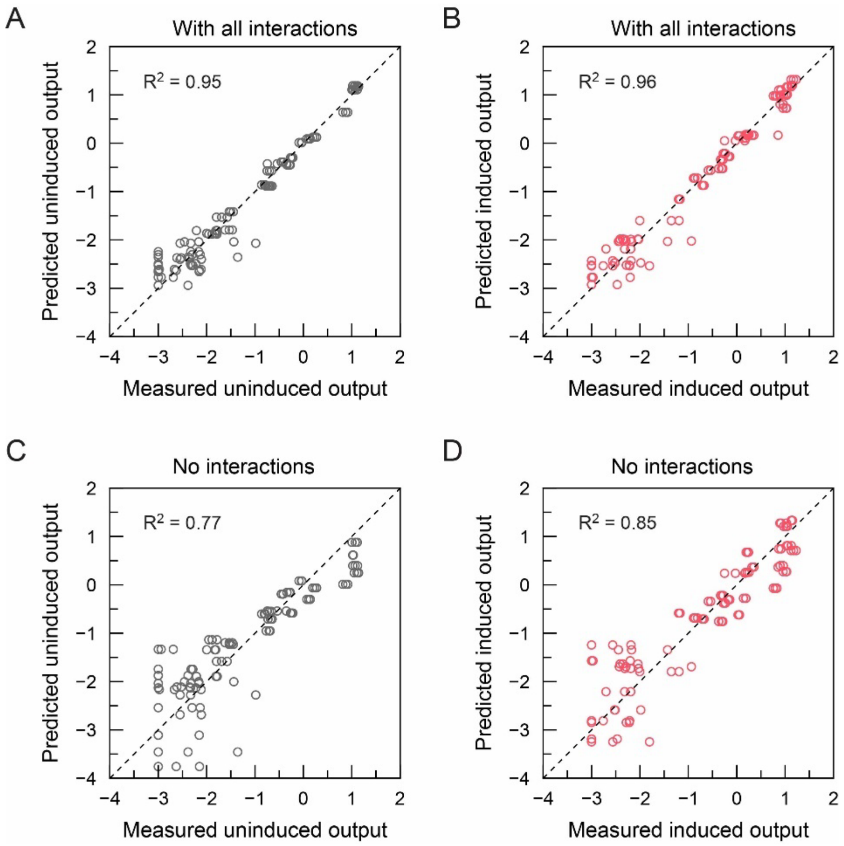

**Figure S10. Linear regression models of $\log_{10}$(uninduced output) and $\log_{10}$(induced output).** The full linear regression models considering 10 terms were constructed for **(A)** log10 of uninduced output and **(B)** log10 of induced output for customed design in Fig.8A. Ten terms included four promoter regions and six pairwise interactions among them. Parameters of the full linear regression models were in SI Table 3 and 4. The linear regression models without interaction terms were constructed for **(C)** log10 of uninduced output and **(D)** log10 of induced output for customed design in Fig.8A.

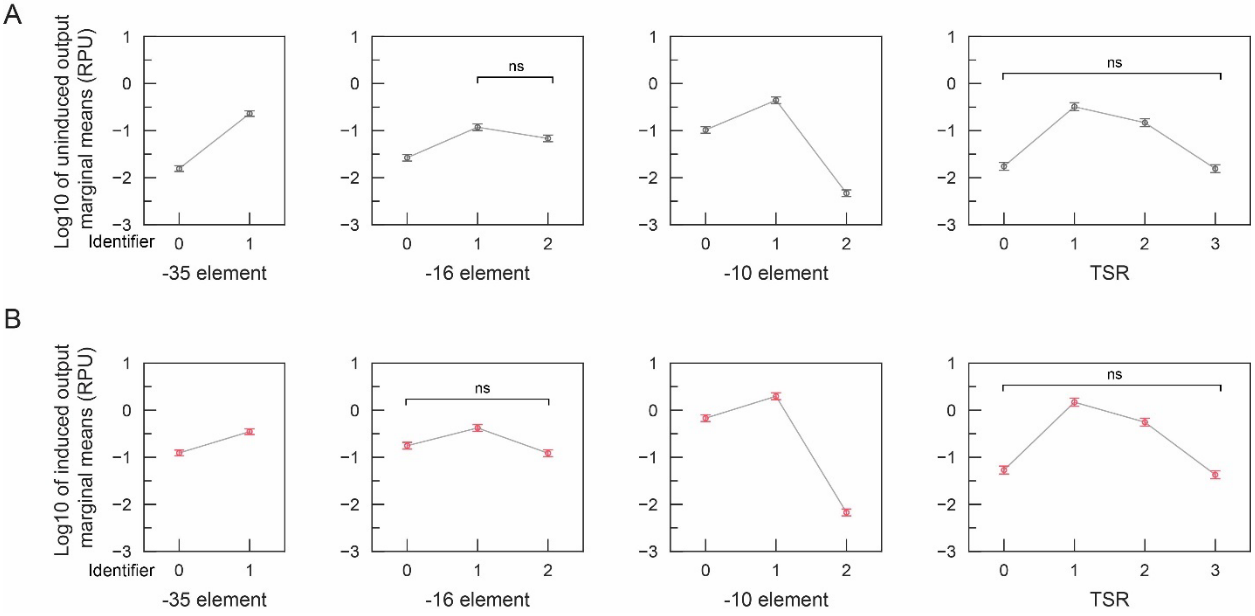

**Figure S11. Reduced linear regression models for effects of promoter region sequences on sensor output.** Promoter regions were compared using **(A)** log10 of uninduced output marginal means and **(B)** log10 of induced output marginal means using corresponding reduced linear regression models. All pairwise differences were significant, except for those denoted as "ns" (non-significant). The significance level was determined using Student's *t*-test, with a significance level of $\alpha = 0.01$. Detailed differences can be found in Table S9.

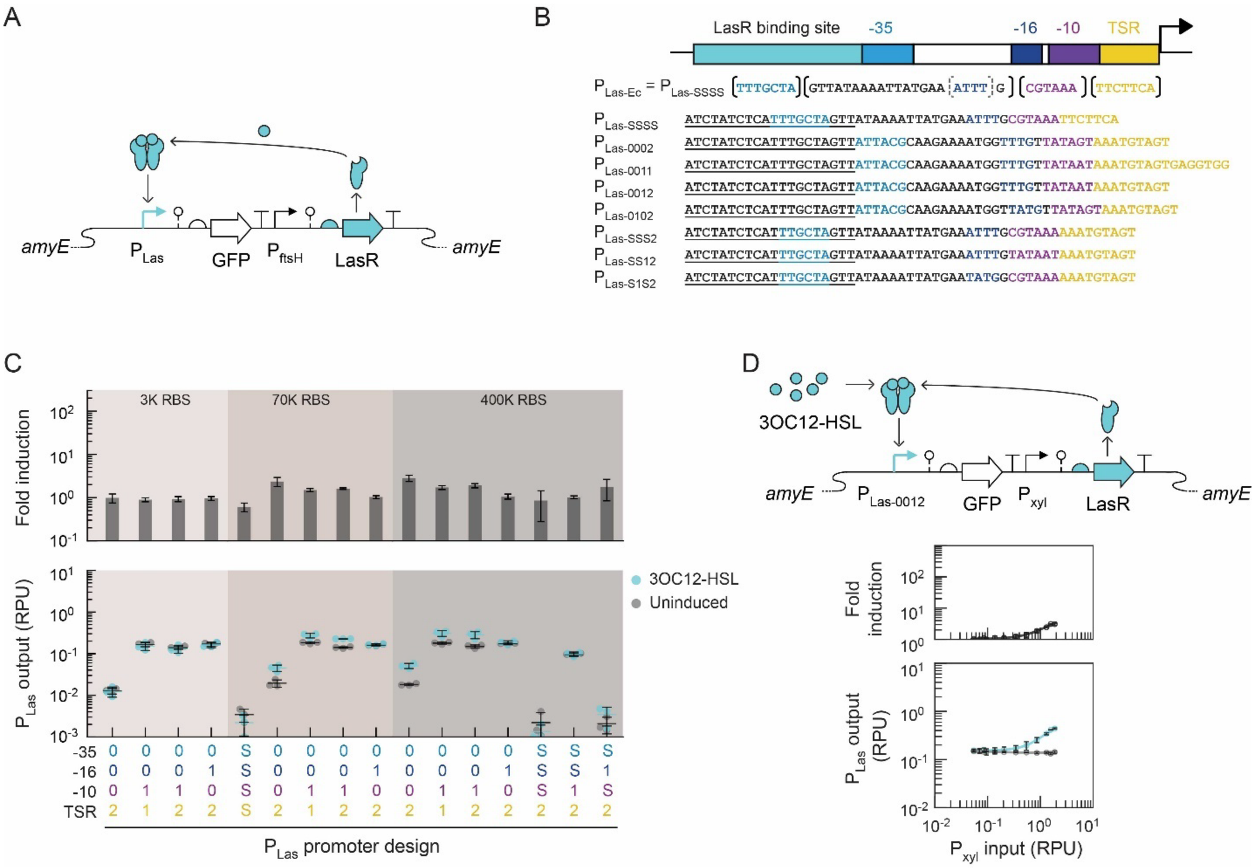

**Figure S12. Engineering synthetic $P_{Las}$ promoter for 3OC12-HSL sensors. (A)** The genetic schematic of the 3OC12-HSL sensor. **(B)** The synthetic $P_{Las}$ promoter sequences. $P_{Las\text{-}0002}$, $P_{Las\text{-}0011}$, $P_{Las\text{-}0012}$, and $P_{Las\text{-}0102}$ promoters were designed by directly substituting the RpaR binding site with the LasR binding site. $P_{Las\text{-}SSS2}$, $P_{Las\text{-}SS12}$, and $P_{Las\text{-}S1S2}$ promoters were designed by altering promoter regions in *E. coli* $P_{Las\text{-}SSSS}$, with S representing Las. **(C)** Synthetic 3OC12-HSL sensor containing $P_{Las}$ promoters and different RBSs of LasR were tested without (gray cycles) or with 20 μM 3OC12-HSL (colored cycles). The RBS strength was calculated using the RBS calculator. The bars represent the average of three independent measurements performed on different days. Error bars are the standard deviation. **(D)** The 3OC12-HSL sensor was tuned by varying expression level of LasR. The cycles represent the average of three experimental measurements performed on different days. Error bars are standard deviation. The solid line is determined by fitting the experimental measurements to the Hill equation.

### Table S1. Genetic part sequences used in this work

| Part Name | Type | DNA sequence | Reference |
|---|---|---|---|
| $P_{FtsH}$ | Promoter | ATGGTTATTGTTTGTATTGGAATGATTTTCTATGGTACTATTGAACATAGTTGTGA | [3] |
| $P_{Xyl}$ | Promoter | CTAAAAAAAATATTGAAAATACTGACGAGGTTATATAAGATGAAAATAAGTTAGTTTGTTTAAACAACAAACTAATAGGTGA | [4] |
| $P_{Rpa-Ec}$ (or $P_{Rpa-0000}$) | Promoter | GC**ACCTGTCCGATCGGACAGTA**TTACGCAAGAAAATGGTTTGTTATAGT*CGAATAT* | [5] |
| $P_{Rpa-0002}$ | Promoter | GC**ACCTGTCCGATCGGACAGTA**TTACGCAAGAAAATGGTTTGTTATAGT*AAATGTAGT* | This study |
| $P_{Rpa-0011}$ | Promoter | GC**ACCTGTCCGATCGGACAGTA**TTACGCAAGAAAATGGTTTGTTATAAT*AAATGTAGTGAGGTGG* | This study |
| $P_{Rpa-0012}$ | Promoter | GC**ACCTGTCCGATCGGACAGTA**TTACGCAAGAAAATGGTTTGTTATAAT*AAATGTAGT* | This study |
| $P_{Rpa-0102}$ | Promoter | GC**ACCTGTCCGATCGGACAGTA**TTACGCAAGAAAATGGTTATGTTATAGT*AAATGTAGT* | This study |
| $P_{Rpa-0112}$ | Promoter | GC**ACCTGTCCGATCGGACAGTA**TTACGCAAGAAAATGGTTATGTTATAAT*AAATGTAGT* | This study |
| $P_{Rpa-0202}$ | Promoter | GC**ACCTGTCCGATCGGACAGTA**TTACGCAAGAAAATGGTTTGTTGTATAGT*AAATGTAGT* | This study |
| $P_{Rpa-0233}$ | Promoter | GC**ACCTGTCCGATCGGACAGTA**TTACGCAAGAAAATGGTTTGTTGTAGTAC*TTTGGGCTAT* | This study |
| $P_{Rpa-1000}$ | Promoter | GC**ACCTGTCCGATCGGACAGTA**TTGACACAAGAAAATGGTTTGTTATAGT*CGAATAT* | This study |
| $P_{Rpa-1002}$ | Promoter | GC**ACCTGTCCGATCGGACAGTA**TTGACACAAGAAAATGGTTTGTTATAGT*AAATGTAGT* | This study |
| $P_{Rpa-1012}$ | Promoter | GC**ACCTGTCCGATCGGACAGTA**TTGACACAAGAAAATGGTTTGTTATAAT*AAATGTAGT* | This study |
| $P_{Rpa-1112}$ | Promoter | GC**ACCTGTCCGATCGGACAGTA**TTGACACAAGAAAATGGTTATGTTATAAT*AAATGTAGT* | This study |
| $P_{Rpa-1113}$ | Promoter | GC**ACCTGTCCGATCGGACAGTA**TTGACACAAGAAAATGGTTATGTTATAAT*TTTGGGCTAT* | This study |
| $P_{Rpa-1213}$ | Promoter | GC**ACCTGTCCGATCGGACAGTA**TTGACACAAGAAAATGGTTTGTTGTATAAT*TTTGGGCTAT* | This study |
| $P_{Rpa-0001}$ | Promoter | GC**ACCTGTCCGATCGGACAGTA**TTACGCAAGAAAATGGTTTGTTATAGT*AAATGTAGTGAGGTGG* | This study |
| $P_{Rpa-0010}$ | Promoter | GC**ACCTGTCCGATCGGACAGTA**TTACGCAAGAAAATGGTTTGTTATAAT*CGAATAT* | This study |
| $P_{Rpa-0021}$ | Promoter | GC**ACCTGTCCGATCGGACAGTA**TTACGCAAGAAAATGGTTTGTTAGTAC*AAATGTAGTGAGGTGG* | This study |
| $P_{Rpa-0023}$ | Promoter | GC**ACCTGTCCGATCGGACAGTA**TTACGCAAGAAAATGGTTTGTTAGTAC*TTTGGGCTAT* | This study |
| $P_{Rpa-0100}$ | Promoter | GC**ACCTGTCCGATCGGACAGTA**TTACGCAAGAAAATGGTTATGTTATAGT*CGAATAT* | This study |
| $P_{Rpa-0101}$ | Promoter | GC**ACCTGTCCGATCGGACAGTA**TTACGCAAGAAAATGGTTATGTTATAGT*AAATGTAGTGAGGTGG* | This study |
| $P_{Rpa-0111}$ | Promoter | GC**ACCTGTCCGATCGGACAGTA**TTACGCAAGAAAATGGTTATGTTATAAT*AAATGTAGTGAGGTGG* | This study |
| $P_{Rpa-0113}$ | Promoter | GC**ACCTGTCCGATCGGACAGTA**TTACGCAAGAAAATGGTTATGTTATAAT*TTTGGGCTAT* | This study |
| $P_{Rpa-0122}$ | Promoter | GC**ACCTGTCCGATCGGACAGTA**TTACGCAAGAAAATGGTTATGTTAGTAC*AAATGTAGT* | This study |

| | | | |
|---|---|---|---|
| $P_{Rpa-0123}$ | Promoter | GC**ACCTGTCCGATCGGACAGTA**ATTACGCAAGAAAATGGTTATGTTAGTAC*TTTGGGCTAT* | This study |
| $P_{Rpa-0200}$ | Promoter | GC**ACCTGTCCGATCGGACAGTA**ATTACGCAAGAAAATGGTTTGTTGTATAGT*CGAATAT* | This study |
| $P_{Rpa-0203}$ | Promoter | GC**ACCTGTCCGATCGGACAGTA**ATTACGCAAGAAAATGGTTTGTTGTATAGT*TTTGGGCTAT* | This study |
| $P_{Rpa-0211}$ | Promoter | GC**ACCTGTCCGATCGGACAGTA**ATTACGCAAGAAAATGGTTTGTTGTATAAT*AAATGTAGTGAGGTGG* | This study |
| $P_{Rpa-0212}$ | Promoter | GC**ACCTGTCCGATCGGACAGTA**ATTACGCAAGAAAATGGTTTGTTGTATAAT*AAATGTAGT* | This study |
| $P_{Rpa-0220}$ | Promoter | GC**ACCTGTCCGATCGGACAGTA**ATTACGCAAGAAAATGGTTTGTTGTAGTAC*CGAATAT* | This study |
| $P_{Rpa-0221}$ | Promoter | GC**ACCTGTCCGATCGGACAGTA**ATTACGCAAGAAAATGGTTTGTTGTAGTAC*AAATGTAGTGAGGTGG* | This study |
| $P_{Rpa-1003}$ | Promoter | GC**ACCTGTCCGATCGGACAGTA**TTGACACAAGAAAATGGTTTGTTATAGT*TTTGGGCTAT* | This study |
| $P_{Rpa-1011}$ | Promoter | GC**ACCTGTCCGATCGGACAGTA**TTGACACAAGAAAATGGTTTGTTATAAT*AAATGTAGTGAGGTGG* | This study |
| $P_{Rpa-1013}$ | Promoter | GC**ACCTGTCCGATCGGACAGTA**TTGACACAAGAAAATGGTTTGTTATAAT*TTTGGGCTAT* | This study |
| $P_{Rpa-1020}$ | Promoter | GC**ACCTGTCCGATCGGACAGTA**TTGACACAAGAAAATGGTTTGTTAGTAC*CGAATAT* | This study |
| $P_{Rpa-1022}$ | Promoter | GC**ACCTGTCCGATCGGACAGTA**TTGACACAAGAAAATGGTTTGTTAGTAC*AAATGTAGT* | This study |
| $P_{Rpa-1102}$ | Promoter | GC**ACCTGTCCGATCGGACAGTA**TTGACACAAGAAAATGGTTATGTTATAGT*AAATGTAGT* | This study |
| $P_{Rpa-1103}$ | Promoter | GC**ACCTGTCCGATCGGACAGTA**TTGACACAAGAAAATGGTTATGTTATAGT*TTTGGGCTAT* | This study |
| $P_{Rpa-1110}$ | Promoter | GC**ACCTGTCCGATCGGACAGTA**TTGACACAAGAAAATGGTTATGTTATAAT*CGAATAT* | This study |
| $P_{Rpa-1120}$ | Promoter | GC**ACCTGTCCGATCGGACAGTA**TTGACACAAGAAAATGGTTATGTTAGTAC*CGAATAT* | This study |
| $P_{Rpa-1121}$ | Promoter | GC**ACCTGTCCGATCGGACAGTA**TTGACACAAGAAAATGGTTATGTTAGTAC*AAATGTAGTGAGGTGG* | This study |
| $P_{Rpa-1201}$ | Promoter | GC**ACCTGTCCGATCGGACAGTA**TTGACACAAGAAAATGGTTTGTTGTATAGT*AAATGTAGTGAGGTGG* | This study |
| $P_{Rpa-1202}$ | Promoter | GC**ACCTGTCCGATCGGACAGTA**TTGACACAAGAAAATGGTTTGTTGTATAGT*AAATGTAGT* | This study |
| $P_{Rpa-1210}$ | Promoter | GC**ACCTGTCCGATCGGACAGTA**TTGACACAAGAAAATGGTTTGTTGTATAAT*CGAATAT* | This study |
| $P_{Rpa-1222}$ | Promoter | GC**ACCTGTCCGATCGGACAGTA**TTGACACAAGAAAATGGTTTGTTGTAGTAC*AAATGTAGT* | This study |
| $P_{Rpa-1223}$ | Promoter | GC**ACCTGTCCGATCGGACAGTA**TTGACACAAGAAAATGGTTTGTTGTAGTAC*TTTGGGCTAT* | This study |
| $P_{Lux-Ec}$ (or $P_{Lux-XXXX}$) | Promoter | ATAGCTTCTTACCGG**ACCTGTAGGATCGTACAGGT**TTACGCAAGAAAATGGTTTGTTACTTTCGAATAAA | [6] |
| $P_{Lux-0002}$ | Promoter | **ACCTGTAGGATCGTACAGGT**ATTACGCAAGAAAATGGTTTGTTATAGT*AAATGTAGT* | This study |
| $P_{Lux-0011}$ | Promoter | **ACCTGTAGGATCGTACAGGT**ATTACGCAAGAAAATGGTTTGTTATAAT*AAATGTAGTGAGGTGG* | This study |
| $P_{Lux-0012}$ | Promoter | **ACCTGTAGGATCGTACAGGT**ATTACGCAAGAAAATGGTTTGTTATAAT*AAATGTAGT* | This study |
| $P_{Lux-0102}$ | Promoter | **ACCTGTAGGATCGTACAGGT**ATTACGCAAGAAAATGGTTATGTTATAGT*AAATGTAGT* | This study |

| | | | |
|---|---|---|---|
| $P_{\text{Lux-0W02}}$ | Promoter | **ACCTGTAGGATCGTACAGGT**ATTACGTTCTTACTTAATTGTTGTATAGT*AAATGTAGT* | This study |
| $P_{\text{Lux-0S02}}$ | Promoter | **ACCTGTAGGATCGTACAGGT**ATTACGTTTTTAAAAAAATATGTTATAGT*AAATGTAGT* | This study |
| $P_{\text{Lux-4012}}$ | Promoter | **ACCTGTAGGATCGTACAGGT**CCCGGGCAAGAAAATGGTTTGTTATAAT*AAATGTAGT* | This study |
| $P_{\text{Lux-3002}}$ | Promoter | **ACCTGTAGGATCGTACAGGT**CTGACACAAGAAAATGGTTTGTTATAGT*AAATGTAGT* | This study |
| $P_{\text{Lux-2032}}$ | Promoter | **ACCTGTAGGATCGTACAGGT**TTGTGACAAGAAAATGGTTTGTGATAAT*AAATGTAGT* | This study |
| $P_{\text{Lux-1062}}$ | Promoter | **ACCTGTAGGATCGTACAGGT**TAGACACAAGAAAATGGTTTGTTATGTT*AAATGTAGT* | This study |
| $P_{\text{Lux-1052}}$ | Promoter | **ACCTGTAGGATCGTACAGGT**TAGACACAAGAAAATGGTTTGTTACTGT*AAATGTAGT* | This study |
| $P_{\text{Lux-1042}}$ | Promoter | **ACCTGTAGGATCGTACAGGT**TAGACACAAGAAAATGGTTTGTTAAATT*AAATGTAGT* | This study |
| $P_{\text{Rhl-Ec}}$ (or $P_{\text{Rhl-LLLL}}$) | Promoter | **TCCTGTGAAATCTGGCAGTT**ACCGTTAGCTTTCGAATTGGCTAAAAA*GTGTTC* | [7] |
| $P_{\text{Rhl-0002}}$ | Promoter | **TCCTGTGAAATCTGGCAGTT**ATTACGCAAGAAAATGGTTTGTTATAGT*AAATGTAGT* | This study |
| $P_{\text{Rhl-0011}}$ | Promoter | **TCCTGTGAAATCTGGCAGTT**ATTACGCAAGAAAATGGTTTGTTATAAT*AAATGTAGTGAGGTGG* | This study |
| $P_{\text{Rhl-0012}}$ | Promoter | **TCCTGTGAAATCTGGCAGTT**ATTACGCAAGAAAATGGTTTGTTATAAT*AAATGTAGT* | This study |
| $P_{\text{Rhl-0102}}$ | Promoter | **TCCTGTGAAATCTGGCAGTT**ATTACGCAAGAAAATGGTTATGTTATAGT*AAATGTAGT* | This study |
| $P_{\text{Rhl-LLL2}}$ | Promoter | **TCCTGTGAAATCTGGCAGTT**ACCGTTAGCTTTCGAATTGGCTAAAAA*AAATGTAGT* | This study |
| $P_{\text{Rhl-LL12}}$ | Promoter | **TCCTGTGAAATCTGGCAGTT**ACCGTTAGCTTTCGAATTGGCTATAAT*AAATGTAGT* | This study |
| $P_{\text{Rhl-L1L2}}$ | Promoter | **TCCTGTGAAATCTGGCAGTT**ACCGTTAGCTTTCGAATATGCTAAAAA*AAATGTAGT* | This study |
| $P_{\text{Cin-Ec}}$ (or $P_{\text{Cin-NNNN}}$) | Promoter | CCCTTTGTGCGTCCAAACGGACGCACGGCGCTCTAAAGCGGGTCGCGATCTTTCAGATTCGCTCCTCGCGCTTTCAGTCTTTGTTTTGGCGCATGTCGTTATCGCAAAACCGCTGCACACTTTTGCGCGACATGCTCTGATCCCCCTCATCTGGGGGGGCCTATCTGAGGGAATTTCCGATCCGGCTCGCCTGAACCATTCTGCTTTCCACGAACTTGAAAACGCT | [8] |
| $P_{\text{CinD-0002}}$ | Promoter | **CATGCTCTGATCCCCCTCATCTGGGGGGGCCTATCTGAGGGAA**ATTACGCAAGAAAATGGTTTGTTATAGT*AAATGTAGT* | This study |
| $P_{\text{CinD-0011}}$ | Promoter | **CATGCTCTGATCCCCCTCATCTGGGGGGGCCTATCTGAGGGAA**ATTACGCAAGAAAATGGTTTGTTATAAT*AAATGTAGTGAGGTGGT* | This study |
| $P_{\text{CinD-0012}}$ | Promoter | **CATGCTCTGATCCCCCTCATCTGGGGGGGCCTATCTGAGGGAA**ATTACGCAAGAAAATGGTTTGTTATAAT*AAATGTAGT* | This study |
| $P_{\text{CinD-0102}}$ | Promoter | **CATGCTCTGATCCCCCTCATCTGGGGGGGCCTATCTGAGGGAA**ATTACGCAAGAAAATGGTTATGTTATAGT*AAATGTAGT* | This study |
| $P_{\text{CinA-0002}}$ | Promoter | **GGCGCATGTCGTTATCGCAAAACCGCTGCACACTTTTGCGCGACATGCTCTGATCCCCCTCATCTGGGGGGGCCTATCTGAGGGAA**ATTACGCAAGAAAATGGTTTGTTATAGT*AAATGTAGT* | This study |
| $P_{\text{CinA-0011}}$ | Promoter | **GGCGCATGTCGTTATCGCAAAACCGCTGCACACTTTTGCGCGACATGCTCTGATCCCCCTCATCTGGGGGGGCCTATC** | This study |

| | | | |
|---|---|---|---|
| | | **TGAGGGAA**ATTACGCAAGAAAATGGTTTGTTATAAT*AAATGTAGTGAGGTGGT* | |
| $P_{CinA\text{-}0012}$ | Promoter | **GGCGCATGTCGTTATCGCAAAACCGCTGCACACTTTTGCGCGACATGCTCTGATCCCCCTCATCTGGGGGGGCCTATCTGAGGGAA**ATTACGCAAGAAAATGGTTTGTTATAAT*AAATGTAGT* | This study |
| $P_{CinA\text{-}0102}$ | Promoter | **GGCGCATGTCGTTATCGCAAAACCGCTGCACACTTTTGCGCGACATGCTCTGATCCCCCTCATCTGGGGGGGCCTATCTGAGGGAA**ATTACGCAAGAAAATGGTTATGTTATAGT*AAATGTAGT* | This study |
| $P_{CinB\text{-}0002}$ | Promoter | **GGGGGCCTATCTGAGGGAA**ATTACGCAAGAAAATGGTTTGTTATAGT*AAATGTAGT* | This study |
| $P_{CinB\text{-}0011}$ | Promoter | **GGGGGCCTATCTGAGGGAA**ATTACGCAAGAAAATGGTTTGTTATAAT*AAATGTAGTGAGGTGGT* | This study |
| $P_{CinB\text{-}0012}$ | Promoter | **GGGGGCCTATCTGAGGGAA**ATTACGCAAGAAAATGGTTTGTTATAAT*AAATGTAGT* | This study |
| $P_{CinB\text{-}0102}$ | Promoter | **GGGGGCCTATCTGAGGGAA**ATTACGCAAGAAAATGGTTATGTTATAGT*AAATGTAGT* | This study |
| $P_{CinC\text{-}0002}$ | Promoter | **CATGCTCTGATCCCCCTCATCTGGGGGGG**ATTACGCAAGAAAATGGTTTGTTATAGT*AAATGTAGT* | This study |
| $P_{CinC\text{-}0011}$ | Promoter | **CATGCTCTGATCCCCCTCATCTGGGGGGG**ATTACGCAAGAAAATGGTTTGTTATAAT*AAATGTAGTGAGGTGGT* | This study |
| $P_{CinC\text{-}0012}$ | Promoter | **CATGCTCTGATCCCCCTCATCTGGGGGGG**ATTACGCAAGAAAATGGTTTGTTATAAT*AAATGTAGT* | This study |
| $P_{CinC\text{-}0102}$ | Promoter | **CATGCTCTGATCCCCCTCATCTGGGGGGG**ATTACGCAAGAAAATGGTTATGTTATAGT*AAATGTAGT* | This study |
| $P_{Las\text{-}Ec}$ (or $P_{Las\text{-}0000}$) | Promoter | TTCGAGCCTAGCAAGGGTCCGGGTTCACCGAA**ATCTATCTCATTTGCTAGTT**ATAAAATTATGAAATTTGCGTAAA*TTCTTCA* | [5] |
| $P_{Las\text{-}0002}$ | Promoter | **ATCTATCTCATTTGCTAGTT**ATTACGCAAGAAAATGGTTTGTTATAGT*AAATGTAGT* | This study |
| $P_{Las\text{-}0011}$ | Promoter | **ATCTATCTCATTTGCTAGTT**ATTACGCAAGAAAATGGTTTGTTATAAT*AAATGTAGTGAGGTGG* | This study |
| $P_{Las\text{-}0012}$ | Promoter | **ATCTATCTCATTTGCTAGTT**ATTACGCAAGAAAATGGTTTGTTATAAT*AAATGTAGT* | This study |
| $P_{Las\text{-}0102}$ | Promoter | **ATCTATCTCATTTGCTAGTT**ATTACGCAAGAAAATGGTTATGTTATAGT*AAATGTAGT* | This study |
| $P_{Las\text{-}SSS2}$ | Promoter | TTCGAGCCTAGCAAGGGTCCGGGTTCACCGAA**ATCTATCTCATTTGCTAGTT**ATAAAATTATGAAATTTGCGTAAA*AAATGTAGT* | This study |
| $P_{Las\text{-}SS12}$ | Promoter | TTCGAGCCTAGCAAGGGTCCGGGTTCACCGAA**ATCTATCTCATTTGCTAGTT**ATAAAATTATGAAATTTGTATAAT*AAATGTAGT* | This study |
| $P_{Las\text{-}S1S2}$ | Promoter | TTCGAGCCTAGCAAGGGTCCGGGTTCACCGAA**ATCTATCTCATTTGCTAGTT**ATAAAATTATGAATATGGCGTAAA*AAATGTAGT* | This study |
| RiboJ | Insulator | AGCTGTCACCGGATGTGCTTTCCGGTCTGATGAGTCCGTGAGGACGAAACAGCCTCTACAAATAATTTTGTTTAA | [9] |
| PlmJ | Insulator | AGTCATAAGTCTGGGCTAAGCCCACTGATGAGTCGCTGAAATGCGACGAAACTTATGACCTCTACAAATAATTTTGTTTAA | [9] |
| GFP-rbs | RBS | TACTAGAGGAAGCTTACATAAGGAGGAACTACT | This study |
| RpaR rbs1 | RBS (8K) | TACGACTGCGAGGTCATAAGAAAGGAGGTTACGTT | This study |

| | | | |
|---|---|---|---|
| LuxR rbs1 | RBS (9K au) | TACGCAGTTTCTACAGAAGAGATTAAA | This study |
| LuxR rbs2 | RBS (50K au) | TACGCCATATAAAAAGGAGACACTACCTCA | This study |
| RhlR rbs1 | RBS (8K au) | TACGAAATAGGAGCCAAGACTT | This study |
| RhlR rbs2 | RBS(80K au) | TACGGGTTGAGTCTGGGAGAATATTTTTT | This study |
| RhlR rbs3 | RBS(800K au) | TACGTTAGGGAGTAAGTAAAGGAGGTAAAATATT | This study |
| CinR rbs1 | RBS (15K au) | TACGTCCGTACATAGCAGAAGGGGGTAAATA | This study |
| LasR rbs1 | RBS (3K au) | TACGCTGAAAAGTTTCGAGGAATCTTTCAC | This study |
| LasR rbs2 | RBS (70K au) | TACGTAGAGTTGAAGGAGAGATATTTTA | This study |
| LasR rbs3 | RBS (400K au) | TACGGAGAGTCGCACAAAGGAGGTATTTTTT | This study |
| *rpaR* | Gene | ATGATCGTCGGCGAAGATCAGCTTTGGGGACGGCGTGCG<br>CTGGAATTCGTCGATTCCGTCGAACGGCTCGAGGCGCCG<br>GCGCTGATCAGCCGGTTCGAATCGCTGATCGCGAGCTGC<br>GGATTTACCGCCTACATCATGGCCGGCCTGCCGTCGCGC<br>AATGCCGGACTACCGGAGCTGACGCTGGCCAATGGCTGG<br>CCGCGAGACTGGTTCGATCTGTATGTCAGCGAAAACTTCA<br>GCGCGGTCGATCCGGTGCCGCGCCACGGCGCTACCACG<br>GTTCATCCTTTCGTATGGTCCGATGCACCCTACGACCGCG<br>ACCGTGATCCGGCCGCCCACCGGGTCATGACCCGGGCG<br>GCGGAATTCGGACTGGTCGAGGGTTACTGCATTCCGCTG<br>CACTACGACGACGGTAGCGCCGCGATCAGCATGGCCGGC<br>AAGGATCCGGACCTCAGCCCGGCCGCGCGCGGCGCGAT<br>GCAGCTGGTCAGCATCTACGCGCATAGTCGCCTGCGCGC<br>ACTCAGCCGGCCAAAGCCGATCCGGCGCAACCGGCTCA<br>CGCCGCGCGAGTGCGAGATCCTGCAATGGGCAGCGCAG<br>GGCAAGACCGCCTGGGAAATCTCGGTAATCCTCTGCATCA<br>CCGAACGCACGGTGAAATTCCATCTGATCGAAGCCGCCC<br>GCAAGCTCGACGCCGCCAACCGCACCGCGGCGGTTGCC<br>AAGGCATTGACGCTCGGATTGATCCGTTTGTGA | [5] |
| *luxR* | Gene | ATGAAAAACATAAATGCCGACGACACATACAGAATAATTAAT<br>AAAATTAAAGCTTGTAGAAGCAATAATGATATTAATCAATGC<br>TTATCTGATATGACTAAAATGGTACATTGTGAATATTATTTAC<br>TCGCGATCATTTATCCTCATTCTATGGTTAAATCTGATATTT<br>CAATCCTAGATAATTACCCTAAAAAATGGAGGCAATATTATG<br>ATGACGCTAATTTAATAAAATATGATCCTATAGTAGATTATTC<br>TAACTCCAATCATTCACCAATTAATTGGAATATATTTGAAAA<br>CAATGCTGTAAATAAAAAATCTCCAAATGTAATTAAAGAAGC<br>GAAAACATCAGGTCTTATCACTGGGTTTAGTTTCCCTATTC<br>ATACGGCTAACAATGGCTTCGGAATGCTTAGTTTTGCACAT<br>TCAGAAAAAGACAACTATATAGATAGTTTATTTTTACATGCG<br>TGTATGAACATACCATTAATTGTTCCTTCTCTAGTTGATAAT<br>TATCGAAAAATAAATATAGCAAATAATAAATCAAACAACGATT<br>TAACCAAAAGAGAAAAAGAATGTTTAGCGTGGGCATGCGA<br>AGGAAAAAGCTCTTGGGATATTTCAAAAATATTAGGTTGCA<br>GTGAGCGTACTGTCACTTTCCATTTAACCAATGCGCAAAT<br>GAAACTCAATACAACAAACCGCTGCCAAAGTATTTCTAAAG | [8] |

| | | | |
|---|---|---|---|
| | | CAATTTTAACAGGAGCAATTGATTGCCCATACTTTAAAAATT AA | |
| *rhlR* | Gene | ATGAGGAATGACGGAGGCTTTTTGCTGTGGTGGGACGGT TTGCGTAGCGAGATGCAGCCGATCCACGACAGCCAGGGC GTGTTCGCCGTCCTGGAAAAGGAAGTGCGGCGCCTGGG CTTCGATTACTACGCCTATGGCGTGCGCCACACGATTCCC TTCACCCGGCCCAAGACCGAGGTCCATGGCACCTATCCC AAGGCCTGGCTGGAGCGATACCAGATGCAGAACTACGGG GCCGTGGATCCGGCGATCCTCAACGGCCTGCGCTCCTCG GAAATGGTGGTCTGGAGCGACAGCCTGTTCGACCAGAGC CGGATGCTCTGGAACGAGGCTCGCGATTGGGGCCTCTGT GTCGGCGCGACCTTGCCGATCCGCGCGCCGAACAATTTG CTCAGCGTGCTTTCCGTGGCGCGCGACCAGCAGAACATC TCCAGCTTCGAGCGCGAGGAAATCCGCCTGCGGCTGCGT TGCATGATCGAGTTGCTGACCCAGAAGCTGACCGACCTG GAGCATCCGATGCTGATGTCCAACCCGGTCTGCCTGAGC CATCGCGAACGCGAGATCCTGCAATGGACCGCCGACGGC AAGAGTTCCGGGGAAATCGCCATCATCCTGAGCATCTCCG AGAGCACGGTGAACTTCCACCACAAGAACATCCAGAAGA AGTTCGACGCGCCGAACAAGACGCTGGCTGCCGCCTAC GCCGCGGCGCTGGGCCTCATCTAA | [5] |
| *cinR* | Gene | ATGATTGAGAATACCTATAGCGAAAAGTTCGAGTCCGCGTT CGAACAGATCAAAGCGGCGGCCAACGTGGATGCCGCCAT CCGTATTCTCCAGGCGGAATATAACCTCGATTTCGTCACCT ACCATCTCGCCCAGACAATCGCGAGCAAGATCGATTCGC CCTTCGTGCGCACCACCTATCCGGATGCCTGGGTTTCCC GTTACCTCCTCAACTGCTATGTGAAGGTCGATCCGATCAT CAAGCAGGGCTTCGAACGCCAGCTGCCCTTCGACTGGA GCGAGGTCGAACCGACGCCGGAGGCCTATGCCATGCTG GTCGACGCCCAGAAACACGGCATCGATGACAATGGCTAC TCCATCCCCGTCGCCGACAAGGCGCAGCGCCGCGCCCT GCTGTCGCTGAATGCCCATATACCGGCCGACGAATGGAC CGAGCTCGTGCGCCGCTGCCGCAATGAGTGGATCGAGAT CGCCCATCTGATCCACCGCAAGGCCGTATATGAGCTGCAT GGCGAAAACGATCCGGTGCCGGCATTGTCGCCGCGCGA GATCGAGTGTCTGCACTGGACCGCCCTCGGCAAGGATTA CAAGGATATTTCGGTCATCCTGGGCATATCAGAGCATACCA CACGCGATTACCTGAAAACCGCCCGCTTCAGGCTCGGCT GCACCACGATCTCGGCCGCCGCGTCGCGGGCTGTTCAAT TGCGCATCATCAATCCCTATAGGATCCGCATGACGCGACG TAATTGGTAA | [8] |
| *lasR* | Gene | ATGGCCTTGGTTGACGGTTTTCTTGAGCTGGAACGCTCAA GTGGAAAATTGGAGTGGAGCGCCATCCTGCAGAAGATGG CGAGCGACCTTGGATTCTCGAAGATCCTGTTCGGCCTGTT GCCTAAGGACAGCCAGGACTACGAGAACGCCTTCATCGT CGGCAACTACCCGGCCGCCTGGCGCGAGCATTACGACC GGGCTGGCTACGCGCGGGTCGACCCGACGGTCAGTCAC TGTACCCAGAGCGTACTGCCGATTTTCTGGGAACCGTCCA TCTACCAGACGCGAAAGCAGCACGAGTTCTTCGAGGAAG CCTCGGCCGCCGGCCTGGTGTATGGGCTGACCATGCCG CTGCATGGTGCTCGCGGCGAACTCGGCGCGCTGAGCCT CAGCGTGGAAGCGGAAAACCGGGCCGAGGCCAACCGTT TCATGGAGTCGGTCCTGCCGACCCTGTGGATGCTCAAGG ACTACGCACTGCAGAGCGGTGCCGGACTGGCCTTCGAAC ATCCGGTCAGCAAACCGGTGGTTCTGACCAGCCGGGAGA AGGAAGTGTTGCAGTGGTGCGCCATCGGCAAGACCAGTT | [5] |

| | | | |
|---|---|---|---|
| | | GGGAGATATCGGTTATCTGCAACTGCTCGGAAGCCAATGTGAACTTCCATATGGGAAATATTCGGCGGAAGTTCGGTGTGACCTCCCGCCGCGTAGCGGCCATTATGGCCGTTAATTTGGGTCTTATTACTCTCTGA | |
| *gfp* | Gene | ATGAGTAAAGGAGAAGAACTTTTCACTGGAGTTGTCCCAATTCTTGTTGAATTAGATGGTGATGTTAATGGGCACAAATTTTCTGTCAGTGGAGAGGGTGAAGGTGATGCAACATACGGAAAACTTACCCTTAAATTTATTTGCACTACTGGAAAACTACCTGTTCCATGGCCAACACTTGTCACTACTTTCGCGTATGGaCTTCAATGCTTTGCGAGATACCCAGATCATATGAAACAGCATGACTTTTTCAAGAGTGCCATGCCCGAAGGTTATGTACAGGAAAGAACTATATTTTTCAAAGATGACGGGAACTACAAGACACGTGCTGAAGTCAAGTTTGAAGGTGATACCCTTGTTAATAGAATCGAGTTAAAAGGTATTGATTTTAAAGAAGATGGAAACATTCTTGGACACAAATTGGAATACAACTATAACTCACACAATGTATACATCATGGCAGACAAACAAAAGAATGGAATCAAAGTTAACTTCAAAATTAGACACAACATTGAAGATGGAAGCGTTCAACTAGCAGACCATTATCAACAAAATACTCCAATTGGCGATGGCCCTGTCCTTTTACCAGACAACCATTACCTGTCCTACCAATCTGCCCTTTCGAAAGATCCCAACGAAAAGAGAGAtCACATGGTCCTTCTTGAGTTTGTAACAGCTGCTGGGATTACACATGGCATGGATGAACTATACAAATAA | This study (modified from [10]) |
| L3S2P21 | Terminator | CTCGGTACCAAATTCCAGAAAAGAGGCCTCCCGAAAGGGGGGCCTTTTTTCGTTTTGGTCC | [11] |
| L3S2P15 | Terminator | CTCGGTACCAAATTCCAGAAAAGAGACGCTTTTAGAGCGTCTTTTTTCGTTTTGGTCC | [11] |
| *amyE* front | Homology arm | ATGTTTGCAAAACGATTCAAAACCTCTTTACTGCCGTTATTCGCTGGATTTTTATTGCTGTTTCATTTGGTTCTGGCAGGACCGGCGGCTGCGAGTGCTGAAACGGCGAACAAATCGAATGAGCTTACAGCACCGTCGATCAAAAGCGGAACCATTCTTCATGCATGGAATTGGTCGTTCAATACGTTAAAACACAATATGAAGGATATTCATGATGCAGGATATACAGCCATTCAGACATCTCCGATTAACCAAGTAAAGGAAGGGAATCAAGGAGATAAAAGCATGTCGAACTGGTACTGGCTGTATCAGCCGACATCGTATCAAATTGGCAACCGTTACTTAGGTACTGAACAAGAATTTAAAGAAATGTGTGCAGCCGCTGAAGAATATGGCATAAAGGTCATTGTTGACGCGGTCATCAATCATACCACCAGTGATTATGCCGCGATTTCCAATGAGGTTAAGAGTATTCCAAACTGGACACATGGAAACACACAAATTAAAAACTGGTCTGATCGATGGGATGTCACGCAGAATT | This study (modified from [12]) |
| *amyE* back | Homology arm | ATCCGTTTAGGCTGGGCGGTGATAGCTTCTCGTTCAGGCAGTACGCCTCTTTTCTTTTCCAGACCTGAGGGAGGCGGAAATGGTGTGAGGTTCCCGGGGAAAAGCCAAATAGGCGATCGCGGGAGTGCTTTATTTGAAGATCAGGCTATCACTGCGGTCAATAGATTTCACAATGTGATGGCTGGACAGCCTGAGGAACTCTCGAACCCGAATGGAAACAACCAGATATTTATGAATCAGCGCGGCTCACATGGCGTTGTGCTGGCAAATGCAGGTTCATCCTCTGTCTCTATCAATACGGCAACAAAATTGCCTGATGGCAGGTATGACAATAAAGCTGGAGCGGGTTCATTTCAAGTGAACGATGGTAAACTGACAGGCACGATCAATGCCAGGTCTGTAGCTGTGCTTTATCCTGATGATATTGCAAAAGCGCCTCATGTTTTCCTTGAGAATTACAAAACAGGTGTAACACATTCTTTCAATGATCAACTGACGATTACCTTGCGTGCAGATGCGAATACAACAAAAGCCGTTTATCAAATCAATAATGGACCAGAGACGGCGTTTAAGGATGGAGATCAATTCACAATCGGAAAAGGAGATCCATTTGGCAAAACATACACCATCATGTTAAAAGGAA | This study (modified from [12]) |

| | | |
|---|---|---|
| | | CGAACAGTGATGGTGTAACGAGGACCGAGAAATACAGTTT<br>TGTTAAAAGAGATCCAGCGTCGGCCAAAACCATCGGCTAT<br>CAAAATCCGAATCATTGGAGCCAGGTAAATGCTTATATCTA<br>TAAACATGATGGGAGCCGAGTAATTGAATTGACCGGATCT<br>TGGCCTGGAAAACCAATGACTAAAAATGCAGACGGAATTT<br>ACACGCTGACGCTGCCTGCGGACACGGATACAACCAACG<br>CAAAAGTGATTTTTAATAATGGCAGCGCCCAAGTGCCCGG<br>TCAGAATCAGCCTGGCTTTGATTACGTGCTAAATGGTTTAT<br>ATAATGACTCGGGCTTAAGCGGTTCTCTTCCCCATTGA |

*For each synthetic promoter designed in this work, the DNA binding site of the LuxR-type regulator (bolded text), –35 and –10 sequences (solid underlined text), –16 sequence (text with dashed underline), and transcription start region (TSR) sequence (italicized text) are indicated.

**Table S2. Hill parameters of sensor response functions**

| Ligand | Sensor promoter | $y_{min}$ | $y_{max}$ | $K$ | $n$ | Induction time |
|---|---|---|---|---|---|---|
| pC-HSL | $P_{Rpa\text{-}0002}$ | 0.025 | 5.77 | 0.0040 | 0.78 | 21 h |
| 3OC6-HSL | $P_{Lux\text{-}0002}$ | 0.041 | 0.60 | 0.48 | 1.37 | 21 h |
| C4-HSL | $P_{Rhl\text{-}LL12}$ | 0.080 | 3.22 | 602.01 | 0.74 | 21 h |
| 3OHC14-HSL | $P_{CinD\text{-}0102}$ | 0.14 | 2.83 | 2.32 | 1.48 | 21 h |
| pC-HSL | $P_{Rpa\text{-}0002}$ | 0.023 | 5.01 | 0.0041 | 0.80 | 5 h |
| 3OC6-HSL | $P_{Lux\text{-}0002}$ | 0.021 | 0.49 | 0.28 | 1.60 | 5 h |
| C4-HSL | $P_{Rhl\text{-}LL12}$ | 0.054 | 1.92 | 230.18 | 0.66 | 5 h |
| 3OHC14-HSL | $P_{CinD\text{-}0102}$ | 0.075 | 2.90 | 2.65 | 1.19 | 5 h |
| xylose | $P_{Xyl}$ | 0.054 | 2.39 | 16.2 | 0.95 | 21 h |

**Table S3. Combinatorial $P_{Rpa}$ promoter library design of experiments matrix**

| Combination | Promoter | –35 | –16 | –10 | TSR |
|---|---|---|---|---|---|
| 1 | $P_{Rpa-0001}$ | 0 | 0 | 0 | 1 |
| 2* | $P_{Rpa-0002}$ | 0 | 0 | 0 | 2 |
| 3* | $P_{Rpa-0010}$ | 0 | 0 | 1 | 0 |
| 4* | $P_{Rpa-0012}$ | 0 | 0 | 1 | 2 |
| 5 | $P_{Rpa-0021}$ | 0 | 0 | 2 | 1 |
| 6 | $P_{Rpa-0023}$ | 0 | 0 | 2 | 3 |
| 7 | $P_{Rpa-0100}$ | 0 | 1 | 0 | 0 |
| 8 | $P_{Rpa-0101}$ | 0 | 1 | 0 | 1 |
| 9 | $P_{Rpa-0111}$ | 0 | 1 | 1 | 1 |
| 10 | $P_{Rpa-0113}$ | 0 | 1 | 1 | 3 |
| 11 | $P_{Rpa-0122}$ | 0 | 1 | 2 | 2 |
| 12 | $P_{Rpa-0123}$ | 0 | 1 | 2 | 3 |
| 13 | $P_{Rpa-0200}$ | 0 | 2 | 0 | 0 |
| 14 | $P_{Rpa-0203}$ | 0 | 2 | 0 | 3 |
| 15 | $P_{Rpa-0211}$ | 0 | 2 | 1 | 1 |
| 16 | $P_{Rpa-0212}$ | 0 | 2 | 1 | 2 |
| 17 | $P_{Rpa-0220}$ | 0 | 2 | 2 | 0 |
| 18 | $P_{Rpa-0221}$ | 0 | 2 | 2 | 1 |
| 19* | $P_{Rpa-1000}$ | 1 | 0 | 0 | 0 |
| 20 | $P_{Rpa-1003}$ | 1 | 0 | 0 | 3 |
| 21 | $P_{Rpa-1011}$ | 1 | 0 | 1 | 1 |
| 22 | $P_{Rpa-1013}$ | 1 | 0 | 1 | 3 |
| 23 | $P_{Rpa-1020}$ | 1 | 0 | 2 | 0 |
| 24 | $P_{Rpa-1022}$ | 1 | 0 | 2 | 2 |
| 25 | $P_{Rpa-1102}$ | 1 | 1 | 0 | 2 |
| 26 | $P_{Rpa-1103}$ | 1 | 1 | 0 | 3 |
| 27 | $P_{Rpa-1110}$ | 1 | 1 | 1 | 0 |
| 28* | $P_{Rpa-1112}$ | 1 | 1 | 1 | 2 |
| 29 | $P_{Rpa-1120}$ | 1 | 1 | 2 | 0 |
| 30 | $P_{Rpa-1121}$ | 1 | 1 | 2 | 1 |
| 31 | $P_{Rpa-1201}$ | 1 | 2 | 0 | 1 |
| 32 | $P_{Rpa-1202}$ | 1 | 2 | 0 | 2 |
| 33 | $P_{Rpa-1210}$ | 1 | 2 | 1 | 0 |
| 34* | $P_{Rpa-1213}$ | 1 | 2 | 1 | 3 |
| 35 | $P_{Rpa-1222}$ | 1 | 2 | 2 | 2 |
| 36 | $P_{Rpa-1223}$ | 1 | 2 | 2 | 3 |

* Promoter sequence also contained in the original set of combinatorial $P_{Rpa}$ promoters screened.

DNA sequences for each promoter region represented by numbers (identifiers) can be found in Figure 2. The full DNA sequence for each promoter is listed in Table S1.

### Table S4. Estimated parameters of the full linear regression model of $\log_{10}$(uninduced output)

| Term | Estimate | Std Error | t Ratio | Prob>\|t\| |
|---|---|---|---|---|
| Intercept | -1.19546 | 0.033139 | -36.07 | <.0001 |
| –35[0] | -0.58934 | 0.035636 | -16.54 | <.0001 |
| –35[1] | 0.589343 | 0.035636 | 16.54 | <.0001 |
| –16[0] | -0.37297 | 0.047086 | -7.92 | <.0001 |
| –16[1] | 0.30735 | 0.047086 | 6.53 | <.0001 |
| –16[2] | 0.065617 | 0.047086 | 1.39 | 0.1675 |
| –10[0] | 0.245403 | 0.047086 | 5.21 | <.0001 |
| –10[1] | 0.848742 | 0.047086 | 18.03 | <.0001 |
| –10[2] | -1.09414 | 0.047086 | -23.24 | <.0001 |
| TSR[0] | -0.56987 | 0.059482 | -9.58 | <.0001 |
| TSR[1] | 0.787289 | 0.058979 | 13.35 | <.0001 |
| TSR[2] | 0.369952 | 0.059482 | 6.22 | <.0001 |
| TSR[3] | -0.58737 | 0.059482 | -9.87 | <.0001 |
| –35[0] X –16[0] | 0.034269 | 0.069749 | 0.49 | 0.6246 |
| –35[0] X –16[1] | -0.13993 | 0.069749 | -2.01 | 0.0484 |
| –35[0] X –16[2] | 0.10566 | 0.069749 | 1.51 | 0.134 |
| –35[1] X –16[0] | -0.03427 | 0.069749 | -0.49 | 0.6246 |
| –35[1] X –16[1] | 0.139929 | 0.069749 | 2.01 | 0.0484 |
| –35[1] X –16[2] | -0.10566 | 0.069749 | -1.51 | 0.134 |
| –35[0] X –10[0] | -0.32237 | 0.069749 | -4.62 | <.0001 |
| –35[0] X –10[1] | -0.16118 | 0.069749 | -2.31 | 0.0236 |
| –35[0] X –10[2] | 0.483552 | 0.069749 | 6.93 | <.0001 |
| –35[1] X –10[0] | 0.322372 | 0.069749 | 4.62 | <.0001 |
| –35[1] X –10[1] | 0.16118 | 0.069749 | 2.31 | 0.0236 |
| –35[1] X –10[2] | -0.48355 | 0.069749 | -6.93 | <.0001 |
| –35[0] X TSR[0] | 0.036961 | 0.078831 | 0.47 | 0.6405 |
| –35[0] X TSR[1] | -0.24973 | 0.073935 | -3.38 | 0.0012 |
| –35[0] X TSR[2] | 0.20016 | 0.078831 | 2.54 | 0.0132 |
| –35[0] X TSR[3] | 0.012611 | 0.078831 | 0.16 | 0.8733 |
| –35[1] X TSR[0] | -0.03696 | 0.078831 | -0.47 | 0.6405 |
| –35[1] X TSR[1] | 0.249732 | 0.073935 | 3.38 | 0.0012 |
| –35[1] X TSR[2] | -0.20016 | 0.078831 | -2.54 | 0.0132 |
| –35[1] X TSR[3] | -0.01261 | 0.078831 | -0.16 | 0.8733 |
| –16[0] X –10[0] | -0.30334 | 0.074047 | -4.1 | 0.0001 |
| –16[0] X –10[1] | -0.01209 | 0.079327 | -0.15 | 0.8793 |
| –16[0] X –10[2] | 0.315424 | 0.074047 | 4.26 | <.0001 |
| –16[1] X –10[0] | 0.088878 | 0.074047 | 1.2 | 0.2338 |
| –16[1] X –10[1] | -0.00065 | 0.074047 | -0.01 | 0.993 |
| –16[1] X –10[2] | -0.08823 | 0.079327 | -1.11 | 0.2696 |
| –16[2] X –10[0] | 0.214461 | 0.079327 | 2.7 | 0.0085 |
| –16[2] X –10[1] | 0.012734 | 0.074047 | 0.17 | 0.8639 |

| | | | | |
|---|---|---|---|---|
| –16[2] X –10[2] | -0.2272 | 0.074047 | -3.07 | 0.003 |
| –16[0] X TSR[0] | 0.048989 | 0.103166 | 0.47 | 0.6363 |
| –16[0] X TSR[1] | -0.05613 | 0.09822 | -0.57 | 0.5694 |
| –16[0] X TSR[2] | -0.00931 | 0.094941 | -0.1 | 0.9221 |
| –16[0] X TSR[3] | 0.016456 | 0.090081 | 0.18 | 0.8555 |
| –16[1] X TSR[0] | -0.24417 | 0.090081 | -2.71 | 0.0083 |
| –16[1] X TSR[1] | 0.267949 | 0.09822 | 2.73 | 0.0079 |
| –16[1] X TSR[2] | 0.088476 | 0.103166 | 0.86 | 0.3938 |
| –16[1] X TSR[3] | -0.11225 | 0.094941 | -1.18 | 0.2407 |
| –16[2] X TSR[0] | 0.195181 | 0.094941 | 2.06 | 0.0432 |
| –16[2] X TSR[1] | -0.21182 | 0.09822 | -2.16 | 0.0342 |
| –16[2] X TSR[2] | -0.07916 | 0.090081 | -0.88 | 0.3823 |
| –16[2] X TSR[3] | 0.095799 | 0.103166 | 0.93 | 0.356 |
| –10[0] X TSR[0] | -0.5572 | 0.094941 | -5.87 | <.0001 |
| –10[0] X TSR[1] | 0.228524 | 0.09822 | 2.33 | 0.0227 |
| –10[0] X TSR[2] | 0.413119 | 0.090081 | 4.59 | <.0001 |
| –10[0] X TSR[3] | -0.08444 | 0.103166 | -0.82 | 0.4156 |
| –10[1] X TSR[0] | 0.1251 | 0.103166 | 1.21 | 0.229 |
| –10[1] X TSR[1] | 0.138392 | 0.09822 | 1.41 | 0.1629 |
| –10[1] X TSR[2] | -0.00198 | 0.094941 | -0.02 | 0.9834 |
| –10[1] X TSR[3] | -0.26151 | 0.090081 | -2.9 | 0.0048 |
| –10[2] X TSR[0] | 0.432099 | 0.090081 | 4.8 | <.0001 |
| –10[2] X TSR[1] | -0.36692 | 0.09822 | -3.74 | 0.0004 |
| –10[2] X TSR[2] | -0.41114 | 0.103166 | -3.99 | 0.0002 |
| –10[2] X TSR[3] | 0.345957 | 0.094941 | 3.64 | 0.0005 |

**Table S5. Estimated parameters of the full linear regression model of $log_{10}$(induced output)**

| Term | Estimate | Std Error | t Ratio | Prob>\|t\| |
|---|---|---|---|---|
| Intercept | -0.65145 | 0.031381 | -20.76 | <.0001 |
| –35[0] | -0.22195 | 0.033746 | -6.58 | <.0001 |
| –35[1] | 0.221952 | 0.033746 | 6.58 | <.0001 |
| –16[0] | -0.07829 | 0.044589 | -1.76 | 0.0832 |
| –16[1] | 0.31797 | 0.044589 | 7.13 | <.0001 |
| –16[2] | -0.23968 | 0.044589 | -5.38 | <.0001 |
| –10[0] | 0.504152 | 0.044589 | 11.31 | <.0001 |
| –10[1] | 0.972992 | 0.044589 | 21.82 | <.0001 |
| –10[2] | -1.47714 | 0.044589 | -33.13 | <.0001 |
| TSR[0] | -0.53131 | 0.056327 | -9.43 | <.0001 |
| TSR[1] | 0.912343 | 0.055851 | 16.34 | <.0001 |
| TSR[2] | 0.398035 | 0.056327 | 7.07 | <.0001 |
| TSR[3] | -0.77906 | 0.056327 | -13.83 | <.0001 |
| –35[0] X –16[0] | 0.188961 | 0.066049 | 2.86 | 0.0055 |
| –35[0] X –16[1] | 0.126291 | 0.066049 | 1.91 | 0.0596 |
| –35[0] X –16[2] | -0.31525 | 0.066049 | -4.77 | <.0001 |
| –35[1] X –16[0] | -0.18896 | 0.066049 | -2.86 | 0.0055 |
| –35[1] X –16[1] | -0.12629 | 0.066049 | -1.91 | 0.0596 |
| –35[1] X –16[2] | 0.315251 | 0.066049 | 4.77 | <.0001 |
| –35[0] X –10[0] | -0.01367 | 0.066049 | -0.21 | 0.8366 |
| –35[0] X –10[1] | -0.13221 | 0.066049 | -2 | 0.0489 |
| –35[0] X –10[2] | 0.145874 | 0.066049 | 2.21 | 0.0302 |
| –35[1] X –10[0] | 0.013667 | 0.066049 | 0.21 | 0.8366 |
| –35[1] X –10[1] | 0.132207 | 0.066049 | 2 | 0.0489 |
| –35[1] X –10[2] | -0.14587 | 0.066049 | -2.21 | 0.0302 |
| –35[0] X TSR[0] | 0.142528 | 0.07465 | 1.91 | 0.06 |
| –35[0] X TSR[1] | -0.27797 | 0.070013 | -3.97 | 0.0002 |
| –35[0] X TSR[2] | 0.12204 | 0.07465 | 1.63 | 0.1062 |
| –35[0] X TSR[3] | 0.013398 | 0.07465 | 0.18 | 0.858 |
| –35[1] X TSR[0] | -0.14253 | 0.07465 | -1.91 | 0.06 |
| –35[1] X TSR[1] | 0.277967 | 0.070013 | 3.97 | 0.0002 |
| –35[1] X TSR[2] | -0.12204 | 0.07465 | -1.63 | 0.1062 |
| –35[1] X TSR[3] | -0.0134 | 0.07465 | -0.18 | 0.858 |
| –16[0] X –10[0] | 0.119436 | 0.07012 | 1.7 | 0.0926 |
| –16[0] X –10[1] | 0.080355 | 0.07512 | 1.07 | 0.2881 |
| –16[0] X –10[2] | -0.19979 | 0.07012 | -2.85 | 0.0056 |
| –16[1] X –10[0] | 0.176922 | 0.07012 | 2.52 | 0.0137 |
| –16[1] X –10[1] | -0.10261 | 0.07012 | -1.46 | 0.1475 |
| –16[1] X –10[2] | -0.07431 | 0.07512 | -0.99 | 0.3257 |
| –16[2] X –10[0] | -0.29636 | 0.07512 | -3.95 | 0.0002 |
| –16[2] X –10[1] | 0.022254 | 0.07012 | 0.32 | 0.7518 |

| | | | | |
|---|---|---|---|---|
| –16[2] X –10[2] | 0.274104 | 0.07012 | 3.91 | 0.0002 |
| –16[0] X TSR[0] | 0.226082 | 0.097695 | 2.31 | 0.0234 |
| –16[0] X TSR[1] | -0.37302 | 0.093011 | -4.01 | 0.0001 |
| –16[0] X TSR[2] | 0.06919 | 0.089906 | 0.77 | 0.4439 |
| –16[0] X TSR[3] | 0.077752 | 0.085304 | 0.91 | 0.3649 |
| –16[1] X TSR[0] | -0.07048 | 0.085304 | -0.83 | 0.4113 |
| –16[1] X TSR[1] | 0.171456 | 0.093011 | 1.84 | 0.0692 |
| –16[1] X TSR[2] | -0.10805 | 0.097695 | -1.11 | 0.2722 |
| –16[1] X TSR[3] | 0.007072 | 0.089906 | 0.08 | 0.9375 |
| –16[2] X TSR[0] | -0.1556 | 0.089906 | -1.73 | 0.0876 |
| –16[2] X TSR[1] | 0.201568 | 0.093011 | 2.17 | 0.0334 |
| –16[2] X TSR[2] | 0.038861 | 0.085304 | 0.46 | 0.65 |
| –16[2] X TSR[3] | -0.08482 | 0.097695 | -0.87 | 0.388 |
| –10[0] X TSR[0] | -0.65169 | 0.089906 | -7.25 | <.0001 |
| –10[0] X TSR[1] | 0.056491 | 0.093011 | 0.61 | 0.5454 |
| –10[0] X TSR[2] | 0.542309 | 0.085304 | 6.36 | <.0001 |
| –10[0] X TSR[3] | 0.052894 | 0.097695 | 0.54 | 0.5898 |
| –10[1] X TSR[0] | 0.158586 | 0.097695 | 1.62 | 0.1087 |
| –10[1] X TSR[1] | -0.10562 | 0.093011 | -1.14 | 0.2597 |
| –10[1] X TSR[2] | 0.057748 | 0.089906 | 0.64 | 0.5226 |
| –10[1] X TSR[3] | -0.11071 | 0.085304 | -1.3 | 0.1983 |
| –10[2] X TSR[0] | 0.493108 | 0.085304 | 5.78 | <.0001 |
| –10[2] X TSR[1] | 0.049132 | 0.093011 | 0.53 | 0.5989 |
| –10[2] X TSR[2] | -0.60006 | 0.097695 | -6.14 | <.0001 |
| –10[2] X TSR[3] | 0.057817 | 0.089906 | 0.64 | 0.5221 |

**Table S6. Estimated parameters of the linear regression model of $log_{10}$(uninduced output) without interaction**

| Term | Estimate | Std Error | t Ratio | Prob>\|t\| |
|---|---|---|---|---|
| Intercept | -1.2232 | 0.062325 | -19.63 | <.0001 |
| –35[0] | -0.59099 | 0.063512 | -9.31 | <.0001 |
| –35[1] | 0.590994 | 0.063512 | 9.31 | <.0001 |
| –16[0] | -0.35348 | 0.08814 | -4.01 | 0.0001 |
| –16[1] | 0.295578 | 0.08814 | 3.35 | 0.0011 |
| –16[2] | 0.057903 | 0.08814 | 0.66 | 0.5127 |
| –10[0] | 0.237689 | 0.08814 | 2.7 | 0.0082 |
| –10[1] | 0.868228 | 0.08814 | 9.85 | <.0001 |
| –10[2] | -1.10592 | 0.08814 | -12.55 | <.0001 |
| TSR[0] | -0.59457 | 0.10818 | -5.5 | <.0001 |
| TSR[1] | 0.732344 | 0.110006 | 6.66 | <.0001 |
| TSR[2] | 0.347074 | 0.10818 | 3.21 | 0.0018 |
| TSR[3] | -0.48484 | 0.10818 | -4.48 | <.0001 |

**Table S7. Estimated parameters of the linear regression model of $\log_{10}$(induced output) without interaction**

| Term | Estimate | Std Error | t Ratio | Prob>\|t\| |
|---|---|---|---|---|
| Intercept | -0.68234 | 0.054986 | -12.41 | <.0001 |
| –35[0] | -0.25033 | 0.056033 | -4.47 | <.0001 |
| –35[1] | 0.250333 | 0.056033 | 4.47 | <.0001 |
| –16[0] | -0.07339 | 0.077762 | -0.94 | 0.3476 |
| –16[1] | 0.304761 | 0.077762 | 3.92 | 0.0002 |
| –16[2] | -0.23137 | 0.077762 | -2.98 | 0.0037 |
| –10[0] | 0.512464 | 0.077762 | 6.59 | <.0001 |
| –10[1] | 0.977889 | 0.077762 | 12.58 | <.0001 |
| –10[2] | -1.49035 | 0.077762 | -19.17 | <.0001 |
| TSR[0] | -0.59251 | 0.095441 | -6.21 | <.0001 |
| TSR[1] | 0.860032 | 0.097052 | 8.86 | <.0001 |
| TSR[2] | 0.424818 | 0.095441 | 4.45 | <.0001 |
| TSR[3] | -0.69234 | 0.095441 | -7.25 | <.0001 |

**Table S8. Estimated parameters of the reduced linear regression model of $\log_{10}$(uninduced output)**

| Term | Estimate | Std Error | t Ratio | Prob>\|t\| |
|---|---|---|---|---|
| Intercept | -1.2232 | 0.040891 | -29.91 | <.0001 |
| –35[0] | -0.58474 | 0.043371 | -13.48 | <.0001 |
| –35[1] | 0.584742 | 0.043371 | 13.48 | <.0001 |
| –16[0] | -0.35348 | 0.057828 | -6.11 | <.0001 |
| –16[1] | 0.295578 | 0.057828 | 5.11 | <.0001 |
| –16[2] | 0.057903 | 0.057828 | 1 | 0.3193 |
| –10[0] | 0.237689 | 0.057828 | 4.11 | <.0001 |
| –10[1] | 0.868228 | 0.057828 | 15.01 | <.0001 |
| –10[2] | -1.10592 | 0.057828 | -19.12 | <.0001 |
| TSR[0] | -0.53549 | 0.072285 | -7.41 | <.0001 |
| TSR[1] | 0.73026 | 0.072285 | 10.1 | <.0001 |
| TSR[2] | 0.391528 | 0.072285 | 5.42 | <.0001 |
| TSR[3] | -0.58629 | 0.072285 | -8.11 | <.0001 |
| –35[0] X –10[0] | -0.26273 | 0.061336 | -4.28 | <.0001 |
| –35[0] X –10[1] | -0.19692 | 0.061336 | -3.21 | 0.0018 |
| –35[0] X –10[2] | 0.45965 | 0.061336 | 7.49 | <.0001 |
| –35[1] X –10[0] | 0.262735 | 0.061336 | 4.28 | <.0001 |
| –35[1] X –10[1] | 0.196915 | 0.061336 | 3.21 | 0.0018 |
| –35[1] X –10[2] | -0.45965 | 0.061336 | -7.49 | <.0001 |
| –10[0] X TSR[0] | -0.58806 | 0.102227 | -5.75 | <.0001 |
| –10[0] X TSR[1] | 0.145919 | 0.102227 | 1.43 | 0.1569 |
| –10[0] X TSR[2] | 0.404544 | 0.102227 | 3.96 | 0.0002 |
| –10[0] X TSR[3] | 0.037593 | 0.102227 | 0.37 | 0.7139 |
| –10[1] X TSR[0] | 0.09913 | 0.102227 | 0.97 | 0.3348 |
| –10[1] X TSR[1] | 0.107972 | 0.102227 | 1.06 | 0.2937 |
| –10[1] X TSR[2] | 0.155092 | 0.102227 | 1.52 | 0.1327 |
| –10[1] X TSR[3] | -0.36219 | 0.102227 | -3.54 | 0.0006 |
| –10[2] X TSR[0] | 0.488926 | 0.102227 | 4.78 | <.0001 |
| –10[2] X TSR[1] | -0.25389 | 0.102227 | -2.48 | 0.0148 |
| –10[2] X TSR[2] | -0.55964 | 0.102227 | -5.47 | <.0001 |
| –10[2] X TSR[3] | 0.324601 | 0.102227 | 3.18 | 0.002 |

**Table S9. Estimated parameters of the reduced linear regression model of $log_{10}$(induced output)**

| Term | Estimate | Std Error | t Ratio | Prob>\|t\| |
|---|---|---|---|---|
| Intercept | -0.68234 | 0.041724 | -16.35 | <.0001 |
| –35[0] | -0.22691 | 0.044255 | -5.13 | <.0001 |
| –35[1] | 0.226911 | 0.044255 | 5.13 | <.0001 |
| –16[0] | -0.07339 | 0.059006 | -1.24 | 0.2167 |
| –16[1] | 0.304761 | 0.059006 | 5.16 | <.0001 |
| –16[2] | -0.23137 | 0.059006 | -3.92 | 0.0002 |
| –10[0] | 0.512464 | 0.059006 | 8.68 | <.0001 |
| –10[1] | 0.977889 | 0.059006 | 16.57 | <.0001 |
| –10[2] | -1.49035 | 0.059006 | -25.26 | <.0001 |
| TSR[0] | -0.58991 | 0.072435 | -8.14 | <.0001 |
| TSR[1] | 0.852225 | 0.073758 | 11.55 | <.0001 |
| TSR[2] | 0.427421 | 0.072435 | 5.9 | <.0001 |
| TSR[3] | -0.68974 | 0.072435 | -9.52 | <.0001 |
| –10[0] X TSR[0] | -0.65189 | 0.104077 | -6.26 | <.0001 |
| –10[0] X TSR[1] | 0.253791 | 0.102202 | 2.48 | 0.0148 |
| –10[0] X TSR[2] | 0.623692 | 0.102674 | 6.07 | <.0001 |
| –10[0] X TSR[3] | -0.22559 | 0.102674 | -2.2 | 0.0305 |
| –10[1] X TSR[0] | 0.364049 | 0.102674 | 3.55 | 0.0006 |
| –10[1] X TSR[1] | -0.28056 | 0.102202 | -2.75 | 0.0073 |
| –10[1] X TSR[2] | -0.03158 | 0.104077 | -0.3 | 0.7622 |
| –10[1] X TSR[3] | -0.05191 | 0.102674 | -0.51 | 0.6144 |
| –10[2] X TSR[0] | 0.287843 | 0.102674 | 2.8 | 0.0062 |
| –10[2] X TSR[1] | 0.026772 | 0.102202 | 0.26 | 0.7939 |
| –10[2] X TSR[2] | -0.59211 | 0.102674 | -5.77 | <.0001 |
| –10[2] X TSR[3] | 0.277497 | 0.104077 | 2.67 | 0.009 |

### Table S10. Pairwise comparison of categorical promoter regions

| Response | Promoter element | Pairs | Pairwise differences | | | | Significance |
|---|---|---|---|---|---|---|---|
| | | | Difference | Std Err Difference | 99% Confidence Interval of the difference | | |
| | | | | | Lower | Upper | |
| $Log_{10}$ (uninduced output) | –35 region | 1 – 0 | 1.169484 | 0.086742 | 0.94127 | 1.397698 | <.0001 |
| | –16 region | 1 – 0 | 0.649058 | 0.100162 | 0.385539 | 0.912577 | <.0001 |
| | | 2 – 0 | 0.411383 | 0.100162 | 0.147865 | 0.674902 | <.0001 |
| | | 1 – 2 | 0.237675 | 0.100162 | -0.02584 | 0.501194 | 0.0198 |
| | –10 region | 1 – 2 | 1.974146 | 0.100162 | 1.710627 | 2.237664 | <.0001 |
| | | 0 – 2 | 1.343606 | 0.100162 | 1.080087 | 1.607125 | <.0001 |
| | | 1 – 0 | 0.63054 | 0.100162 | 0.367021 | 0.894059 | <.0001 |
| | TSR | 1 – 3 | 1.316554 | 0.118041 | 1.005994 | 1.627114 | <.0001 |
| | | 1 – 0 | 1.265753 | 0.118041 | 0.955193 | 1.576313 | <.0001 |
| | | 2 – 3 | 0.977822 | 0.118041 | 0.667262 | 1.288382 | <.0001 |
| | | 2 – 0 | 0.927021 | 0.118041 | 0.616461 | 1.237581 | <.0001 |
| | | 1 – 2 | 0.338732 | 0.118041 | 0.028172 | 0.649292 | 0.0051 |
| | | 0 – 3 | 0.050801 | 0.118041 | -0.25976 | 0.361361 | 0.6679 |
| $Log_{10}$ (induced output) | –35 region | 1–0 | 0.453822 | 0.088509 | 0.221066 | 0.686578 | <.0001 |
| | –16 region | 1–2 | 0.53613 | 0.102202 | 0.267367 | 0.804893 | <.0001 |
| | | 1–0 | 0.378152 | 0.102202 | 0.109389 | 0.646916 | 0.0004 |
| | | 0–2 | 0.157978 | 0.102202 | -0.11079 | 0.426741 | 0.1256 |
| | –10 region | 1–2 | 2.468243 | 0.102202 | 2.199479 | 2.737006 | <.0001 |
| | | 0–2 | 2.002818 | 0.102202 | 1.734055 | 2.271581 | <.0001 |
| | | 1–0 | 0.465425 | 0.102202 | 0.196662 | 0.734188 | <.0001 |
| | TSR | 1–3 | 1.541962 | 0.11964 | 1.22734 | 1.856584 | <.0001 |
| | | 1–0 | 1.442135 | 0.11964 | 1.127513 | 1.756756 | <.0001 |

| | | | | | | | |
|---|---|---|---|---|---|---|---|
| | | 2–3 | 1.117157 | 0.118012 | 0.806816 | 1.427498 | <.0001 |
| | | 2–0 | 1.01733 | 0.118012 | 0.706989 | 1.327671 | <.0001 |
| | | 1–2 | 0.424804 | 0.11964 | 0.110183 | 0.739426 | 0.0006 |
| | | 0–3 | 0.099827 | 0.118012 | -0.21051 | 0.410168 | 0.3998 |

*The comparison was done by Student's *t* test (two-tailed).

### Table S11. Plasmids constructs used in this work

| Plasmid Name | Description |
|---|---|
| pMZ1000 | A synthetic pC-HSL sensor on integration plasmid (pNH4). $P_{Rpa-0000}$ drives the expression of GFP. A constitutive promoter $P_{FtsH}$ and a synthetic RpaR rbs1 drive the expression of RpaR. |
| pMZ1001 | A synthetic pC-HSL sensor on integration plasmid (pNH4). $P_{Rpa-0002}$ drives the expression of GFP. A constitutive promoter $P_{FtsH}$ and a synthetic RpaR rbs1 drive the expression of RpaR. |
| pMZ1002 | A synthetic pC-HSL sensor on integration plasmid (pNH4). $P_{Rpa-0011}$ drives the expression of GFP. A constitutive promoter $P_{FtsH}$ and a synthetic RpaR rbs1 drive the expression of RpaR. |
| pMZ1003 | A synthetic pC-HSL sensor on integration plasmid (pNH4). $P_{Rpa-0012}$ drives the expression of GFP. A constitutive promoter $P_{FtsH}$ and a synthetic RpaR rbs1 drive the expression of RpaR. |
| pMZ1004 | A synthetic pC-HSL sensor on integration plasmid (pNH4). $P_{Rpa-0102}$ drives the expression of GFP. A constitutive promoter $P_{FtsH}$ and a synthetic RpaR rbs1 drive the expression of RpaR. |
| pMZ1005 | A synthetic pC-HSL sensor on integration plasmid (pNH4). $P_{Rpa-0112}$ drives the expression of GFP. A constitutive promoter $P_{FtsH}$ and a synthetic RpaR rbs1 drive the expression of RpaR. |
| pMZ1006 | A synthetic pC-HSL sensor on integration plasmid (pNH4). $P_{Rpa-0202}$ drives the expression of GFP. A constitutive promoter $P_{FtsH}$ and a synthetic RpaR rbs1 drive the expression of RpaR. |
| pMZ1007 | A synthetic pC-HSL sensor on integration plasmid (pNH4). $P_{Rpa-0233}$ drives the expression of GFP. A constitutive promoter $P_{FtsH}$ and a synthetic RpaR rbs1 drive the expression of RpaR. |
| pMZ1008 | A synthetic pC-HSL sensor on integration plasmid (pNH4). $P_{Rpa-1000}$ drives the expression of GFP. A constitutive promoter $P_{FtsH}$ and a synthetic RpaR rbs1 drive the expression of RpaR. |
| pMZ1009 | A synthetic pC-HSL sensor on integration plasmid (pNH4). $P_{Rpa-1002}$ drives the expression of GFP. A constitutive promoter $P_{FtsH}$ and a synthetic RpaR rbs1 drive the expression of RpaR. |
| pMZ1010 | A synthetic pC-HSL sensor on integration plasmid (pNH4). $P_{Rpa-1012}$ drives the expression of GFP. A constitutive promoter $P_{FtsH}$ and a synthetic RpaR rbs1 drive the expression of RpaR. |
| pMZ1011 | A synthetic pC-HSL sensor on integration plasmid (pNH4). $P_{Rpa-1112}$ drives the expression of GFP. A constitutive promoter $P_{FtsH}$ and a synthetic RpaR rbs1 drive the expression of RpaR. |
| pMZ1012 | A synthetic pC-HSL sensor on integration plasmid (pNH4). $P_{Rpa-1113}$ drives the expression of GFP. A constitutive promoter $P_{FtsH}$ and a synthetic RpaR rbs1 drive the expression of RpaR. |
| pMZ1013 | A synthetic pC-HSL sensor on integration plasmid (pNH4). $P_{Rpa-1213}$ drives the expression of GFP. A constitutive promoter $P_{FtsH}$ and a synthetic RpaR rbs1 drive the expression of RpaR. |
| pMZ1014 | A synthetic pC-HSL sensor on integration plasmid (pNH4). $P_{Rpa-0001}$ drives the expression of GFP. A constitutive promoter $P_{FtsH}$ and a synthetic RpaR rbs1 drive the expression of RpaR. |
| pMZ1015 | A synthetic pC-HSL sensor on integration plasmid (pNH4). $P_{Rpa-0010}$ drives the expression of GFP. A constitutive promoter $P_{FtsH}$ and a synthetic RpaR rbs1 drive the expression of RpaR. |
| pMZ1016 | A synthetic pC-HSL sensor on integration plasmid (pNH4). $P_{Rpa-0021}$ drives the expression of GFP. A constitutive promoter $P_{FtsH}$ and a synthetic RpaR rbs1 drive the expression of RpaR. |
| pMZ1017 | A synthetic pC-HSL sensor on integration plasmid (pNH4). $P_{Rpa-0023}$ drives the expression of GFP. A constitutive promoter $P_{FtsH}$ and a synthetic RpaR rbs1 drive the expression of RpaR. |
| pMZ1018 | A synthetic pC-HSL sensor on integration plasmid (pNH4). $P_{Rpa-0100}$ drives the expression of GFP. A constitutive promoter $P_{FtsH}$ and a synthetic RpaR rbs1 drive the expression of RpaR. |
| pMZ1019 | A synthetic pC-HSL sensor on integration plasmid (pNH4). $P_{Rpa-0101}$ drives the expression of GFP. A constitutive promoter $P_{FtsH}$ and a synthetic RpaR rbs1 drive the expression of RpaR. |
| pMZ1020 | A synthetic pC-HSL sensor on integration plasmid (pNH4). $P_{Rpa-0111}$ drives the expression of GFP. A constitutive promoter $P_{FtsH}$ and a synthetic RpaR rbs1 drive the expression of RpaR. |
| pMZ1021 | A synthetic pC-HSL sensor on integration plasmid (pNH4). $P_{Rpa-0113}$ drives the expression of GFP. A constitutive promoter $P_{FtsH}$ and a synthetic RpaR rbs1 drive the expression of RpaR. |
| pMZ1022 | A synthetic pC-HSL sensor on integration plasmid (pNH4). $P_{Rpa-0122}$ drives the expression of GFP. A constitutive promoter $P_{FtsH}$ and a synthetic RpaR rbs1 drive the expression of RpaR. |
| pMZ1023 | A synthetic pC-HSL sensor on integration plasmid (pNH4). $P_{Rpa-0123}$ drives the expression of GFP. A constitutive promoter $P_{FtsH}$ and a synthetic RpaR rbs1 drive the expression of RpaR. |
| pMZ1024 | A synthetic pC-HSL sensor on integration plasmid (pNH4). $P_{Rpa-0200}$ drives the expression of GFP. A constitutive promoter $P_{FtsH}$ and a synthetic RpaR rbs1 drive the expression of RpaR. |
| pMZ1025 | A synthetic pC-HSL sensor on integration plasmid (pNH4). $P_{Rpa-0203}$ drives the expression of GFP. A constitutive promoter $P_{FtsH}$ and a synthetic RpaR rbs1 drive the expression of RpaR. |
| pMZ1026 | A synthetic pC-HSL sensor on integration plasmid (pNH4). $P_{Rpa-0211}$ drives the expression of GFP. A constitutive promoter $P_{FtsH}$ and a synthetic RpaR rbs1 drive the expression of RpaR. |
| pMZ1027 | A synthetic pC-HSL sensor on integration plasmid (pNH4). $P_{Rpa-0212}$ drives the expression of GFP. A constitutive promoter $P_{FtsH}$ and a synthetic RpaR rbs1 drive the expression of RpaR. |

| | |
|---|---|
| pMZ1028 | A synthetic pC-HSL sensor on integration plasmid (pNH4). $P_{Rpa\text{-}0220}$ drives the expression of GFP. A constitutive promoter $P_{FtsH}$ and a synthetic RpaR rbs1 drive the expression of RpaR. |
| pMZ1029 | A synthetic pC-HSL sensor on integration plasmid (pNH4). $P_{Rpa\text{-}0221}$ drives the expression of GFP. A constitutive promoter $P_{FtsH}$ and a synthetic RpaR rbs1 drive the expression of RpaR. |
| pMZ1030 | A synthetic pC-HSL sensor on integration plasmid (pNH4). $P_{Rpa\text{-}1003}$ drives the expression of GFP. A constitutive promoter $P_{FtsH}$ and a synthetic RpaR rbs1 drive the expression of RpaR. |
| pMZ1031 | A synthetic pC-HSL sensor on integration plasmid (pNH4). $P_{Rpa\text{-}1011}$ drives the expression of GFP. A constitutive promoter $P_{FtsH}$ and a synthetic RpaR rbs1 drive the expression of RpaR. |
| pMZ1032 | A synthetic pC-HSL sensor on integration plasmid (pNH4). $P_{Rpa\text{-}1013}$ drives the expression of GFP. A constitutive promoter $P_{FtsH}$ and a synthetic RpaR rbs1 drive the expression of RpaR. |
| pMZ1033 | A synthetic pC-HSL sensor on integration plasmid (pNH4). $P_{Rpa\text{-}1020}$ drives the expression of GFP. A constitutive promoter $P_{FtsH}$ and a synthetic RpaR rbs1 drive the expression of RpaR. |
| pMZ1034 | A synthetic pC-HSL sensor on integration plasmid (pNH4). $P_{Rpa\text{-}1022}$ drives the expression of GFP. A constitutive promoter $P_{FtsH}$ and a synthetic RpaR rbs1 drive the expression of RpaR. |
| pMZ1035 | A synthetic pC-HSL sensor on integration plasmid (pNH4). $P_{Rpa\text{-}1102}$ drives the expression of GFP. A constitutive promoter $P_{FtsH}$ and a synthetic RpaR rbs1 drive the expression of RpaR. |
| pMZ1036 | A synthetic pC-HSL sensor on integration plasmid (pNH4). $P_{Rpa\text{-}1103}$ drives the expression of GFP. A constitutive promoter $P_{FtsH}$ and a synthetic RpaR rbs1 drive the expression of RpaR. |
| pMZ1037 | A synthetic pC-HSL sensor on integration plasmid (pNH4). $P_{Rpa\text{-}1110}$ drives the expression of GFP. A constitutive promoter $P_{FtsH}$ and a synthetic RpaR rbs1 drive the expression of RpaR. |
| pMZ1038 | A synthetic pC-HSL sensor on integration plasmid (pNH4). $P_{Rpa\text{-}1120}$ drives the expression of GFP. A constitutive promoter $P_{FtsH}$ and a synthetic RpaR rbs1 drive the expression of RpaR. |
| pMZ1039 | A synthetic pC-HSL sensor on integration plasmid (pNH4). $P_{Rpa\text{-}1121}$ drives the expression of GFP. A constitutive promoter $P_{FtsH}$ and a synthetic RpaR rbs1 drive the expression of RpaR. |
| pMZ1040 | A synthetic pC-HSL sensor on integration plasmid (pNH4). $P_{Rpa\text{-}1201}$ drives the expression of GFP. A constitutive promoter $P_{FtsH}$ and a synthetic RpaR rbs1 drive the expression of RpaR. |
| pMZ1041 | A synthetic pC-HSL sensor on integration plasmid (pNH4). $P_{Rpa\text{-}1202}$ drives the expression of GFP. A constitutive promoter $P_{FtsH}$ and a synthetic RpaR rbs1 drive the expression of RpaR. |
| pMZ1042 | A synthetic pC-HSL sensor on integration plasmid (pNH4). $P_{Rpa\text{-}1210}$ drives the expression of GFP. A constitutive promoter $P_{FtsH}$ and a synthetic RpaR rbs1 drive the expression of RpaR. |
| pMZ1043 | A synthetic pC-HSL sensor on integration plasmid (pNH4). $P_{Rpa\text{-}1222}$ drives the expression of GFP. A constitutive promoter $P_{FtsH}$ and a synthetic RpaR rbs1 drive the expression of RpaR. |
| pMZ1044 | A synthetic pC-HSL sensor on integration plasmid (pNH4). $P_{Rpa\text{-}1223}$ drives the expression of GFP. A constitutive promoter $P_{FtsH}$ and a synthetic RpaR rbs1 drive the expression of RpaR. |
| pMZ2000 | A synthetic 3OC6-HSL sensor on integration plasmid (pNH4). $P_{Lux\text{-}Ec}$ (or $P_{Lux\text{-}XXXX}$) drives the expression of GFP. A constitutive promoter $P_{FtsH}$ and a synthetic LuxR rbs1 drive the expression of LuxR. |
| pMZ2001 | A synthetic 3OC6-HSL sensor on integration plasmid (pNH4). $P_{Lux\text{-}0002}$ drives the expression of GFP. A constitutive promoter $P_{FtsH}$ and a synthetic LuxR rbs1 drive the expression of LuxR. |
| pMZ2002 | A synthetic 3OC6-HSL sensor on integration plasmid (pNH4). $P_{Lux\text{-}0011}$ drives the expression of GFP. A constitutive promoter $P_{FtsH}$ and a synthetic LuxR rbs1 drive the expression of LuxR. |
| pMZ2003 | A synthetic 3OC6-HSL sensor on integration plasmid (pNH4). $P_{Lux\text{-}0012}$ drives the expression of GFP. A constitutive promoter $P_{FtsH}$ and a synthetic LuxR rbs1 drive the expression of LuxR. |
| pMZ2004 | A synthetic 3OC6-HSL sensor on integration plasmid (pNH4). $P_{Lux\text{-}0102}$ drives the expression of GFP. A constitutive promoter $P_{FtsH}$ and a synthetic LuxR rbs1 drive the expression of LuxR. |
| pMZ2005 | A synthetic 3OC6-HSL sensor on integration plasmid (pNH4). $P_{Lux\text{-}0W02}$ drives the expression of GFP. A constitutive promoter $P_{FtsH}$ and a synthetic LuxR rbs1 drive the expression of LuxR. |
| pMZ2006 | A synthetic 3OC6-HSL sensor on integration plasmid (pNH4). $P_{Lux\text{-}0S02}$ drives the expression of GFP. A constitutive promoter $P_{FtsH}$ and a synthetic LuxR rbs1 drive the expression of LuxR. |
| pMZ2007 | A synthetic 3OC6-HSL sensor on integration plasmid (pNH4). $P_{Lux\text{-}0002}$ drives the expression of GFP. A constitutive promoter $P_{FtsH}$ and a synthetic LuxR rbs2 drive the expression of LuxR. |
| pMZ2008 | A synthetic 3OC6-HSL sensor on integration plasmid (pNH4). $P_{Lux\text{-}0011}$ drives the expression of GFP. A constitutive promoter $P_{FtsH}$ and a synthetic LuxR rbs2 drive the expression of LuxR. |
| pMZ2009 | A synthetic 3OC6-HSL sensor on integration plasmid (pNH4). $P_{Lux\text{-}0012}$ drives the expression of GFP. A constitutive promoter $P_{FtsH}$ and a synthetic LuxR rbs2 drive the expression of LuxR. |
| pMZ2010 | A synthetic 3OC6-HSL sensor on integration plasmid (pNH4). $P_{Lux\text{-}0102}$ drives the expression of GFP. A constitutive promoter $P_{FtsH}$ and a synthetic LuxR rbs2 drive the expression of LuxR. |
| pMZ2011 | A synthetic 3OC6-HSL sensor on integration plasmid (pNH4). $P_{Lux\text{-}0W02}$ drives the expression of GFP. A constitutive promoter $P_{FtsH}$ and a synthetic LuxR rbs2 drive the expression of LuxR. |
| pMZ2012 | A synthetic 3OC6-HSL sensor on integration plasmid (pNH4). $P_{Lux\text{-}0S02}$ drives the expression of GFP. A constitutive promoter $P_{FtsH}$ and a synthetic LuxR rbs2 drive the expression of LuxR. |

| | |
|---|---|
| pMZ2013 | A synthetic 3OC6-HSL sensor on integration plasmid (pNH4). $P_{Lux-1042}$ drives the expression of GFP. A constitutive promoter $P_{FtsH}$ and a synthetic LuxR rbs2 drive the expression of LuxR. |
| pMZ2014 | A synthetic 3OC6-HSL sensor on integration plasmid (pNH4). $P_{Lux-1052}$ drives the expression of GFP. A constitutive promoter $P_{FtsH}$ and a synthetic LuxR rbs2 drive the expression of LuxR. |
| pMZ2015 | A synthetic 3OC6-HSL sensor on integration plasmid (pNH4). $P_{Lux-1062}$ drives the expression of GFP. A constitutive promoter $P_{FtsH}$ and a synthetic LuxR rbs2 drive the expression of LuxR. |
| pMZ2016 | A synthetic 3OC6-HSL sensor on integration plasmid (pNH4). $P_{Lux-2032}$ drives the expression of GFP. A constitutive promoter $P_{FtsH}$ and a synthetic LuxR rbs2 drive the expression of LuxR. |
| pMZ2017 | A synthetic 3OC6-HSL sensor on integration plasmid (pNH4). $P_{Lux-3002}$ drives the expression of GFP. A constitutive promoter $P_{FtsH}$ and a synthetic LuxR rbs2 drive the expression of LuxR. |
| pMZ2018 | A synthetic 3OC6-HSL sensor on integration plasmid (pNH4). $P_{Lux-4012}$ drives the expression of GFP. A constitutive promoter $P_{FtsH}$ and a synthetic LuxR rbs2 drive the expression of LuxR. |
| pMZ2019 | A synthetic 3OC6-HSL sensor on integration plasmid (pNH4). $P_{Lux-0002}$ drives the expression of GFP. An inducible promoter $P_{Xyl}$ and a synthetic LuxR rbs2 drive the expression of LuxR. |
| pMZ3000 | A synthetic C4-HSL sensor on integration plasmid (pNH4). $P_{Rhl-Ec}$ (or $P_{Rhl-LLLL}$) drives the expression of GFP. A constitutive promoter $P_{FtsH}$ and a synthetic RhlR rbs1 drive the expression of RhlR. |
| pMZ3001 | A synthetic C4-HSL sensor on integration plasmid (pNH4). $P_{Rhl-0002}$ drives the expression of GFP. A constitutive promoter $P_{FtsH}$ and a synthetic RhlR rbs1 drive the expression of RhlR. |
| pMZ3002 | A synthetic C4-HSL sensor on integration plasmid (pNH4). $P_{Rhl-0011}$ drives the expression of GFP. A constitutive promoter $P_{FtsH}$ and a synthetic RhlR rbs1 drive the expression of RhlR. |
| pMZ3003 | A synthetic C4-HSL sensor on integration plasmid (pNH4). $P_{Rhl-0012}$ drives the expression of GFP. A constitutive promoter $P_{FtsH}$ and a synthetic RhlR rbs1 drive the expression of RhlR. |
| pMZ3004 | A synthetic C4-HSL sensor on integration plasmid (pNH4). $P_{Rhl-0102}$ drives the expression of GFP. A constitutive promoter $P_{FtsH}$ and a synthetic RhlR rbs1 drive the expression of RhlR. |
| pMZ3005 | A synthetic C4-HSL sensor on integration plasmid (pNH4). $P_{Rhl-0012}$ drives the expression of GFP. A constitutive promoter $P_{FtsH}$ and a synthetic RhlR rbs2 drive the expression of RhlR. |
| pMZ3006 | A synthetic C4-HSL sensor on integration plasmid (pNH4). $P_{Rhl-0102}$ drives the expression of GFP. A constitutive promoter $P_{FtsH}$ and a synthetic RhlR rbs2 drive the expression of RhlR. |
| pMZ3007 | A synthetic C4-HSL sensor on integration plasmid (pNH4). $P_{Rhl-0002}$ drives the expression of GFP. A constitutive promoter $P_{FtsH}$ and a synthetic RhlR rbs3 drive the expression of RhlR. |
| pMZ3008 | A synthetic C4-HSL sensor on integration plasmid (pNH4). $P_{Rhl-0011}$ drives the expression of GFP. A constitutive promoter $P_{FtsH}$ and a synthetic RhlR rbs3 drive the expression of RhlR. |
| pMZ3009 | A synthetic C4-HSL sensor on integration plasmid (pNH4). $P_{Rhl-0012}$ drives the expression of GFP. A constitutive promoter $P_{FtsH}$ and a synthetic RhlR rbs3 drive the expression of RhlR. |
| pMZ3010 | A synthetic C4-HSL sensor on integration plasmid (pNH4). $P_{Rhl-0102}$ drives the expression of GFP. A constitutive promoter $P_{FtsH}$ and a synthetic RhlR rbs3 drive the expression of RhlR. |
| pMZ3011 | A synthetic C4-HSL sensor on integration plasmid (pNH4). $P_{Rhl-LLL2}$ drives the expression of GFP. A constitutive promoter $P_{FtsH}$ and a synthetic RhlR rbs3 drive the expression of RhlR. |
| pMZ3012 | A synthetic C4-HSL sensor on integration plasmid (pNH4). $P_{Rhl-LL12}$ drives the expression of GFP. A constitutive promoter $P_{FtsH}$ and a synthetic RhlR rbs3 drive the expression of RhlR. |
| pMZ3013 | A synthetic C4-HSL sensor on integration plasmid (pNH4). $P_{Rhl-L1L2L}$ drives the expression of GFP. A constitutive promoter $P_{FtsH}$ and a synthetic RhlR rbs3 drive the expression of RhlR. |
| pMZ3014 | A synthetic C4-HSL sensor on integration plasmid (pNH4). $P_{Rhl-LL12}$ drives the expression of GFP. An inducible promoter $P_{xyl}$ and a synthetic RhlR rbs3 drive the expression of RhlR. |
| pMZ4000 | A synthetic 3OHC14-HSL sensor on integration plasmid (pNH4). $P_{Cin-Ec}$ (or $P_{Cin-NNNN}$) drives the expression of GFP. A constitutive promoter $P_{FtsH}$ and a synthetic CinR rbs1 drive the expression of CinR. |
| pMZ4001 | A synthetic 3OHC14-HSL sensor on integration plasmid (pNH4). $P_{CinD-0002}$ containing CinR binding box D drives the expression of GFP. A constitutive promoter $P_{FtsH}$ and a synthetic CinR rbs1 drive the expression of CinR. |
| pMZ4002 | A synthetic 3OHC14-HSL sensor on integration plasmid (pNH4). $P_{CinD-0011}$ containing CinR binding box D drives the expression of GFP. A constitutive promoter $P_{FtsH}$ and a synthetic CinR rbs1 drive the expression of CinR. |
| pMZ4003 | A synthetic 3OHC14-HSL sensor on integration plasmid (pNH4). $P_{CinD-0012}$ containing CinR binding box D drives the expression of GFP. A constitutive promoter $P_{FtsH}$ and a synthetic CinR rbs1 drive the expression of CinR. |
| pMZ4004 | A synthetic 3OHC14-HSL sensor on integration plasmid (pNH4). $P_{CinD-0102}$ containing CinR binding box D drives the expression of GFP. A constitutive promoter $P_{FtsH}$ and a synthetic CinR rbs1 drive the expression of CinR. |

| | |
|---|---|
| pMZ4005 | A synthetic 3OHC14-HSL sensor on integration plasmid (pNH4). $P_{CinA\text{-}0002}$ containing CinR binding box A drives the expression of GFP. A constitutive promoter $P_{FtsH}$ and a synthetic CinR rbs1 drive the expression of CinR. |
| pMZ4006 | A synthetic 3OHC14-HSL sensor on integration plasmid (pNH4). $P_{CinA\text{-}0011}$ containing CinR binding box A drives the expression of GFP. A constitutive promoter $P_{FtsH}$ and a synthetic CinR rbs1 drive the expression of CinR. |
| pMZ4007 | A synthetic 3OHC14-HSL sensor on integration plasmid (pNH4). $P_{CinA\text{-}0012}$ containing CinR binding box A drives the expression of GFP. A constitutive promoter $P_{FtsH}$ and a synthetic CinR rbs1 drive the expression of CinR. |
| pMZ4008 | A synthetic 3OHC14-HSL sensor on integration plasmid (pNH4). $P_{CinA\text{-}0102}$ containing CinR binding box A drives the expression of GFP. A constitutive promoter $P_{FtsH}$ and a synthetic CinR rbs1 drive the expression of CinR. |
| pMZ4009 | A synthetic 3OHC14-HSL sensor on integration plasmid (pNH4). $P_{CinB\text{-}0002}$ containing CinR binding box B drives the expression of GFP. A constitutive promoter $P_{FtsH}$ and a synthetic CinR rbs1 drive the expression of CinR. |
| pMZ4010 | A synthetic 3OHC14-HSL sensor on integration plasmid (pNH4). $P_{CinB\text{-}0011}$ containing CinR binding box B drives the expression of GFP. A constitutive promoter $P_{FtsH}$ and a synthetic CinR rbs1 drive the expression of CinR. |
| pMZ4011 | A synthetic 3OHC14-HSL sensor on integration plasmid (pNH4). $P_{CinB\text{-}0012}$ containing CinR binding box B drives the expression of GFP. A constitutive promoter $P_{FtsH}$ and a synthetic CinR rbs1 drive the expression of CinR. |
| pMZ4012 | A synthetic 3OHC14-HSL sensor on integration plasmid (pNH4). $P_{CinB\text{-}0102}$ containing CinR binding box B drives the expression of GFP. A constitutive promoter $P_{FtsH}$ and a synthetic CinR rbs1 drive the expression of CinR. |
| pMZ4013 | A synthetic 3OHC14-HSL sensor on integration plasmid (pNH4). $P_{CinC\text{-}0002}$ containing CinR binding box C drives the expression of GFP. A constitutive promoter $P_{FtsH}$ and a synthetic CinR rbs1 drive the expression of CinR. |
| pMZ4014 | A synthetic 3OHC14-HSL sensor on integration plasmid (pNH4). $P_{CinC\text{-}0011}$ containing CinR binding box C drives the expression of GFP. A constitutive promoter $P_{FtsH}$ and a synthetic CinR rbs1 drive the expression of CinR. |
| pMZ4015 | A synthetic 3OHC14-HSL sensor on integration plasmid (pNH4). $P_{CinC\text{-}0012}$ containing CinR binding box C drives the expression of GFP. A constitutive promoter $P_{FtsH}$ and a synthetic CinR rbs1 drive the expression of CinR. |
| pMZ4016 | A synthetic 3OHC14-HSL sensor on integration plasmid (pNH4). $P_{CinC\text{-}0102}$ containing CinR binding box C drives the expression of GFP. A constitutive promoter $P_{FtsH}$ and a synthetic CinR rbs1 drive the expression of CinR. |
| pMZ4017 | A synthetic 3OHC14-HSL sensor on integration plasmid (pNH4). $P_{CinD\text{-}0102}$ containing CinR binding box D drives the expression of GFP. An inducible promoter $P_{Xyl}$ and a synthetic CinR rbs1 drive the expression of CinR. |
| pMZ5000 | A synthetic 3OC12-HSL sensor on integration plasmid (pNH4). $P_{Las\text{-}Ec}$ (or $P_{Las\text{-}ssss}$) drives the expression of GFP. A constitutive promoter $P_{FtsH}$ and a synthetic LasR rbs2 drive the expression of LasR. |
| pMZ5001 | A synthetic 3OC12-HSL sensor on integration plasmid (pNH4). $P_{Las\text{-}0002}$ drives the expression of GFP. A constitutive promoter $P_{FtsH}$ and a synthetic LasR rbs1 drive the expression of LasR. |
| pMZ5002 | A synthetic 3OC12-HSL sensor on integration plasmid (pNH4). $P_{Las\text{-}0011}$ drives the expression of GFP. A constitutive promoter $P_{FtsH}$ and a synthetic LasR rbs1 drive the expression of LasR. |
| pMZ5003 | A synthetic 3OC12-HSL sensor on integration plasmid (pNH4). $P_{Las\text{-}0012}$ drives the expression of GFP. A constitutive promoter $P_{FtsH}$ and a synthetic LasR rbs1 drive the expression of LasR. |
| pMZ5004 | A synthetic 3OC12-HSL sensor on integration plasmid (pNH4). $P_{Las\text{-}0102}$ drives the expression of GFP. A constitutive promoter $P_{FtsH}$ and a synthetic LasR rbs1 drive the expression of LasR. |
| pMZ5005 | A synthetic 3OC12-HSL sensor on integration plasmid (pNH4). $P_{Las\text{-}0002}$ drives the expression of GFP. A constitutive promoter $P_{FtsH}$ and a synthetic LasR rbs2 drive the expression of LasR. |
| pMZ5006 | A synthetic 3OC12-HSL sensor on integration plasmid (pNH4). $P_{Las\text{-}0011}$ drives the expression of GFP. A constitutive promoter $P_{FtsH}$ and a synthetic LasR rbs2 drive the expression of LasR. |
| pMZ5007 | A synthetic 3OC12-HSL sensor on integration plasmid (pNH4). $P_{Las\text{-}0012}$ drives the expression of GFP. A constitutive promoter $P_{FtsH}$ and a synthetic LasR rbs2 drive the expression of LasR. |
| pMZ5008 | A synthetic 3OC12-HSL sensor on integration plasmid (pNH4). $P_{Las\text{-}0102}$ drives the expression of GFP. A constitutive promoter $P_{FtsH}$ and a synthetic LasR rbs2 drive the expression of LasR. |
| pMZ5009 | A synthetic 3OC12-HSL sensor on integration plasmid (pNH4). $P_{Las\text{-}0002}$ drives the expression of GFP. A constitutive promoter $P_{FtsH}$ and a synthetic LasR rbs3 drive the expression of LasR. |

| | |
|---|---|
| pMZ5010 | A synthetic 3OC12-HSL sensor on integration plasmid (pNH4). $P_{Las\text{-}0011}$ drives the expression of GFP. A constitutive promoter $P_{FtsH}$ and a synthetic LasR rbs3 drive the expression of LasR. |
| pMZ5011 | A synthetic 3OC12-HSL sensor on integration plasmid (pNH4). $P_{Las\text{-}0012}$ drives the expression of GFP. A constitutive promoter $P_{FtsH}$ and a synthetic LasR rbs3 drive the expression of LasR. |
| pMZ5012 | A synthetic 3OC12-HSL sensor on integration plasmid (pNH4). $P_{Las\text{-}0102}$ drives the expression of GFP. A constitutive promoter $P_{FtsH}$ and a synthetic LasR rbs3 drive the expression of LasR. |
| pMZ5013 | A synthetic 3OC12-HSL sensor on integration plasmid (pNH4). $P_{Las\text{-}SSS2}$ drives the expression of GFP. A constitutive promoter $P_{FtsH}$ and a synthetic LasR rbs3 drive the expression of LasR. |
| pMZ5014 | A synthetic 3OC12-HSL sensor on integration plasmid (pNH4). $P_{Las\text{-}SS12}$ drives the expression of GFP. A constitutive promoter $P_{FtsH}$ and a synthetic LasR rbs3 drive the expression of LasR. |
| pMZ5015 | A synthetic 3OC12-HSL sensor on integration plasmid (pNH4). $P_{Las\text{-}S1S2}$ drives the expression of GFP. A constitutive promoter $P_{FtsH}$ and a synthetic LasR rbs3 drive the expression of LasR. |
| pMZ5016 | A synthetic 3OC12-HSL sensor on integration plasmid (pNH4). $P_{Las\text{-}0012}$ drives the expression of GFP. An inducible promoter Pxyl and a synthetic LasR rbs3 drive the expression of LasR mutant (LasR L125W–A127T). |

*All plasmids were constructed on integration plasmids for chromosomal integration of the described construct in *B. subtilis* at the *amyE* locus for subsequent sensor assaying. The corresponding *B. subtilis* strains for each integration plasmid construct were named as MZXXXX (using the same digits as the plasmid name).
